# Trans-Allosteric Activation Releases Distinct Conformational Traps in Kinase Heterodimers

**DOI:** 10.64898/2026.08.31.748385

**Authors:** Akira Imamoto, Yichao Wu, Ai Shinobu, Mariko Okada

## Abstract

Protein kinases function as dynamic, mechanically coupled nodes, yet the conformational drivers of multimeric activation remain unclear. Here, we present AlloQuant, a computational suite that translates AlphaFold3 structural ensembles into quantitative metrics of kinase regulation, including internal network rigidity, metastable-state populations, and sub-angstrom conformational drivers. Applying AlloQuant to CDK1, we demonstrate that binding of the Cyclin B1 (CCNB1) cofactor mechanically decouples a hyper-rigid inactive kinase core, allowing activating phosphorylation (pT161) to subsequently re-impose localized tension on the catalytic machinery. Conversely, the C-terminal Src kinase (CSK) faces a distinct conformational trap. While nucleotide-free monomeric CSK spontaneously samples a pre-active geometry, ATP binding excludes the active αC-In conformation in all but 1 of 225 models. We show that docking partner engagement overcomes this blockade. Autophosphorylation of SRC at the activation loop (Y419) redistributes SRC conformational states without altering bulk rigidity. This redistribution is structurally coupled to the conformational state of CSK via the regulatory spine, not the catalytic machinery. Rather than mechanically deforming CSK, SRC engagement acts by conformational selection, committing roughly a quarter of CSK molecules to a fully active state. Thus, trans-allosteric kinase activation operates by defining the accessible conformational landscape of the receiver kinase. That control is exerted through mechanical remodeling in cofactor-dependent complexes and through conformational selection in transient kinase-kinase heterodimers. These findings establish AlloQuant as a general framework for quantifying how a binding partner reshapes a kinase’s conformational landscape, applicable across the kinome because it assigns landmarks by profile-HMM alignment.

**AUTHOR SUMMARY:** Protein kinases act as molecular switches that regulate processes such as cell division; their dysregulation is a hallmark of cancer. Many kinases cannot activate autonomously and must instead dock with partner proteins. Such activation is hard to observe, being structural rather than chemical. Artificial-intelligence structure prediction now yields diverse conformational ensembles, not single snapshots. To analyze them we developed AlloQuant, which quantifies protein rigidity and favored conformational states. Applied to two kinase pairs, AlloQuant revealed that partner binding releases structural traps through radically different mechanisms. In the cell-division kinase CDK1, the partner loosens an overly rigid catalytic core, priming it for a chemical modification that then re-tightens the active site. In the kinase CSK, by contrast, the ATP fuel required for catalysis prevents the active shape from forming, making a docking partner essential. Partner binding does not forcibly reshape CSK; it biases the structural odds, committing a fraction of molecules to the active form. Because AlloQuant works on any ensemble of structures, the same measurements can gauge how a binding partner reshapes the conformational options of other proteins, and identifying the traps that restrict activation will improve predictions of how a partner or a targeted therapeutic can spring them.

## INTRODUCTION

Protein kinases operate as a dynamic, mechanically coupled network within the protein rather than a static bulk [1]. The transition from a dormant, inactive state to a catalytically competent conformation requires precise spatial realignment of a small, conserved set of regulatory elements. While the influence of localized post-translational modifications, such as activation loop phosphorylation, is well established, many kinases also require extensive multimeric docking to reach their functional states.

The canonical paradigm for docking-mediated activation is the Cyclin-Dependent Kinase (CDK) family. CDK1 requires physical binding of its regulatory partner, Cyclin B1 (CCNB1), alongside specific activating and inhibitory phosphorylations, to govern cell cycle progression. Cell-cycle transitions are driven by successive CDK-cyclin pairs, and genetic dissection has narrowed that set considerably. The interphase CDKs prove individually dispensable, whereas CDK1 is required and can itself drive the division cycle; for review, see [2]. The activating T161 mark is installed by CDK7-cyclin H (CAK), while the inhibitory T14 and Y15 marks are placed by Wee1 and Myt1 and removed by CDC25, so an assembled complex can be held poised as readily as released. Cyclin binding visibly forces rotation of the PSTAIRE (αC) helix [3], but the conformational shifts and internal mechanical rewiring that prime the CDK1 core for catalysis remain incompletely mapped.

Such mechanical requirements become more complex in transient kinase-kinase heterodimers that lack dedicated cofactors. Within the Src family of non-receptor tyrosine kinases (SFKs), the C-terminal Src kinase (CSK) phosphorylates SRC at a conserved C-terminal tyrosine (Y530 in humans) to lock it into an autoinhibited state [4–7]. That lock is conditional. SRC already autophosphorylated in its activation loop is still phosphorylated at Y530 by CSK yet remains active, in reconstituted systems and in oxidatively stressed cells alike [8, 9]. Two independent knockouts establish that this regulation is physiologically essential. Mice lacking Csk show constitutive Src-family activation and arrest in mid-gestation with neural defects [10, 11]. If CSK operated solely as an enzyme acting on a flexible tail, a transient catalytic encounter would suffice. Yet efficient negative regulation requires the broader SRC architecture, and phosphorylation relies on a conserved docking footprint across the CSK kinase domain [12–14]. In systems such as the ERBB receptor family, this extensive docking serves as a conduit for asymmetric trans-allosteric regulation, where a "driver" kinase mechanically forces a "receiver" kinase into an active conformation [15]. We follow that usage for the receiver, and call its partner the activator. We hypothesize that the transient CSK-SRC heterodimer is regulated by an analogous partner-dependent mechanism.

Here, we ask whether trans-allosteric activation follows a common mechanism across kinase multimeric complexes. AlphaFold3 returns ensembles rather than single structures. However, no standardized tool converts them into quantitative statements about kinase regulation. We therefore developed AlloQuant, a high-dimensional analysis suite that maps sub-angstrom structural drivers and global network rigidity across AlphaFold3-generated conformational ensembles. Evaluating the canonical CDK1-CCNB1 complex first, we define how cofactor docking systematically decouples a rigidly trapped kinase to enable catalytic scaffolding. Applying these principles to the CSK-SRC system, we find that SRC, as a substrate, undergoes a phosphorylation-gated conformational redistribution, without bulk stiffening, that concentrates the ensemble in states competent to engage CSK, and that the two kinases’ conformations are coupled within individual complexes, biasing the dormant enzyme (CSK) toward its active state. By mapping these states, we redefine multimeric kinase regulation as a coupled trans-allosteric event in which the partner determines which conformations the receiver can reach, through mechanical change in the cofactor complex and through conformational selection in the kinase-kinase pair. Because AlloQuant is built for the kinase family rather than for these two complexes, the same measurements apply to any kinase system for which a structural ensemble can be generated.

## RESULTS

### AlloQuant: a computational suite for quantifying allostery in AlphaFold3 kinase ensembles

We have designed AlloQuant as a two-module suite that turns a set of AlphaFold3 models into reproducible readouts of conformational state and internal mechanical coupling (Fig. 1).

**Figure 1.**
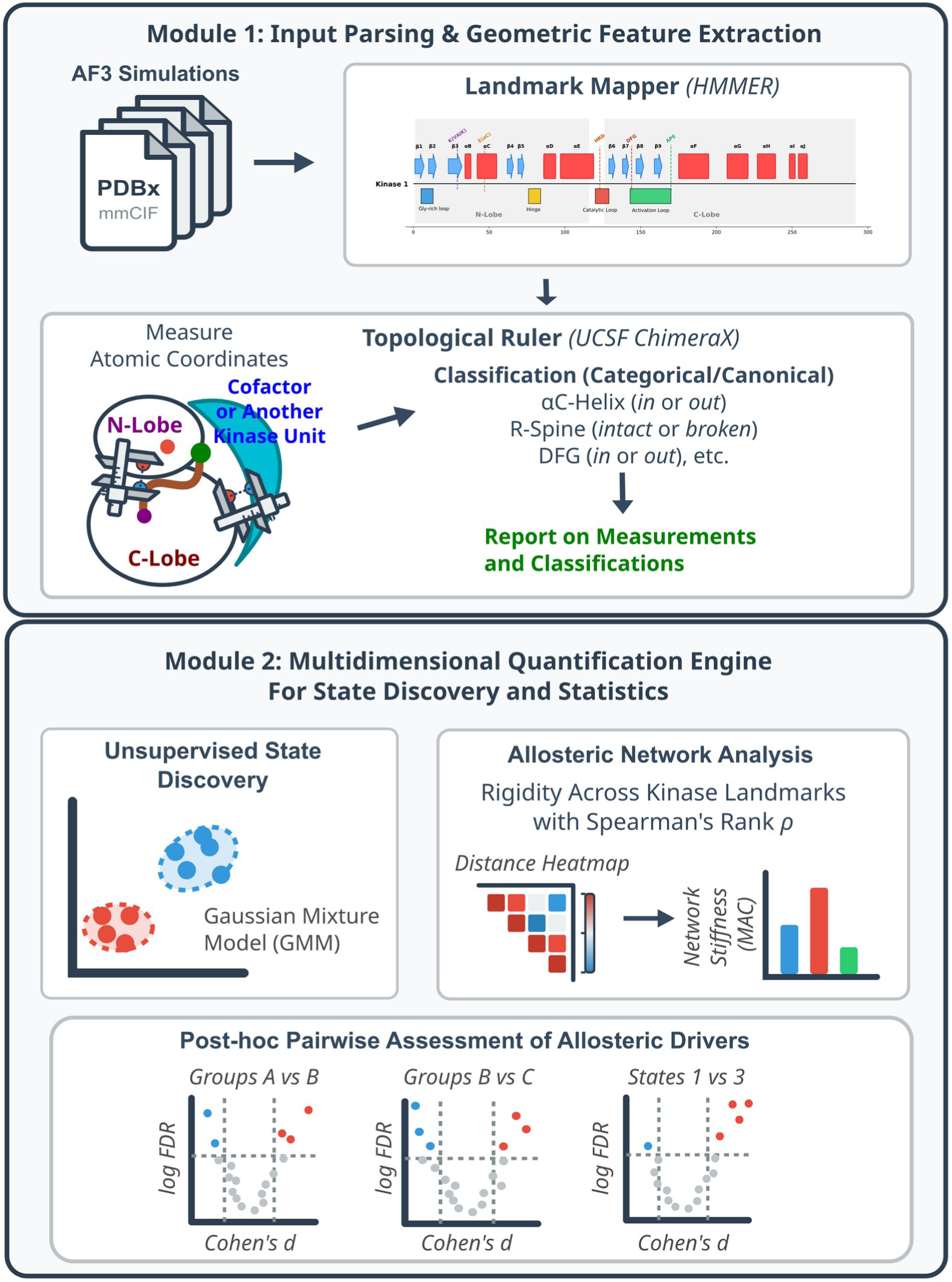
The AlloQuant suite: an interactive two-module pipeline for quantifying trans-allosteric kinase mechanics from structural ensembles. Module 1 batch-processes structures en masse into conformational ensembles based on the user’s experimental design. AlphaFold3 models (PDBx/mmCIF) are annotated by the Landmark Mapper, which uses profile hidden Markov models (HMMER) to assign canonical kinase landmarks across the N- and C-lobes: the Gly-rich loop, the β3 lysine (VAIK), the αC glutamate (E-αC), the HRD and DFG motifs, and the activation loop (APE). The Topological Ruler (UCSF ChimeraX) then measures inter-atomic distances anchored to these landmarks, including contacts to a bound cofactor or partner kinase, and assigns canonical categorical states (αC-helix in/out, regulatory spine intact/broken, DFG in/out). It reports per-model continuous measurements and classifications. Module 2 receives the Module 1 measurements to conduct explicit allosteric analysis. Unsupervised state discovery partitions the ensemble into metastable states (Gaussian Mixture Model). Allosteric network analysis quantifies internal mechanical coupling as the Mean Absolute Correlation (MAC) of the Spearman rank-correlation (ρ) matrix among structural metrics, capturing the network stiffening and decoupling that distinguishes conditions. Post-hoc pairwise assessment then isolates the micro-metric drivers separating any two groups or states, shown as volcano plots of significance (−log FDR) versus signed effect size. Both modules run interactively with the user’s choices (e.g., GMM vs. K-means for state discovery; Cohen’s *d* vs. rank-biserial effect size); this study used GMM and Cohen’s *d*.

Module 1 annotates structure. It parses monomeric or multimeric coordinates (PDBx/mmCIF) and, rather than depending on fixed residue numbering, maps the canonical kinase landmarks onto any sequence by profile-HMM alignment to the Pfam Pkinase model (HMMER), so that one pipeline applies across the kinome; it then uses headless UCSF ChimeraX to measure 23 inter-landmark distances and 8 angles per chain from which it makes 9 categorical conformational calls, and, in multimers, scores the asymmetric roles of the subunits (S1 Dataset). AlloQuant’s readouts are anchored in the structural vocabulary of the kinase fold, and each metric reports on a conserved regulatory element defined by decades of structural work. A kinase toggles between active and inactive conformations by repositioning a small, coupled set of these elements. The regulatory αC-helix rotates “in” or “out” to form or break the β3-Lys–αC-Glu salt bridge [3], and the activation-loop DFG motif flips between a catalytically competent “in” and a disengaged “out” geometry [16]. Two hydrophobic “spines” assemble only in the active state: the catalytic spine that clamps the ATP adenine and the regulatory spine that couples the αC-helix to the activation loop [17, 18], the latter gated by the αC-β4 loop [19]. We label activation-loop backbone geometry with the Dunbrack dihedral nomenclature [20]. Because these elements move as one mechanically coupled network rather than in isolation [1], any single snapshot is uninformative about regulation. AlloQuant is built to quantify how their positions and couplings are distributed across the whole ensemble structures.

Once Module 1 completes its evaluations, Module 2 then quantifies the ensemble beyond the macro-structural categories described above. It discovers metastable states without supervision (PCA then GMM/BIC), measures global network rigidity as a single scalar, the Mean Absolute Correlation (MAC; below), and ranks the sub-angstrom structural drivers that separate any two states by signed effect size (Cohen’s *d* or rank-biserial correlation; see Methods and S2 Dataset). Because every measurement is anchored to an HMMER-assigned landmark rather than an absolute residue number, one 48-column definition applies across the kinome, making the measurements comparable between kinases and the suite directly reusable on new kinase systems.

Among many structural assessments conducted in Module 2, the Mean Absolute Correlation (MAC) is one key quantity, reporting mechanical stiffness as a readout of allosteric effects. It is the average absolute Spearman correlation between all pairs of inter-landmark distances across an ensemble. A high MAC means the landmarks move together, the signature of a mechanically coupled, tension-loaded domain, and a low MAC means they move independently. We report it for whole conditions (global MAC) and for individual metastable states (intrinsic MAC), which are related but not interchangeable (see Methods for details).

As an initial validation, we tested AlloQuant Module 1 on the CSK–SRC crystal structure 3D7T [21]. AlloQuant recovered the same donor–receiver asymmetry and reproduced the reference classifier Kincore [22] to the reported precision (Note 1 in S1 Text; S1 Table). The published crystal structure is complexed with the ATP-competitive inhibitor staurosporine and the CSK variant in the PDB structure carries the K361A/K362A substitutions while part of the activation loop is unresolved. We therefore tested only landmark placements and interface topology rather than the conformational distributions. Nonetheless, it is noteworthy that the crystal structure is an essentially inactive conformation for both CSK and SRC in canonical classifications (S1 Table).

Upon the validation of Module 1 above, we then applied the suite to two model heterodimers, CDK1-CCNB1, a kinase bound by a non-kinase cofactor, and CSK-SRC, a transient kinase-kinase pair lacking any dedicated cofactor.

### Model 1: CDK1-CCNB1 (kinase-non kinase heterodimer)

#### CCNB1 Decouples the Natively Trapped, Rigid CDK1 Monomer

To resolve the mechanics of CDK1 activation, we tracked the master mitotic regulator from its inactive monomeric baseline to its fully phosphorylated complex (Figs. 2 and 3). The unphosphorylated CDK1 monomer is natively trapped in a stiff, tightly coupled network (Figs. 2A and 3A) and occupies a single deep inactive basin (State 4; Figs. 2A-2C). Despite an intact regulatory spine it displays an extruded αC-helix (77% αC-Out), an unpacked activation loop, and a non-canonical BLBminus backbone in place of the Active BLAminus signature (Fig. 2D; Note 2 in S1 Text). Across every condition the DFG motif scores spatially “in” and therefore does not discriminate along the activation path. Binding of the rigid CCNB1 cofactor (+CCNB1) reshapes this landscape. CCNB1 is rigid in the literal sense that its own fold does not change. Its median per-residue Cα RMSF is 0.12 to 0.14 Å in all four complexes, against 0.14 to 0.39 Å for CDK1 in the same models (Note 15 in S1 Text, Table 9). CCNB1 forces the αC-helix inward and packs the activation loop, establishing a near-uniform Active BLAminus architecture even before phosphorylation (Figs. 2D and 3B), as a cyclin-fold activator can do without the T-loop mark in the CDK5-p25 complex [23], and it does so by lowering global rigidity (Fig. 2A). CCNB1 thus decouples the trapped CDK1 core, lowering global MAC from 0.37 to 0.22, and the resulting ensemble no longer occupies State 4 but a more flexible baseline (*p.adj* = 4.5×10^-6^). This baseline is not mechanically uniform. The +CCNB1 condition exists in two GMM metastable states of significantly different intrinsic rigidity (State 8, 50%, intrinsic MAC 0.18; State 7, 39%, 0.26; *p.adj* = 0.009; S2 Table). Whether this reflects a dynamic stiffness fluctuation within the complex or two distinct routes out of the apo monomer remains unclear.

**Figure 2.**
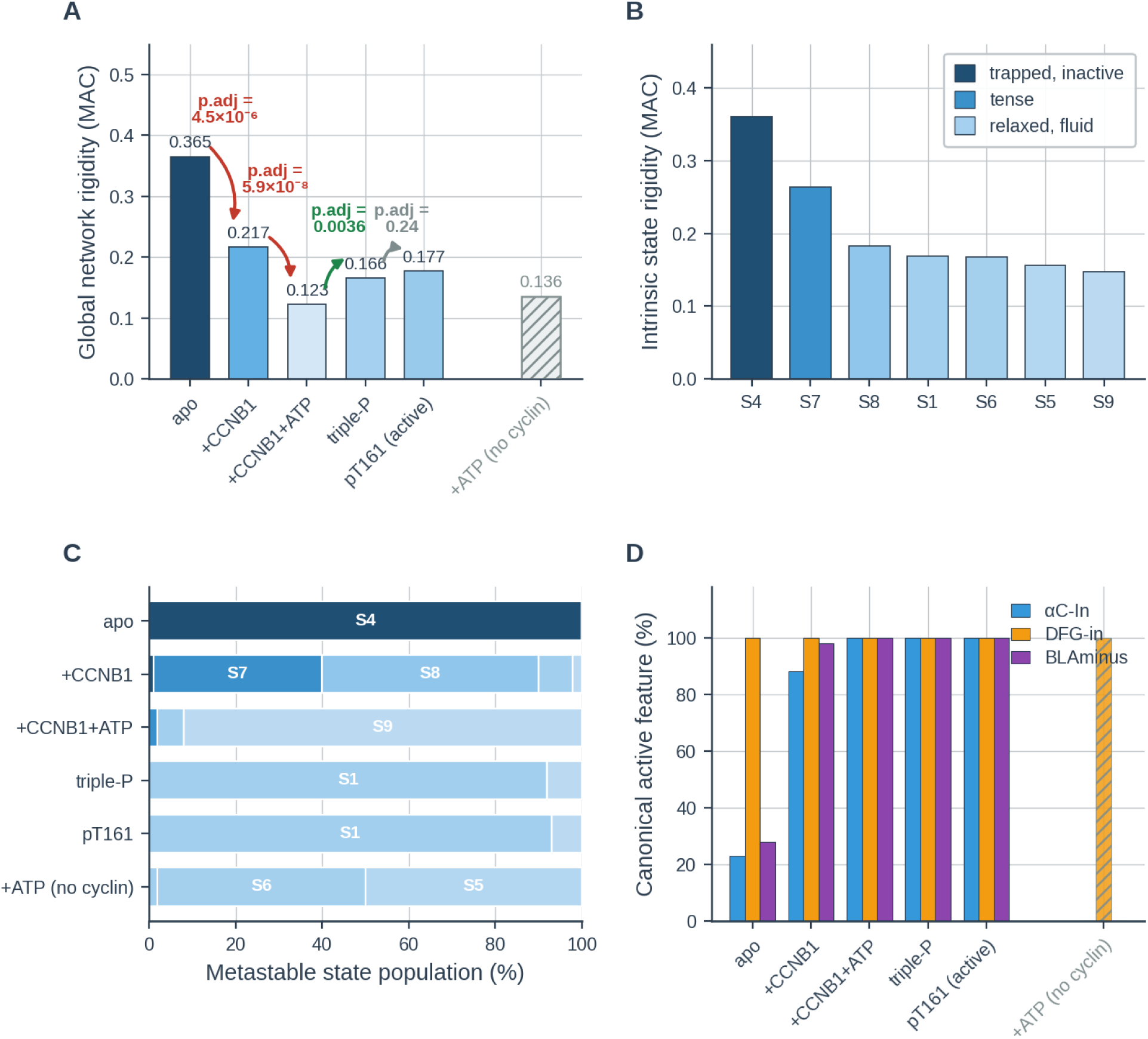
CCNB1 decouples the trapped kinase core, and pT161 re-tensions it. AlloQuant analysis of CDK1–CCNB1 conformational ensembles (*n* = 100 models per condition). **(A)** Global network rigidity (Mean Absolute Correlation, MAC) along the biological activation path: apo →+CCNB1 → +CCNB1+ATP → triple-P (pT14/pY15/pT161) → pT161 (active). Bars are colored on the rigidity ramp, and each transition is annotated with its Benjamini–Hochberg (BH) adjusted *p*-value (*p.adj*). CCNB1 and then ATP decouple the rigid apo network, and phosphorylation retensions it. The inhibited and active phospho-complexes remain indistinguishable (*p.adj* = 0.24). Recomputed on the landmark distances that two conditions share, each transition retains 96 to 97% of its amplitude, except +CCNB1 to +CCNB1+ATP at 69% (Methods; Note 3 in S1 Text; S5 Table). A cyclin-free CDK1+ATP control is hatched. **(B)** Within-state (intrinsic) MAC of the GMM metastable states, ordered most- to least-rigid and shaded on the continuous rigidity ramp. Pairwise Wilcoxon tests resolve three statistically distinct rigidity tiers (legend; S2 Table): the trapped, inactive apo state (State 4); an intermediate tense tier (State 7); and a relaxed, fluid tier (States 8, 1, 6, 5 and 9). The first two tiers each differ significantly from every other state, and the five members of the relaxed tier are mutually indistinguishable across all ten of their pairwise comparisons. **(C)** Metastable state populations (%) per condition, in the order of (A), with segments shaded by intrinsic rigidity. **(D)** Categorical active-state signatures (% of models scored αC-In, DFG-in, Active BLAminus) across the same path. DFG-in is saturated at 100% in every condition and therefore does not discriminate between them. AlphaFold3 essentially never samples a DFG-out geometry for this system. The no-cyclin control is hatched throughout.

**Figure 3.**
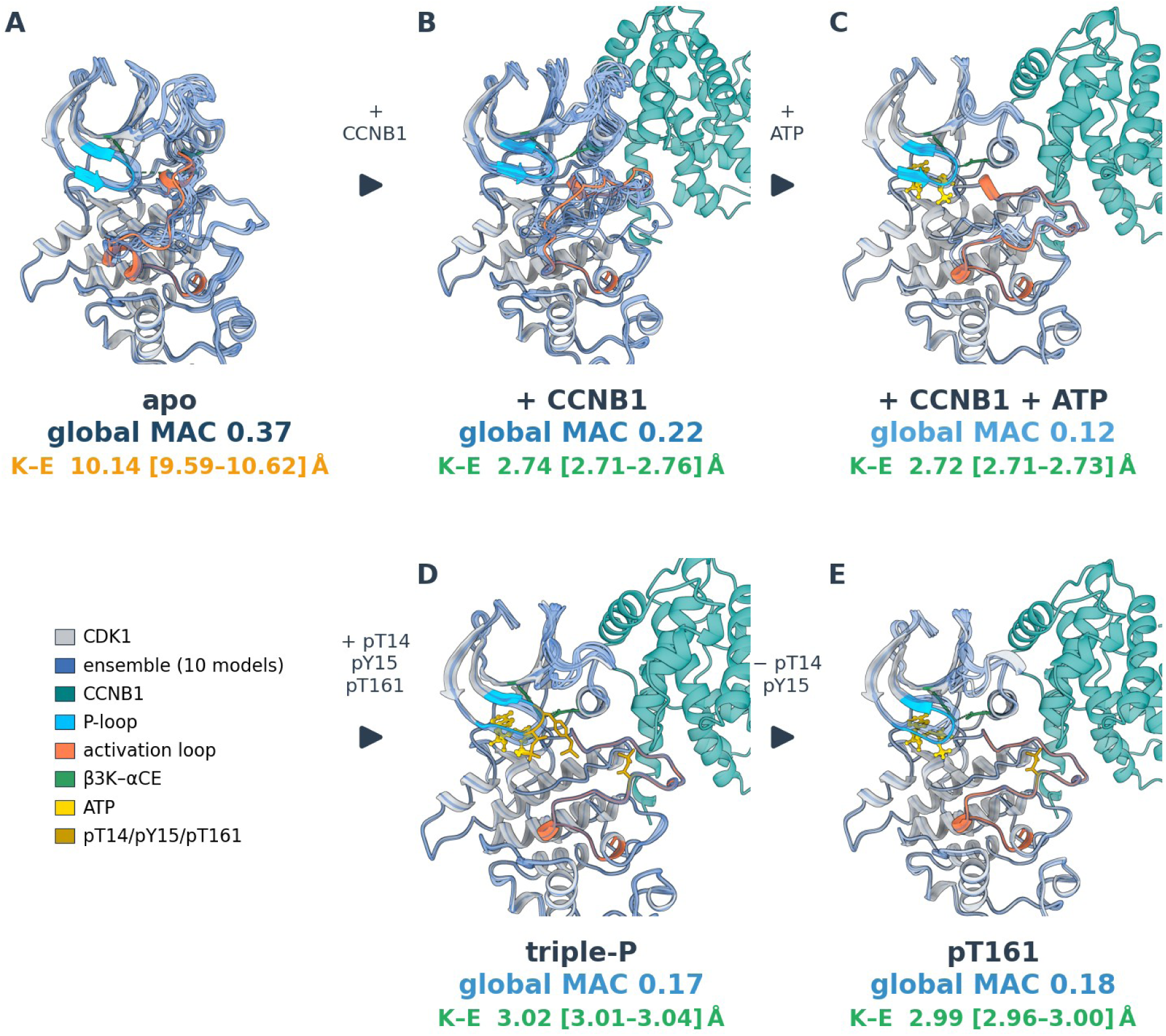
The CDK1 activation cycle as an ordered structural series. Each panel shows the CDK1 kinase domain in a common orientation, rendered as a 10-model Cα-rope ensemble (one representative per seed, sampled from the 100-model ensemble), with canonical elements colored in a centroid representative. Element colors are defined in the key at lower left. CCNB1 and the ensemble traces are semi-transparent. The global MAC (rigidity ramp) and K–E salt-bridge distance (β3-Lys–αC-Glu; median [95% bootstrap CI]) below each panel quantify the full 100-model ensemble; K–E is colored green (formed, < 4Å) or amber (open). Metastable state composition per condition is given in Fig. 2C. Panels follow the biological activation order, each transition annotated: **(A)** apo CDK1, a trapped, over-rigid monomer; **(B)** +CCNB1; **(C)** +CCNB1+ATP, the maximally decoupled intermediate; **(D)** the inhibitory triple-phosphorylated complex (pT14/pY15/pT161); and **(E)** the active pT161 complex. The three marks are installed by distinct kinases (T161 by CAK/CDK7–cyclin H; the inhibitory T14/Y15 by Wee1 and Myt1), and the T14/Y15 pair is removed by CDC25 to license mitotic activation (transition arrows). The inhibited (D) and active (E) complexes are statistically indistinguishable in global rigidity (*p.adj* = 0.24) and occupy the same metastable state at near-identical occupancy (92% versus 93%); they differ only in a localized active-site phosphate clash (S1 Fig). *n* = 100 models per condition (20 seeds × 5 samples).

#### ATP Binding Fluidizes the Heterodimer Prior to Phosphorylation

After CCNB1 association, ATP binding gives the most flexible unphosphorylated ensemble (+CCNB1+ATP; p.adj = 5.9×10^-8^, compared to +CCNB1; Fig 2A). Adding ATP decouples the global allosteric network still further, and the ensemble occupies State 9 rather than the two-state +CCNB1 intermediate. State 9 is the most intrinsically flexible, loosely coupled basin in the entire landscape (Figs. 2B-2C and 3C; S2 Table). In this intermediate the internal spine bridge is established, contracting from 7.9 Å in the two +CCNB1 states to 4.0 Å in State 9 (Note 3 in S1 Text), while the activation loop is already canonically BLAminus, indicating that ATP maximizes conformational sampling and completes the internal spine, leaving the final tensioning step to phosphorylation.

#### Activating Phosphorylation (pT161) Operates as a Mechanical Lever

Activation-loop phosphorylation at T161 is the structural determinant that locks this fluidized intermediate into a functional active site. The pT161 Holo complex escapes State 9 into a mature catalytic ensemble (State 1, 93%) that is uniformly Active BLAminus, αC-In and DFG-in. The difference is not a uniform compaction. In the phosphorylated ensemble the top of the active site is closer while the deep scaffold and hydrophobic shell are further apart, and the active-site elements (HRD, P-loop, DFG), uncorrelated beforehand, become strongly correlated with one another (Note 4 in S1 Text). In doing so pT161 re-establishes localized tension (Figs. 2A and 3E), leaving a more strongly coupled ensemble at the active site.

#### Inhibitory Phosphorylations (pT14/pY15) Warp the Active-Site Geometry

While single phosphorylation at T161 is the catalytically active condition for CDK1, triple phosphorylation at T14/Y15/T161 is known to provide an intermediate state [24] before the final active form (pT161; triple-P in Fig. 2). To resolve how the inhibitory phosphorylations at T14 and Y15 shut CDK1 down, we applied AlloQuant to a series of triple-P ensembles (S1 Fig). Strikingly, the inhibited triple-phosphorylated complex co-populates the same mature State 1 ensemble as the active pT161 complex (92% versus 93%), with statistically indistinguishable global rigidity and almost no allosteric rewiring (Figs. 2 and 3D; Note 5 in S1 Text). Instead, inhibition is executed by sub-angstrom distortions confined to the nucleotide pocket. The DFG motif is compressed against the catalytic Mg^2+^ while the P-loop is pushed away from ATP, breaking the geometric tolerances required for phosphoryl transfer without dismantling the kinase (S1 Fig; Note 5 in S1 Text).

The pT14 phosphate, on the glycine-rich P-loop, projects directly onto the reactive β/γ-phosphates of ATP, apposing two like-charged phosphate clusters at the very site of the transferable phosphate (S1 Fig). This is a direct electrostatic clash, not metal sequestration as the Mg^2+^ ion stays canonically coordinated by the ATP phosphates and the DFG aspartate. On the other hand, the second mark, pY15, appears to point away from the pocket. Thus, our triple-P ensemble suggests that pT14 is the dominant structural actor of this inhibitory control in our model. Previous studies predicted that T14 phosphorylation along pY15 would further block the peptide-binding site and might prevent either the binding or the appropriate configuration of ATP in CDK2-CCNA complex [25] and the inhibitory phosphorylation at the two P-loop sites may have separable mechanics and functions [24, 26]. Our ensembles suggest that CDK1, while the nucleotide stays bound, pT14 disrupts the configuration of its transferable phosphate. The complementary prediction of substrate-site blockade lies outside our models that contain no peptide substrate, however.

### Model 2: CSK-SRC (kinase-kinase heterodimer)

#### ATP Loading Locks Monomeric CSK Out of the Active Conformation

Before examining the heterodimer, we asked whether monomeric CSK could achieve the active αC-In conformation without a docking partner. AlphaFold3 predictions of CSK alone (full-MSA, 45 independent seeds, n = 225 per condition) show that apo CSK spontaneously adopts the pre-active geometry. All 225 apo models are αC-In (100%) and 96% achieve the BLAminus DFG state. Strikingly, ATP loading imposes a severe conformational blockade. Only 1 of 225 holo-CSK models is αC-In (0.4%; Fisher’s exact p = 4×10^−132^). Nucleotide coordination alone therefore excludes the active αC-In conformation in isolated CSK, raising the hypothesis that a docking partner is required to overcome this blockade (S2 Fig).

#### SRC Docking and Autophosphorylation Prime the Heterodimer

CSK phosphorylates the C-terminal tyrosine of SRC (Y530 in human SRC; homologous to Y527 in chicken SRC), yet the extensive physical docking between the two kinases (Fig. 4A) implies a mechanism beyond simple tail presentation. Using a Truth Overwrite algorithm that detects where AlphaFold3 overrides the designed apo/holo ligand configuration, we found that the presence of an active, phosphorylated SRC significantly re-allocates ATP into CSK in simulations (S3 Fig; Note 6 in S1 Text), thus suggesting, though AlphaFold3 reports no energies, that the SRC in dimer lowers the barrier to CSK ATP coordination and primes the complex for trans-phosphorylation, a piece of evidence that engaging SRC exerts allosteric effects on CSK.

**Figure 4.**
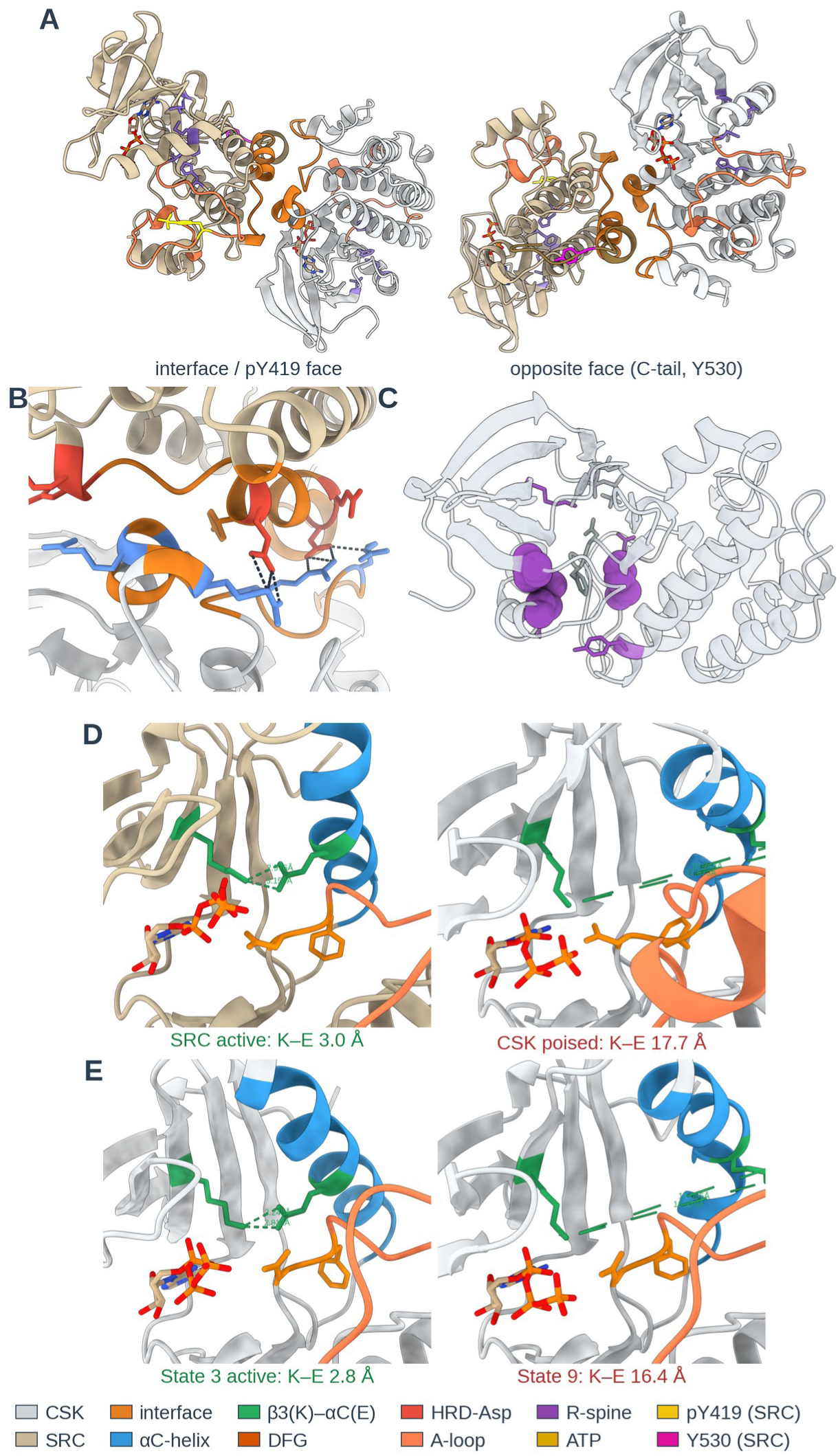
The structural basis of the CSK–SRC trans-allosteric handshake. Representative AlphaFold3 models of the CSK–SRC heterodimer (primed condition, csk-holo/src-py159-holo). Elements are colored throughout: CSK (light grey) and SRC (tan) kinase domains; docking interface (orange); αC-helix (blue); the β3-Lys–αC-Glu (β3K–αCE) salt bridge (green); DFG (dark orange); catalytic HRD-Asp (red); activation loop (coral); regulatory spine (purple); and ATP (gold). In (A), the SRC autophosphorylation site pY419 (yellow) and the SRC C-terminal substrate tyrosine Y530 (magenta) are also shown. **(A)** The docked kinase–kinase complex in two views 180° apart, engaging through a C-lobe:C-lobe interface. Unlike a kinase–cyclin complex, both partners are catalytic domains. **(B)** The docking interface: a basic CSK surface (αD/αG arginines) engages an acidic SRC αI surface via a salt-bridge network (dashed). **(C)** The coupled regulatory-spine module (purple) forms a connected column through the CSK core, spatially distinct from the catalytic/active-site machinery: the structural correlate of the spine-concentrated coupling in Fig. 6. **(D)** The single-structure handshake in one representative complex: an active SRC (β3K–αCE salt bridge formed, 3.0 Å) docked onto a poised CSK whose salt bridge remains broken (17.7 Å), the dominant SRC-State 8 × CSK-State 5 pairing of Fig. 5B (25 of 100 models). **(E)** The fully active CSK State 3 (salt bridge closed, 2.8 Å; shown left) versus the incompletely assembled State 9 (broken, 16.4 Å; shown right). Only 28% of the primed ensemble reaches State 3. Panels D and E share a common active-site orientation focused on the β3K–αCE (K–E) salt bridge. Quoted K–E distances are the minimum β3-Lys Nζ to αC-Glu carboxylate-oxygen distance (AlloQuant SB_Dist).

AlloQuant also reported that SRC holds an active global signature (BLAminus, αC-In) in essentially every docking state (S4A Fig), and its bulk rigidity is constant across the biological conditions (Note 7 in S1 Text; S3 Table). Autophosphorylation at Y419 (equivalent to Chicken SRC Y416) changes the conformations that SRC occupies. It redistributes nucleotide-loaded SRC from a single metastable state into a two-state ensemble (State 4 to States 7 and 8; Fig. 5A). Against the State 4 baseline the changes spread across the K–E salt bridge and the catalytic cleft, with none exceeding 0.7 Å (S7A and S7B Fig; Note 7 in S1 Text), and the internal spine bridge, already closed to 4.3 Å by ATP loading, moves by less than 0.15 Å. SRC’s contribution is therefore a phosphorylation-gated redistribution.

**Figure 5.**
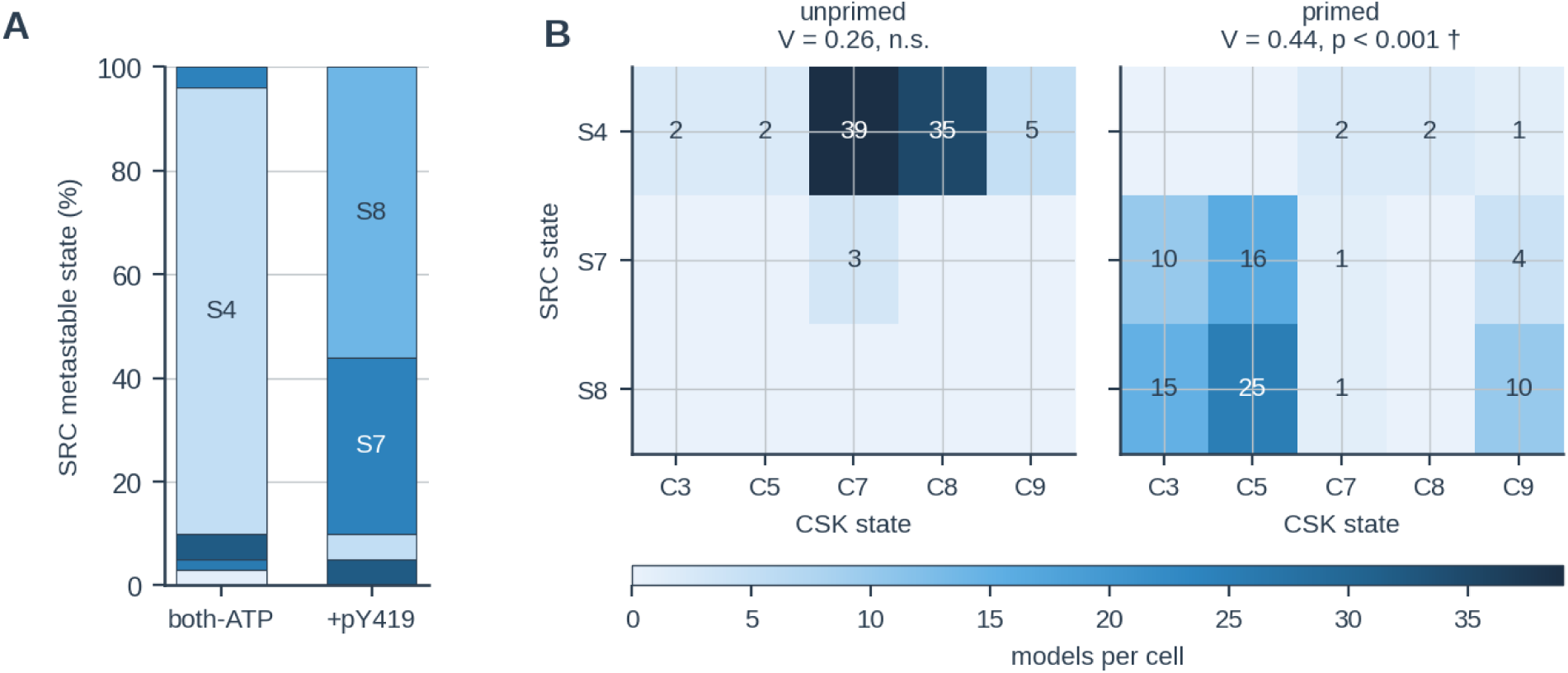
SRC autophosphorylation drives a conformational handshake that commits a subpopulation of CSK to the active state. AlloQuant analysis of the two nucleotide-loaded biological CSK–SRC conditions: both-ATP (csk-holo/src-holo) and primed (csk-holo/src-py159-holo, SRC pY419). Overwrite-artifact and non-biological conditions are excluded (Methods; S5 Table). *n* = 100 models per condition. **(A)** SRC metastable-state populations, colored on the intrinsic rigidity ramp. Nucleotide-loaded SRC lacking the mark occupies a single state (State 4, 86%), whereas pY419 redistributes the ensemble into States 8 (56%) and 7 (34%) at unchanged bulk rigidity (Note 7 in S1 Text). **(B)** Single-structure state handshake: per-model contingency of SRC state (rows) against CSK state (columns) in the both-ATP and primed complexes, cells colored and labeled by model count. Cramér’s *V* and its permutation *p* use the full table, not only the dominant states shown (Methods). Without the mark the two domains’ states are uncoupled (*V* = 0.26, *p* = 0.23, n.s.); in the primed complex they are coupled (*V* = 0.44, *p* < 0.001), the active SRC States 8 and 7 co-occurring with the more active CSK States 5, 3 and 9 in the same structures. CSK’s active-state signatures: S4 Fig. †, this value is limited by the number of permutation draws, 0.0010 at 10,000 draws and 0.00066 at 200,000 (Methods).

#### SRC Priming Commits a Subpopulation of CSK to the Active State

Because CSK lacks typical positive regulatory phosphorylation sites, how it reaches its active state has remained debated. We found CSK progression to be strongly SRC-dependent (Note 9 in S1 Text). With SRC unphosphorylated, the predicted CSK ensemble is dominated by an inactive basin with a fully extruded αC-helix, and loading ATP without the SRC mark leaves that dominance intact (CSK States 7 and 8, together 81% of that ensemble, neither containing a single αC-In or Active BLAminus model; Fig. 5B, left). SRC pY419 shifts the balance, raising CSK’s ensemble-level Active BLAminus population from 17% to 47% and αC-In from 5% to 27% (versus the CSK-ATP condition; S4B Fig). These per-condition averages understate the mechanism. CSK activation is not graded but confined to a single metastable state, State 3 (Fig. 4E, left). This state occupies 28% of the primed ensemble and is 98% αC-In and 98% Active BLAminus, whereas States 1, 2, 5, 7 and 8 contain no αC-In model at all, and the residual BLAminus population comes from States 9 and 4, which adopt the active backbone with the αC-helix still extruded (Fig. 4E, right; S4 Table; S4D Fig). The CSK-SRC docking interaction reported by the Kuriyan group is therefore not merely a proximity device, yet full catalytic commitment requires steps beyond SRC docking alone. In their crystal structure neither chain adopts an active conformation, by AlloQuant’s classification and Kincore’s alike (S1 Table). Differential driver analysis confirms that this activation is a coordinated active-site rearrangement. The internal catalytic salt bridge and deep scaffold close while the substrate cleft opens (Note 9 in S1 Text). The docking interface itself is not remodelled, however. Between the unprimed and primed SRC states the interface residue pairs shift by 0.07 Å on average, below AlphaFold3’s own predicted aligned error at those positions, and 30 of the 31 interface residues keep the same nearest partner (Note 8 in S1 Text). The pooled MAC rises in the primed dimer simply because the ensemble has shifted to different states, not because any state became more coupled (Note 10 in S1 Text; S7 and S8 Tables). SRC priming therefore acts by selecting an active conformation of CSK that is already accessible within the docked complex, rather than by deforming the receiver.

#### The Two Kinases Are Conformationally Coupled Within Individual Complexes

Because each AlphaFold3 model pairs one CSK and one SRC domain in a single structure, we could ask whether the two kinases’ conformations are coupled model-by-model, at the level of discrete states and of continuous geometry alike. The reciprocity between the partners is conformational rather than bulk-mechanical, and nucleotide’s own influence is local, not a reorganization of bulk coupling (Note 11 in S1 Text). Within the pY419-primed complex the SRC and CSK metastable states are significantly associated structure-by-structure, the active SRC states co-occurring with the more active CSK states in the same models, a coupling absent from the unprimed complex and therefore gated by the SRC activation-loop mark (Fig. 5B; Note 11 in S1 Text).

Correlating every SRC internal metric against its CSK counterpart across the 795 paired models, computed within each condition and with model confidence partialled out, revealed a specific, reproducible coupling concentrated in the regulatory-spine module, whereas the principal catalytic and active-site elements showed none (Figs. 4C and 6; Note 12 in S1 Text). Although individually modest in magnitude, these correlations are highly significant and robust to both experimental design and model confidence. Thus, within individual complexes, the spine elements of the two kinases co-vary across models while their catalytic cores vary independently, a single-structure correlate of trans-allosteric coupling between the two regulatory spines across the docking interface (Fig. 4D).

**Figure 6.**
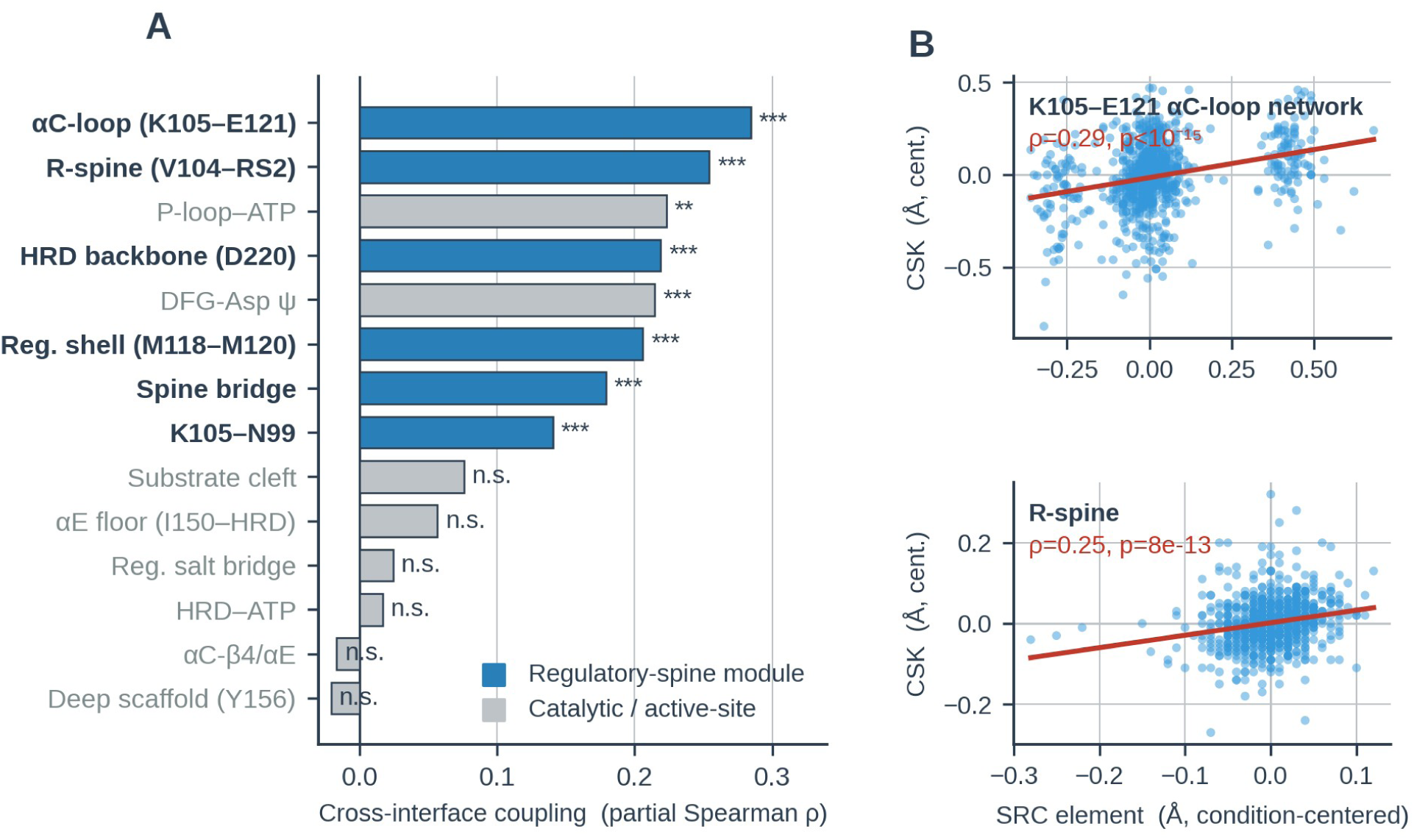
Homologous regulatory-spine elements of CSK and SRC are conformationally coupled within individual predicted complexes. Because each AlphaFold3 model contains one CSK and one SRC kinase domain, internal geometric metrics of the two domains can be correlated model-by-model. All correlations are computed *within* each experimental condition (Spearman ρ, meta-analyzed across the seven conditions) and, for the spectrum in (A), partial out model confidence (ipTM and interface PAE), so that neither the shared experimental design nor prediction quality can account for the signal (*n* = 795 paired models). **(A)** Cross-interface coupling spectrum: within-condition partial Spearman ρ between each SRC element and its CSK counterpart. The regulatory-spine module (blue) comprises the K105–E121 αC-loop network, the regulatory R-spine (V104–RS2), the regulatory shell (M118–M120), the HRD backbone (D220), the internal spine bridge, and the K105–N99 contact. All six couple significantly across the docking interface. Of the eight catalytic and active-site elements (grey), only two do: the P-loop–ATP contact and the DFG-Asp ψ backbone dihedral (*** *p.adj* < 10^-3^, ** *p.adj* < 10^-2^, adjusted across the fourteen elements shown; n.s., not significant). **(B)** Condition-centered co-variation for the two most strongly coupled elements (each point one model, values centered on the per-condition median): the K105–E121 αC-loop network (unadjusted ρ = 0.29) and the regulatory R-spine (ρ = 0.25); red lines are linear fits. The coupling is concentrated in, though not strictly confined to, the regulatory spine (Note 12 in S1 Text).

## DISCUSSION

AlloQuant revealed a shared allosteric principle from two kinase models: 1) in neither system can the kinase reach a catalytically competent conformation on its own, and 2) in neither does the partner act by proximity alone. Both require physical engagement of the receiver kinase (CDK1 or CSK) with another protein as the activator in the complex, and in both the engagement acts by changing which conformations the receiver occupies rather than by deforming a single structure along a continuous path. Their modes of action, however, differ considerably in the constitutive status of the activator, in whether engagement remodels the receiver’s internal mechanics at all, in where the activation barrier sits, and in how completely the transition is executed.

The two receivers are trapped in opposite ways. Counter to the expectation that a dormant kinase is conformationally loose and that activation rigidifies it [1, 16], monomeric CDK1 is locked in a hyper-coupled basin behind a catalytically incompetent αC-Out face. The engagement of the cofactor CCNB1 releases CDK1 rigidity and ATP relaxes the node network further (Note 2 in S1 Text). In contrast, CSK’s trap is the mirror image. Whereas apo CSK already samples a pre-active geometry almost exclusively, nucleotide coordination excludes the active conformation. Unlike CDK1, engaging a primed SRC leaves CSK’s coupling within states unchanged, and what the partner moves is the ensemble itself (Note 10 in S1 Text). Partner-driven activation without global mechanical change has precedent. TPX2 binding produces no global conformational change in Aurora A, locking the active conformation by locally repositioning the activation segment [27]. Thus, a contrasting theme is that while a dedicated cofactor (i.e., CCNB1) releases tension in an over-constrained receiver, a docked kinase (i.e., SRC) leaves its receiver’s mechanics as it finds them and re-weights the states the receiver occupies. What the two share is not a change in the receiver’s mechanical set-point but that the partner determines which conformations the receiver can reach.

The two dimer models also differ in how far the activator drives the transition. CCNB1 is a constitutive scaffold, establishing a near-uniform Active BLAminus architecture before any phosphorylation. While the pT161 complex converges on one mature ensemble, pT14/pY15 keeps it from catalytic engagement (Note 5 in S1 Text) until those marks are removed [24]. An obligate switch at the mitotic commitment point is well served by a cofactor, an activating mark and a licensing dephosphorylation acting in concert to drive the transition to completion. In contrast, SRC is a conditional activator. As it is the substrate, its capacity to prime CSK is gated by its own autophosphorylation. SRC pY419 commits a subset of CSK molecules fully rather than moving all of them part-way. A tonic regulator of Src-family activity may be better served that way, setting a tunable steady-state flux of tail phosphorylation, not a binary output. Despite these differences, both models appear to need a final dephosphorylation before the output is realized. The marks sit on opposite partners, removal completing the receiver’s activation in CDK1 and the activator’s inhibition in the CSK-SRC pair. Autophosphorylated SRC accepts the Y530 mark while remaining active [8, 9]. As in the triple-phosphorylated CDK1 state carrying pT14/pY15 and pT161, the inhibitory mark CSK writes on a primed SRC may stay silent until the activation loop is dephosphorylated (Note 16 in S1 Text). pY419 therefore plays two roles: one keeping SRC active despite the inhibitory mark and the other licensing CSK to write it.

CSK-dependent regulation in cells also relies on N-terminal domains absent from our models, and they serve different ends on the two chains. SRC’s SH2 domain is what makes the Y530 mark consequential, engaging the phosphorylated tail to assemble the autoinhibited state [6]. CSK’s SH2 and SH3 instead position the kinase. An SH3-deleted CSK fails to suppress SRC in cells while remaining fully active on substrates in vitro [28]. The SH2 domain anchors CSK through phosphotyrosine ligands such as paxillin and focal adhesion kinase [29], thus recruiting CSK to the membrane and raising its kinase activity [30]. Such scaffolds raised by either or both SH2 and SH3 domains may provide a stage for phosphatases to finalize SRC inactivation. PTEN dephosphorylates SRC directly at the Y419-equivalent site [31], and PEST-family phosphatases act at the homologous site in Lck [32]. One PEST-family member binds CSK through its SH3 domain, the kinase and the phosphatase then cooperating to suppress SRC-family activity in a manner requiring their association [33]. Therefore, the SH3 domain our construct lacks may also be able to recruit the enzyme that makes CSK’s mark count.

Conformational state is nonetheless what sets CSK’s output. Acting on a SRC that cannot autophosphorylate, full-length CSK shows biphasic kinetics, implying two CSK forms and so more than one route into the active state [34]. Those forms were left structurally unassigned, and the interdomain motions considered there lie outside our construct. Our unprimed complex names a candidate inside the kinase domain, where SRC is docked but unmarked and only 5% of CSK molecules reach the αC-In geometry, leaving the majority form one whose active site is not assembled. Their two forms are distinguished by affinity for SRC and ours by active-site assembly, a correspondence that remains to be tested.

Because the CSK-SRC model contains two kinase domains, the kinase-kinase pair permitted a test unavailable in the cofactor system: whether the partners’ conformations are coupled structure by structure. They are, and pY419 gates it. In the primed complex SRC’s state predicts CSK’s, and in the unprimed complex it does not (Cramér’s V = 0.44 versus 0.26, n.s.). The coupling is concentrated in the regulatory-spine module rather than the catalytic machinery (795 paired models). The reciprocity is therefore conformational and pY419-gated, not bulk-mechanical. SRC’s global rigidity is indifferent to CSK’s nucleotide status. This also circumscribes the “anvil” metaphor. What pY419 gates is the occupancy of docking-competent conformations, not a remodeling of the docking surface, whose contacts are essentially invariant across SRC’s states. The two spines themselves sit some 50 Å apart, well distal to the interface, arguing for allosteric transmission through the assembled architecture rather than direct spine-to-spine contact.

Two limits frame these conclusions. AlphaFold3 returns discrete states with no explicit time axis. Thus, our metrics describe the geometry and relative populations of conformations encoded in the model’s learned priors, not measured motions, rates, or thermodynamics. The models also contain isolated kinase domains and no substrate. For CSK the cost of that construct is known. The isolated catalytic domain is some 100-fold less catalytically efficient in vitro than full-length CSK, and an excess of the CSK SH3 domain supplied in *trans* recovers only a small part of that deficit [35]. Our models contain no SH3 domain, so whether the contact is intramolecular is untested here. The SRC SH2/SH3 modules and linkers that constrain tail accessibility are absent, so productive presentation of Y530 is not addressed, and the predicted substrate-site blockade by pT14/pY15 lies outside a substrate-free model.

Through the analysis of two model kinase dimers, we have demonstrated the general utility of AlloQuant for documenting allosteric interactions in kinase complexes. Because the structures were generated by AlphaFold3, our analysis offers hypotheses, not proofs. The central one is that heterodimer partners govern kinase activation by setting which internal conformations the receiver can occupy, and that this control, beyond proximity or orthosteric contact, may itself be a determinant of the active-state transition. Three predictions follow. Interface mutations that disrupt the primed SRC docking conformation, or the CCNB1 contact, should impair activation without abolishing binding. Methods sensitive to conformational distributions (hydrogen–deuterium exchange, NMR relaxation, single-molecule FRET) should detect CDK1 decoupling on CCNB1 engagement, and for CSK a redistribution of state populations on SRC pY419 rather than a change in bulk flexibility. And the sub-stoichiometric CSK transition predicts a lower intrinsic catalytic efficiency for CSK than for cofactor-completed kinases such as CDK1. AlloQuant is agnostic to the ensemble’s origin. It annotates the CSK-SRC crystal structure as readily as predictions and extends to molecular-dynamics trajectories. Because it assigns landmarks by profile-HMM alignment, these tests are tractable across the kinome.

## METHODS

### Structural Data Generation and Preprocessing

Predicted structural coordinates and physical state parameters were generated using AlphaFold3 [36]. Output ensembles in PDBx/mmCIF format were aggregated and parsed utilizing a custom Python parsing engine. To establish a universal spatial coordinate system across the structurally diverse KINOME, simulated sequences were aligned against the reference Protein Kinase domain Hidden Markov Model profile (Pfam accession PF00069) utilizing the hmmalign function in HMMER (v3.4). Sub-angstrom structural metrics and dihedral angles corresponding to these landmark residues were subsequently extracted via headless batch execution of UCSF ChimeraX (v1.11.1). To eliminate cross-chain ligand crosstalk (an artifact where distances to ligands situated in the adjacent subunit kinase are erroneously recorded for the apo target subunit), a dynamic correction algorithm was applied. Specifically, if the target subunit was strictly defined as ‘apo’ in the experimental design matrix, all physical coordinates relating to ATP or Mg^2+^ were explicitly masked prior to downstream analysis. Distance metrics were subsequently classified into universal (present across all conditions) and multi-group variables to maintain robust sample sizes during statistical testing. To evaluate how AlphaFold3 allocates the ligand between the two subunits, a "Truth Overwrite" discrepancy algorithm was implemented. While multimeric AlphaFold 3 models were seeded with distinct user-defined ligand states (e.g., ATP presence/absence), the algorithm evaluated the final 3D physical coordinates to detect ligand re-allocation. Instances where the network placed the ligand away from the seeded site, ejecting or misaligning it. A true "Holo" physical state was strictly defined by the spatial proximity of the ATP molecule to both the catalytic spine (C-spine) and the conserved HRD motif aspartate within the predicted geometry, regardless of the initial input seed. Instances where the predicted geometry significantly diverged from the intended input configuration were quantified as ligand re-allocation events.

Because AlphaFold3 is free to place a seeded ligand where its own prediction places it, the designed apo/holo configuration is not necessarily the one realized in the returned coordinates. A Truth Overwrite check was therefore applied to every model. The final ATP geometry was scored against both the catalytic (C-)spine and the HRD-motif aspartate, and a model was flagged as overwritten when that geometry no longer matched its seeded assignment. Each model was then attributed to the condition it physically realized rather than the one designed, which is why per-condition counts in the reviewed ensembles are unequal. Two designs proved overwritten in the large majority of models (csk-apo/src-holo, 98%, and csk-apo/src-py159-holo, 87%, against 2% for the reciprocal csk-holo/src-apo) and are reported as overwrite artifacts; together with the non-biological designs carrying pY419 on an otherwise apo SRC, they are excluded from the biological comparisons of Fig. 5 (S3 Fig; Note 6 in S1 Text).

For mechanistic interpretation we further restricted attention to physiologically plausible ligand/phosphorylation combinations. Designs pairing SRC activation-loop phosphorylation (pY419) with a nucleotide-free SRC (src-py159-apo) were treated as non-biological, because Y419 autophosphorylation is a product of the catalytically active, ATP-bound SRC, and a phosphorylated-yet-apo SRC does not represent a stable physiological state. SRC autophosphorylation in vitro strictly requires ATP-Mg [9].

### Categorical Structural State Analysis

Categorical structural classifications were evaluated across all experimental complexes by mapping canonical kinase features. These included the activation loop DFG (Asp-Phe-Gly) motif spatial configuration, the αC-helix orientation ("In" vs. "Out"), and the structural integrity of the Regulatory (R-spine) and Catalytic (C-spine) hydrophobic networks [17]. Spatial classifications into standardized active/inactive nomenclature were defined according to established Dunbrack distance coordinates (D1 and D2) [20]. To determine the statistical enrichment of canonical active conformations driven by the binding partner’s state, features were binarized (Target State vs. Other). Pairwise Fisher’s exact tests were performed across all complex combinations, with *p*-values adjusted for multiple comparisons utilizing the Benjamini-Hochberg False Discovery Rate (FDR) procedure [37].

### Micro-Metric Evaluation and Allosteric Network Coupling

Continuous structural variables (e.g., inter-residue atomic distances) were evaluated for condition-dependent shifts. Omnibus variance across experimental groups was assessed utilizing the Kruskal-Wallis rank-sum test, followed by pairwise Wilcoxon rank-sum tests with Benjamini-Hochberg false discovery rate (BH-FDR) correction utilizing the *rstatix* package (v0.7.2). To map internal allosteric networks and quantify mechanical signal propagation, Spearman’s rank correlation (*ρ*) matrices were computed for all inter-residue distance metrics within each experimental condition.

To quantify the global structural tension of the kinase domain, we defined a Global Network Rigidity score based on the Mean Absolute Correlation (MAC) of pairwise inter-landmark distance fluctuations. This approach is conceptually related to prior correlation-based analyses of intramolecular coupling, in which highly correlated fluctuations are interpreted as evidence of a mechanically coupled, rigidified network, whereas low correlation indicates independent, decoupled flexibility [38]. MAC is analogous in interpretive rationale to the correlated-motion framework of McClendon et al., though it differs methodologically in both the underlying variable (inter-landmark distances rather than dihedral torsions) and the correlation statistic employed (signed Spearman rank correlation rather than mutual information). Because ρ can be positive or negative, simple averaging of the raw correlation matrix would allow strongly anti-correlated (inversely coupled) distance pairs to numerically cancel strongly positively correlated pairs, understating the true extent of network coupling. We therefore computed MAC by averaging the absolute values of all unique pairwise correlations (the strict upper triangle of the ρ matrix), so that both direct and inverse coupling contribute additively to the rigidity score.

Because distance availability is condition-dependent, MAC was calculated over two distinct distance sets. In the ATP-loaded CSK conditions, 21 of 23 inter-landmark distances were resolvable, yielding 210 unique pairwise edges; in the apo condition, only 16 of 23 distances were resolvable (120 edges), since ATP- and Mg-anchored metrics are undefined in the absence of bound nucleotide (global MAC). For the unsupervised metastable-state analysis (below), we restricted to the 16 distances populated in every condition, providing a common basis for comparing MAC across states (Intrinsic MAC). MAC is formally defined as:

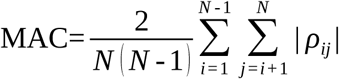

Where *N* represents the total number of universally shared inter-residue distance metrics within the structural ensemble, and *ρ_ij_* is the Spearman rank correlation coefficient between distance metric *i* and distance metric *j*. High MAC scores (e.g., *MAC* ≫ 0.15) mathematically indicate a tension-loaded, "stiff" conformation restricted to a low number of degrees of freedom, while significant reductions in MAC denote allosteric decoupling, in which those elements vary more independently of one another. MAC is additionally computed within individual metastable states; see Unsupervised Metastable State Discovery, below. Because the 210 edges are constructed from 21 shared metrics, they are not independent observations, so the pairwise tests on edge distributions overstate the effective sample size. The p-values are reported as a consistent ranking of contrast strength rather than as calibrated significance, and no physical claim in this work rests on a p-value alone.

To exclude the possibility that condition-dependent MAC differences reflect differences in AlphaFold3 prediction confidence rather than mechanical coupling, the correlation network was recomputed with per-model confidence partialled out of every edge. Each Spearman *ρ* was replaced by a partial Spearman correlation conditioned on the per-model confidence covariates (ipTM and mean interface PAE for the heterodimer ensembles; pTM for the CDK1 series, whose apo conditions are monomeric and therefore have no interface metrics), applied sequentially by the closed-form partial-correlation identity, and the between-condition contrasts were re-tested on the partialled edge distributions. Because partialling removes variance, it lowers |*ρ*| and hence MAC in every group, so the quantity under test is the between-condition contrast, in direction and significance, not the absolute MAC value (S6 Fig; Note 13 in S1 Text; S5 and S6 Tables).

### Cross-Chain Conformational Coupling (Paired Heterodimer Analysis)

Because each multimeric AlphaFold3 model contains both kinase domains in a single structure, the per-model geometric measurements of the two chains are naturally paired. To test whether the two domains’ conformations are coupled at the level of individual predicted structures, the per-chain metric tables were joined on their shared model identifier and, for every pairing of one enzyme-chain metric with one substrate-chain metric, a Spearman rank correlation was computed *within each experimental condition* and combined across conditions by inverse-variance-weighted Fisher *z* meta-analysis; this within-condition design removes any correlation induced purely by the shared experimental grouping. To exclude a prediction-quality artifact, correlations were additionally recomputed as partial Spearman correlations controlling for per-model AlphaFold3 confidence (ipTM and mean interface PAE). Significance across the full metric grid was controlled by the Benjamini-Hochberg FDR procedure. Coupling at the level of discrete conformational states was assessed separately. Within a single experimental condition, the two chains’ GMM metastable-state assignments were cross-tabulated and their association quantified by Cramér’s *V*, with significance determined by a label-permutation test (10,000 permutations of one chain’s state labels, repeated at 200,000 permutations to bound the Monte Carlo error on the resulting p-value. For the primed complex the two draw counts gave p = 1.0×10-3 and 6.6×10-4, and the unprimed comparison was non-significant at every draw count tested) to provide a distribution-free *p*-value robust to sparse contingency cells. This paired analysis is implemented as a standalone add-on and does not modify the core per-chain pipeline.

### Structural Analysis of the Docked Complex

Representative structures for visualization and geometric measurement were selected as per-condition or per-state centroids. The model minimizing the standardized Euclidean distance to the ensemble mean across the panel of inter-residue metrics. Per-residue flexibility (S5A Fig) was quantified as the Cα root-mean-square fluctuation (RMSF) across the primed ensemble after least-squares superposition on the rigid, combined C-lobe cores of both chains. Inter-chain interface residues were defined as those contributing any heavy atom within 4 Å of the partner chain, and interface polar contacts were identified in UCSF ChimeraX; inter-domain, regulatory-spine, and C-terminal-tail distances reported in the text and figures were measured between the corresponding Cα centroids or side-chain atoms of these representative models. Structure rendering used UCSF ChimeraX.

For the CDK1–CCNB1 complexes, RMSF was computed chain-locally. Each chain was fitted on its own conformation in the first model of that condition using its Cα atoms only, so that motion of one subunit relative to the other is excluded and the value reports internal deformation alone. This differs from the shared-frame superposition used for S5A Fig, which retains inter-chain motion, so the two sets of values are not directly comparable. Per-residue RMSF was taken about each chain’s mean position across the 100 models of the condition, over residues resolved in every model (CDK1 297, CCNB1 268).

Because the interface displacements under test are sub-angstrom, they were compared against AlphaFold3’s own estimate of coordinate uncertainty at the same positions. For each model the predicted aligned error (PAE) matrix was read from the per-model confidence output and reduced to residue resolution. Tokens were grouped by chain and residue identifier, which is necessary because modified residues such as the SRC pY419 phosphotyrosine are atom-tokenised rather than residue-tokenised. The minimum PAE was taken over each residue block, and the matrix was symmetrised as (PAE*_ij_* + PAE*_ji_*)/2, PAE being defined asymmetrically as the expected positional error of one residue when the structure is aligned on another. Taking the block minimum is the most favourable reading of the model’s own confidence, so the resulting uncertainties are lower bounds. Each interface contact pair’s mean displacement between SRC metastable states was then expressed as a fraction of that same pair’s mean PAE across the ensemble. The same reduction was used to compare confidence between states within defined residue blocks: the interface contact set, each chain’s C-lobe core, and the full inter-chain block. These two quantities answer different questions and are reported separately. Displacements between ensembles are highly reproducible in the statistical sense, the between-model spread at these pairs being roughly an order of magnitude smaller than the PAE, so they are significant within the ensemble. PAE instead bounds what a displacement of that size can be taken to mean about the physical structure, and is used here for that purpose alone (Note 8 in S1 Text).

### Unsupervised Metastable State Discovery

To identify distinct conformational basins, universally viable pairwise distance metrics were projected into a reduced continuous phase space using Principal Component Analysis (PCA). Unsupervised metastable state discovery was performed on the principal components representing the majority of shared structural variance utilizing Gaussian Mixture Models (GMM) via the *mclust* package (v6.1.2) [39]. GMM was specifically selected over strict spherical-partitioning algorithms to accurately accommodate the elongated, ellipsoidal data distributions characteristic of continuous protein conformational dynamics. The optimal number of underlying metastable states (*k*) was determined automatically by maximizing the Bayesian Information Criterion (BIC). Methodological validation of the clustering was performed using *K*-means, evaluated by the gap statistic [40] utilizing a global standard error maximum rule (globalSEmax) to prevent premature local minima trapping along elongated coordinate axes. Throughout, “metastable state” denotes a GMM component of the predicted structural ensemble, an operational and statistically defined conformational basin. It is not a kinetically characterized species, since AlphaFold3 samples structures independently and reports no dynamics.

The MAC defined above is applied at two levels of grouping, which are related but not interchangeable. Global (condition-level) MAC builds one correlation network from all models of a single experimental condition, and so describes the ensemble that condition produces. Intrinsic (within-state) MAC builds one network from the members of a single metastable state, pooled across conditions, and so describes that conformational basin itself. The distinction matters because a global value depends on the state composition of the condition it is computed over, and does so in either direction. Mixing structurally distinct populations adds spread between basins, which dilutes correlation, so a condition spread across several states can yield a lower global MAC than its constituent states show individually; CDK1-CCNB1+ATP has a global MAC of 0.12 while being 92% State 9, whose intrinsic MAC is 0.15. But mixing also induces correlation among every metric that separates those states, which inflates it, and this term dominates when a condition splits into a genuinely new population, as the primed CSK ensemble does (Note 10 in S1 Text). A global MAC difference between two conditions is therefore not interpretable as a difference in internal coupling until the state composition of both has been accounted for. The two quantities were tested separately, by pairwise Wilcoxon rank-sum on the corresponding edge distributions with BH-FDR correction applied within each family. A significant difference between two conditions accordingly neither implies nor is implied by a difference between their dominant states. As a sensitivity check, the 26 models from the residual csk-apo/src-py159-holo condition (n=33 post-overwrite) assigned to States 7–8 were excluded from the intrinsic-MAC computation. Removing them increased MAC for both states and strengthened two comparisons from ** to ***; no significant result lost significance, confirming the full-pool analysis is conservative (Note 14 in S1 Text).

MAC and Cα root-mean-square fluctuation (RMSF) are both used here as measures of how rigid a protein is, and either can be computed on a trajectory or on a structural ensemble, but they capture different aspects and can move independently. MAC is a correlation taken over pairs of inter-landmark distances. It reports whether parts of the fold change together, and because it is built from rank correlations it is invariant to the amplitude of those changes. RMSF is an amplitude taken per residue about a mean position. It reports how far a part moves, and is indifferent to whether anything else moves with it. A fold whose small residual motions are tightly coupled therefore scores high on MAC and low on RMSF, and one whose large motions are independent does the reverse. The two are reported separately throughout and are never placed on a common axis. MAC additionally requires the landmark network of Module 1 and so applies only to kinase chains, whereas RMSF needs no landmarks and is the only one of the two available for the CCNB1 cofactor.

### Validation of Functional Signatures

Following the mathematical delineation of metastable basins, structural identities were mapped by cross-tabulating the unsupervised clusters against established binary categorical features. Statistical enrichment of specific biological signatures within each metastable state was evaluated utilizing Fisher’s exact tests. For highly dimensional contingency tables where computational limits were exceeded, Monte Carlo simulations (B=2000 permutations) were employed to estimate exact *p*-values. All significance thresholds were subjected to BH-FDR correction.

### Differential Allosteric Driver Analysis

To identify the specific micro-metric deviations driving the separation of structurally similar metastable sub-populations (e.g., conformational clamping within a shared functional metastable state), a distribution-aware differential distance analysis was conducted. For universally shared continuous metrics, distributions between selected metastable states were compared using the Wilcoxon rank-sum test with Benjamini-Hochberg FDR correction. To appropriately scale the magnitude of the structural shift against the inherent coordinate variance (flexibility) within each state, effect sizes were quantified using either parametric or non-parametric methods. Parametric effect sizes were calculated utilizing Cohen’s *d* derived from pooled standard deviations, with significance defined by an absolute effect size |*d*| > 0.5, a medium effect [41]. Alternatively, to robustly accommodate non-normal structural distributions and tied coordinate data intrinsic to discrete structural ensembles (e.g., identical atomic distances across multiple AlphaFold3 seeds), the non-parametric Rank-Biserial Correlation (*r_rb_*) was utilized [42]. To circumvent algorithmic limitations associated with tied ranks, *r_rb_* was derived mathematically from the Mann-Whitney *U* test statistic (*W*) via the exact transformation:

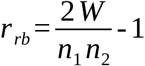

For non-parametric evaluations, significant drivers were defined by an absolute effect size ∣*r_rb_*∣ > 0.3, a threshold explicitly chosen to maintain mathematical and conceptual parity with a medium parametric effect size (*d* = 0.5) [41]. Across both methods, significant differential allosteric drivers were strictly required to satisfy an FDR-adjusted *p* < 0.05. We found that our two effect size approaches, Cohen’s *d* and *r_rb_*, generated similar results consistent with the *Central Limit Theorem* as our ensembles consist of 100 simulations per group (20 random seeds x 5 models). To provide physical directionality to the state transitions, the sign of the effect size was strictly oriented relative to the second group in the comparison (Group B). Therefore, a negative effect size (e.g., *d* < 0 or *r_rb_* < 0) indicates physical compaction in Group B relative to Group A (a shortening of inter-residue distances, such as interface crushing or loop closure), whereas a positive effect size indicates structural expansion or solvent-exposure.

All data preprocessing, multivariate clustering, statistical testing, and visualization were orchestrated utilizing custom Python/R (version 4.5.3) pipelines constructed in Ubuntu 22.04 and 24.04 environments. Key R packages utilized included *tidyverse* (v2.0.0) for data manipulation, *rstatix* (v0.7.3) for non-parametric statistics, *mclust* (v6.1.2) for Gaussian Mixture Modeling, *cluster* (v2.1.8.2) for Gap Statistic validation, *factoextra* (v2.0.0) for multivariate visualization, and *plotly* (v4.12.0) for interactive three-dimensional PCA visualization. The pipelines constructed for this study are posted in the GitHub repository (https://github.com/aimamoto/kinase_simulation_analysis). Main-text figures were regenerated for presentation from the pipeline’s computed outputs using a consistent color palette and type scale; the reported values are those produced directly by AlloQuant, the figure-generation scripts are included in the repository.

## Author Contributions

A.I. conceived the project and designed the overall study architecture. A.I., A.S., and M.O. refined the initial study design through discussions of the structural and biological aspects. A.I. performed the AlphaFold3 simulations and analysis of structural assessments of the structural ensembles. Y.W. set up the local computing environment and configured the TSUBAME4.0 supercomputer for executing the AlphaFold3 simulations. A.I., A.S., and M.O. wrote the manuscripts. A.S. and M.O. secured funding support for the study.

## Acknowledgements

The authors thank Josephina Sampson, Prashant Jain, Paola Laurino and the members of the A.S. lab for critical reading of the manuscript and for constructive comments. This work was supported by JST CREST Program (JPMJCR21N3) and JST ASPIRE Program (JPMJAP24B1) to M.O., and by MEXT JSPS KAKENHI (grant number 26K15044) to A.S. Computer resources were provided by the TSUBAME4.0 supercomputer through the HPCI System Research Project (Project ID: hp250051). This work was also supported by the World Premier International Research Center Initiative (WPI), MEXT, Japan. During the preparation of this work, the authors used Claude (Anthropic), an AI assistant, for language editing and analysis-code documentation; it was not used to generate or alter research data, results, or figures, and the authors reviewed and verified all content and take full responsibility for the work.

## Code and Data Availability

AlloQuant, including both analysis modules and the figure-generation scripts, is available at https://github.com/aimamoto/kinase_simulation_analysis and archived at Zenodo (doi:10.5281/zenodo.22208357). The per-model metric definitions (*AlloQuant_master_CSV_data_dictionary_v7r3*) and the output-file manifest (*AlloQuant_output_file_manifest_v7r3*) are in the repository’s docs/ directory and are also provided with this submission as S1 and S2 Dataset, so that every reported metric can be interpreted without consulting the repository. The AlloQuant outputs underlying every reported value are provided as S3 Dataset and archived in the same record. These comprise the per-model measurements for every ensemble analysed, the complete Module 2 statistical output, the derived interface analyses, the cross-chain coupling and prediction-confidence controls, and the 3D7T validation. The AlphaFold3 inputs defining every ensemble, the experimental design matrices and the construct sequences, are included in the same deposit, from which the ensembles can be regenerated. The predicted coordinates themselves are not deposited, running to several gigabytes.

## Supplementary Information

### Supplementary Notes

These notes hold the full quantitative basis (network-rigidity values, metastable-state populations, and sub-angstrom differential drivers with signed effect sizes) for the findings summarized in Results. Effect sizes are Cohen’s *d* (positive, distance increases; negative, distance decreases); network rigidity is the Mean Absolute Correlation (MAC); *p*.adj denotes Benjamini–Hochberg-adjusted *p* values.

**Note 1. External validation against an experimental CSK–SRC structure.** AlloQuant’s readouts are otherwise internal to the AlphaFold3 ensembles, so Module 1 was run blind on PDB 3D7T, the 2.9 Å crystal structure of the CSK–SRC kinase-domain complex [1], and every call was compared with Kincore [2], a condition-agnostic classifier that labels any coordinate file and therefore requires no ligand or construct matching (S1 Table). Where independent ground truth exists the agreement is exact. The DFG-Asp backbone dihedrals (CSK −112.49°/120.17°; SRC −79.66°/41.23°), both Dunbrack coordinates, both β3-Lys–αC-Glu salt-bridge distances (3.50 and 2.62 Å), the DFG spatial class, the αC orientation and the regulatory-spine state all match, and both tools score both chains non-active. AlloQuant additionally reports quantities that a single-structure description does not tabulate. It assigned the asymmetric roles blind, assigning CSK as the N-lobe receiver and SRC as the C-lobe donor, the same topology inferred from the ensembles, with reciprocal interface distances of 2.69 and 11.44 Å, and it separates the two chains’ internal mechanics within one crystal. The CSK spine is intact with a closed salt bridge, while the SRC spine is broken and DFG-out, the Src-like inactive conformation described previously [3]. Projecting both chains out of sample into the unsupervised state space (Methods) places CSK in State 3, the fully active basin, and SRC in State 2, each at posterior 1.00. The crystal was deliberately not added to the mixture. A validation structure that helps fit the model would then help define the basin it is assigned to, and refitting with it changes the Bayesian-information-criterion component count and relabels most published models. The comparison is bounded by the ligand. Staurosporine is ATP-competitive and mimics the adenine but carries no phosphate groups, and Kincore types it Type 1 in both sites, so an active-like CSK architecture is expected rather than surprising, and every phosphate- or Mg-anchored metric is undefined; CSK also carries K361A/K362A with a partly disordered activation loop, and at n = 1 per chain no MAC is defined. This benchmark therefore validates landmark placement, interface topology and the geometry of the state space in Module 1, not the trans-allosteric mechanism itself as designed in Module 2.

**Note 2. CCNB1 decouples the trapped CDK1 monomer.** Monomeric CDK1-Apo shows a global network rigidity of MAC = 0.365 (Fig. 2A) and maps exclusively to the deep inactive State 4 (intrinsic MAC = 0.361; Fig. 2B). Despite 100% R-spine intact, this baseline is 77% αC-Out and only 28% Active BLAminus, its activation-loop backbone sitting predominantly in the non-canonical BLBminus conformation (63%; Fig. 2D). The DFG motif already scores spatially in here and in every other condition, so that readout does not discriminate along this path. Simulated CCNB1 docking rotates the αC-helix from 77% “Out” to 88% “In”, resolves the activation loop into a near-uniform Active BLAminus architecture (28% to 98%), and lowers global rigidity to MAC = 0.217 (p.adj = 4.5×10^-6^; Fig. 2A). This trans-allosteric decoupling moves the complex out of State 4 into a two-state intermediate of significantly different intrinsic rigidity (p.adj = 0.009) as State 8 (50% of models, intrinsic MAC = 0.183) and State 7 (39%, 0.264). This decoupling is a change in coupling within the ensemble rather than in its composition. Recomputing MAC within each occupied state reproduces 91% of the pooled change (−0.135 of the observed −0.149), the transition retains 97% of its amplitude on a metric panel matched between the two conditions, and the single-state apo condition yields no composition contribution at all, a null the procedure recovers exactly (compare Note 10).

**Note 3. ATP fluidizes the pre-phosphorylation complex.** Introducing ATP to the CDK1– CCNB1 heterodimer (CDK1-Holo + CCNB1) lowers the global condition MAC from 0.217 to 0.123 (p.adj = 5.9×10^-8^) and drives the ensemble out of the two-state +CCNB1 intermediate into State 9, which takes 92% of the condition (Fig. 2C). State 9 carries the numerically lowest intrinsic MAC in the landscape (0.147), although it is not statistically separable from the other members of the relaxed tier (p.adj ≥ 0.24). Its distinguishing feature is not the activation-loop rotamer, which is already fully canonical at this step (100% Active BLAminus, 100% αC-In), but assembly of the internal spine bridge, which contracts from 7.9 Å in the two +CCNB1 states to 4.0 Å. ATP therefore maximizes network decoupling and completes the internal spine, leaving the final tensioning step to phosphorylation. This transition is the most panel-sensitive of the CDK1 series. Recomputed on the metrics shared by both conditions it retains 69% of its amplitude (+0.050 to +0.035 on 16 metrics), against 96 to 97% for every other transition in Fig. 2A.

**Note 4. pT161 acts as a mechanical lever.** The pT161 Holo complex escapes State 9 into the mature State 1 ensemble (93% of models; 98% Active BLAminus, 98% αC-In, 100% DFG-in). Pairwise differential driver analysis (unphosphorylated Holo intermediate to active pT161) reveals a seesaw: compression of the K105-N99 regulatory toggle (Cohen’s d = −1.20, p.adj = 4.0×10^-7^) with reciprocal expansion of the deep Y156-N99 scaffold (d = +2.09, p.adj = 3.2×10^-35^) and the M118-M120 hydrophobic shell (d = +2.56, p.adj = 9.6×10^-36^). Differential correlation analysis shows pT161 synchronizes the catalytic HRD motif with the P-loop (HRD_ATP : PLoop_ATP, ρ = 0.09 to 0.66; p.adj = 1.5×10^-5^) and couples DFG mechanics (DFG_Mg : DFG_ATP, ρ = −0.09 to 0.40; p.adj = 2.3×10^-3^). Global condition MAC rises to 0.177, re-imposing localized tension on an ensemble that was already canonically BLAminus before phosphorylation.

**Note 5. Inhibitory pT14/pY15 warp the active-site geometry.** The active (pT161) and inhibited (pT14/pY15/pT161) complexes co-populate the mature State 1 ensemble (93% and 92% of models respectively; Fig 2C) and are indistinguishable in global rigidity (p.adj = 0.24; Fig. 2A), with almost no differential allosteric rewiring. Differential distance analysis localizes inhibition to the nucleotide pocket. The DFG motif is hyper-compressed against Mg2+ (DFG_Mg_Dist, d = −0.99, p.adj = 1.1×10^-9^) while the P-loop is displaced away from ATP (PLoop_ATP_Dist, d = +0.81, p.adj = 9.5×10^-8^). In the triple-phosphorylated ensemble the pT14 phosphate oxygens fall within 4 Å of the ATP β/γ-phosphates in 26% of models (median 4.7 Å) versus 5% for the unphosphorylated threonine of the active complex; the Mg^2+^ ion stays canonically coordinated by the ATP phosphates and the DFG aspartate (≈2 Å in every model), and pY15 projects away from the pocket (median 10 Å to ATP). Full distance distributions and statistics are given in the S1 Fig legend.

**Note 6. SRC docking primes CSK.** The Truth Overwrite algorithm (AlloQuant Module 1) detects models in which AlphaFold3 re-allocates ATP away from the seeded apo/holo design. Comparing the unphosphorylated heterodimer (csk-wtcat-apo / src-wtcat-holo) to the primed counterpart (csk-wtcat-apo / src-wtcat-py159-holo) revealed a significant shift in ligand allocation (FDR-adjusted *p* = 0.0054; S3 Fig). Re-allocation was also seen for csk-holo / src-apo versus csk-apo / src-holo (*p* = 1.5×10^-12^). Per-condition overwrite rates are given in the S3 Fig legend.

**Note 7. SRC autophosphorylation redistributes the activator among conformational states.** SRC is 96 to 100% Active (BLAminus, αC-In) in every biological condition (S4A Fig), and in every metastable state (S4C Fig). Its global MAC is bulk-flat across those conditions (0.124 apo/apo; 0.120 csk-holo/src-apo; 0.129 both-holo; 0.124 primed; S3 Table). A higher SRC MAC (0.178) appears only in the excluded csk-apo/src-holo design (87% ATP ejection in the Truth Overwrite analysis; S3 Fig), which we do not interpret. Y419 (residue 159 in the isolated kinase domain) redistributes nucleotide-loaded SRC from a single metastable state (State 4, 86%) into State 8 (56%) and State 7 (34%; Fig. 5A) with essentially unchanged bulk rigidity. Both members of the primed pair are more internally coupled than the flexible baseline State 1 (intrinsic MAC 0.146 for State 7 and 0.127 for State 8, against 0.104; p.adj = 0.0016 and 0.010), but they are not separable from each other (p.adj = 0.25), so neither is treated here as the more rigid member. The relevant comparison for the effect of the mark is against State 4, the state the mark depletes. Against that baseline both primed states differ significantly in 19 of 21 metrics, but the differences are small and consistently in the direction of opening rather than compaction. The catalytic cleft widens (Cleft_Gape_Dist 15.66 to 16.32 Å into State 8, Cohen’s *d* = +1.38) and the nucleotide contacts loosen (PLoop_ATP_Dist 3.23 to 3.84 Å; HRD_ATP_Dist 5.28 to 5.88 Å; DFG_ATP_Dist 2.46 to 2.80 Å). The internal spine bridge, which ATP has already closed to 4.28 Å in State 4, is by contrast among the least mobile terms, moving +0.07 Å into State 7 and −0.14 Å into State 8. The spine closure is therefore a consequence of nucleotide loading and not of the activation-loop mark. Every remaining significant driver moves by 0.23 Å or less, including the regulatory salt bridge (SB_Dist +0.26 Å) and the shell and R-spine contacts. The largest apparent movements in this contrast are not contacts and are treated in Note 8.

**Note 8. The docking interface does not detectably change on SRC priming.** Two quantities in the differential driver analysis are labelled as interface distances and measure different things. On the SRC donor chain, ‘Interface_N_Lobe_Rec_Dist’ is a minimum distance between the CSK C-lobe and the SRC N-lobe, and runs 10.09 to 11.44 Å between SRC States 4 and 8 (*d* = +1.85). Those two blocks lie 10 to 11 Å apart and are therefore never in contact, past the pipeline’s own 8 Å contact threshold. The engaged face, meaning the residue pairs actually in van der Waals contact, moves 2.56 to 2.64 Å over the same contrast (*d* = +1.71), that is +0.08 Å. The large standardised effects reflect the tightness of the within-state distributions rather than the size of the displacement, as the caveat in Note 7 already notes. Measured cutoff-free across all 33 interface pairs, the face is unchanged. The mean displacement is +0.07 Å, only four pairs move by more than 0.3 Å, and 30 of the 31 interface residues retain the same nearest partner across the interface in both states. Apparent switches in contact *frequency* are threshold effects. The four pairs whose frequency changes most all sit within 0.25 Å of the 4.5 Å cutoff, and F197– Y254 loses two thirds of its contact frequency on a 0.061 Å shift. These displacements fall below AlphaFold3’s own coordinate uncertainty at the same positions (Methods). The median predicted aligned error over the interface pairs is 2.13 Å against a median displacement of 0.099 Å, so every measured shift is at most 27.8% of its own pair’s error and typically 3.9%. Confidence at the interface is additionally state-dependent, in the direction that would manufacture an apparent relaxation. Mean interface error rises from 2.81 to 3.40 Å between States 4 and 8 (*p* = 1.2 × 10^-17^) while both kinase cores become *more* confidently predicted. We therefore report the interface as unchanged within resolution, rather than as measured to be rigid.

**Note 9. SRC priming commits a subpopulation of CSK to the active state.** Unsupervised GMM shows CSK progression is strongly SRC-dependent. Unphosphorylated baseline (csk-apo/src-apo): CSK is deeply trapped in inactive State 1 (73% of models), 0% αC-In, 87% BLBplus / 13% BLAminus. ATP without the SRC mark (csk-holo/src-apo): shifts into States 7 and 8 (49% and 32%) but stays locked inactive (5% αC-In, 17% BLAminus, 83% BLBplus). SRC pY419-primed holo: accesses the fully active State 3 (Fig. 4E; members 97.8% αC-In, 97.8% BLAminus, 100% DFG-in), raising ensemble Active BLAminus from 17% to 47% and αC-In from 5% to 27% (S4B Fig). State 3 nonetheless accounts for only 28% of the primed ensemble, and the ensemble-level rise is therefore a change in state occupancy rather than a graded shift across models: αC-In is confined to State 3 (States 1, 2, 5, 7 and 8 contain no αC-In model), and the BLAminus population outside State 3 comes from States 9 and 4, which adopt the active backbone with the αC-helix extruded (S4 Table; S4D Fig). Pairwise driver analysis (State 7 to State 3) shows a coordinated active-site rearrangement at constant bulk rigidity (intrinsic MAC 0.121 to 0.127, n.s.). The regulatory salt bridge closes (SB_Dist, Cohen’s d = −7.17, a 14.3 to 3.3 Å transition), the deep Y156-N99 scaffold and αE floor compact (Y156_N99_Dist d = −4.99; I150_HRD_Dist d = −4.40), the substrate cleft opens (Cleft_Gape_Dist d = +1.48) and the HRD–nucleotide spacing widens (HRD_ATP_Dist d = +1.66). The K105 network reorganizes rather than collapses (K105-E121 d = −1.67; K105-E107 d = +1.74; K105-N99 *d* = +1.13). The docking interface itself is unchanged within resolution (Note 8).

**Note 10. The pooled CSK rigidity rise follows from ensemble composition.** Global MAC is one correlation network per condition (Methods), so pooling models drawn from different metastable states induces correlation among every metric that separates those states. In the CSK receiver the two effects coincide. Pooled MAC rises from 0.148 in the both-ATP baseline (csk-holo/src-holo) to 0.240 on complexing with primed SRC (csk-holo/src-py159-holo), a change of +0.092 (S3 Table), and priming is also the transition that redistributes the CSK ensemble, State 3 rising from 3% to 28% of models (S4 Table). Recomputing the same statistic within each occupied state, on that condition’s own metric panel so that the variable set stays matched, gives 0.198 (State 7, 45% of the condition) and 0.161 (State 8, 40%) for the baseline, against 0.166 (State 5, 42%), 0.168 (State 3, 28%) and 0.236 (State 9, 15%) for the primed complex, with occupancy-weighted values of 0.181 and 0.179. All four conditions, their small-sample floors and their size-matched nulls are tabulated in S7 Table. Composition alone reproduces the observed +0.092 to within 0.002, whereas redistribution toward more strongly coupled states accounts for +0.006 of it. Intrinsic MAC reaches the same conclusion independently, the activated State 3 being no more strongly coupled than the State 7 it displaces (0.127 against 0.121, n.s.; Note 9). Because no CSK state is well populated under both conditions (State 7, 45 models unprimed against 4 primed; State 5, 2 against 42), the recomputation bounds a within-state coupling change rather than excluding it.

**Note 11. Nucleotide acts locally; reciprocity is conformational, not bulk-mechanical.** SRC global rigidity is unchanged between CSK-apo and CSK-ATP conditions (MAC ≈ 0.12 across the biological conditions); nucleotide tunes specific contacts (the β3-Lys–αC-Glu network and the regulatory spine) without reorganizing bulk coupling. Within the pY419-primed complex, SRC and CSK metastable states are significantly associated per structure (Cramér’s V = 0.44, permutation p < 0.001), the active SRC States 8 and 7 co-occurring with the more active CSK States 5, 3 and 9 in the same models. This coupling is absent from the unprimed complex (V = 0.26, p = 0.23, n.s.) and is therefore gated by the SRC activation-loop mark (Fig. 5B).

**Note 12. Homologous regulatory spines are coupled within individual complexes.** Within-condition partial Spearman correlations (795 paired models across seven conditions; ipTM and interface PAE partialled out, Benjamini–Hochberg-adjusted across the full metric grid) are dominated by the regulatory-spine module: the K105–E121 αC-loop network (ρ = 0.28, FDR = 1.3×10^-13^), the regulatory R-spine (V104–RS2; ρ = 0.25, FDR = 1.2×10^-10^), the HRD backbone (D220; ρ = 0.22, FDR = 1.0×10^-7^), the regulatory shell (M118–M120; ρ = 0.21, FDR = 6.2×10^-7^), the internal spine bridge (ρ = 0.18, FDR = 2.7×10^-5^) and the K105–N99 contact (ρ = 0.14, FDR = 2.6×10^-3^). The catalytic and active-site elements are uncoupled: the regulatory salt bridge (ρ = 0.02), the αE-helix floor (0.06), the substrate cleft (0.08), the deep Y156 scaffold (−0.02) and the HRD–nucleotide spacing (0.02) are all non-significant after adjustment (FDR ≥ 0.22; Figs. 4C and 6). Two further elements outside the spine module do survive adjustment: the DFG-Asp backbone ψ dihedral (ρ = 0.21, FDR = 1.8×10^-7^) and the P-loop–nucleotide spacing (ρ = 0.22, FDR = 0.025). The cross-interface coupling is therefore concentrated in, but not strictly confined to, the regulatory spine.

**Note 13. Network rigidity differences are not a prediction-confidence artifact.** MAC is correlation-based, so condition-to-condition differences could in principle reflect AlphaFold3 confidence (better-predicted ensembles being more internally self-consistent) rather than mechanical coupling. Recomputing every network edge as a partial Spearman correlation conditioned on per-model confidence (Methods; S6 Fig) leaves the CSK–SRC series essentially unchanged. With ipTM and interface PAE partialled out, MAC shifts by ≤ 0.006 across the seven CSK conditions and ≤ 0.012 across the SRC series, and the rise in CSK global MAC on SRC priming reads 0.147 to 0.246 after partialling versus 0.148 to 0.240 before (FDR = 6.1×10^-9^; S3 Table), with SRC remaining bulk-flat. Nineteen contrasts were significant before partialling (13 CSK, 6 SRC) and twenty afterwards (14 and 6), and no contrast that remains significant reverses direction. For the CDK1 series only pTM is available as a covariate, because the apo conditions are monomeric and have no interface. There the four nucleotide-bound conditions move by ≤ 0.005, whereas the two nucleotide-free ensembles prove appreciably confidence-associated. The trapped apo monomer falls from 0.365 to 0.231 and CDK1–CCNB1 from 0.217 to 0.171, so the CCNB1 decoupling loses roughly 60% of its amplitude (ΔMAC 0.148 to 0.060) even though its direction and significance hold. All four transitions annotated in Fig. 2A retain their verdicts: apo to +CCNB1 (FDR = 0.017), +CCNB1 to +CCNB1+ATP (2.8×10^-3^), +CCNB1+ATP to triple-P (0.017), and triple-P versus active pT161 (not significant, 0.14). Four contrasts fall below significance after partialling: three in the CDK1 series, two of them involving the nucleotide-free CDK1–CCNB1 ensemble, one of which (+CCNB1 versus active pT161) also inverts sign as it dissolves. The third, cyclin-free CDK1+ATP versus triple-P (FDR 0.016 to 0.063), was marginal to begin with, plus one already-marginal CSK contrast (FDR 0.038 to 0.057). Conversely, two CSK contrasts that were marginal beforehand cross the threshold after partialling. None of these six is cited in this work; all are listed in S6 Table, and per-condition raw and partial MAC values with the covariates used per group are given in S5 Table. This control addresses prediction confidence only. It does not test whether a difference in global MAC reflects a condition’s metastable-state composition, which is assessed separately in Note 10. The CSK rise in fact reads marginally larger after partialling, so it is the composition control and not this one that accounts for it.

**Note 14. Intrinsic MAC for primed-SRC states is conservative with respect to the ATP-leakage condition.** The csk-apo/src-py159-holo condition yielded only 33 models after Truth Overwrite filtering (87% of designed models had ATP migrate to CSK). Of these, 26 were assigned to the primed-SRC states (State 7: n=8; State 8: n=18). These models are excluded from the biological comparisons of Fig. 5 but are included in the intrinsic-MAC computation, which pools all models of a state regardless of condition. Removing them reduced State 7 from 49 to 41 members and State 8 from 75 to 57, and the full pairwise Wilcoxon analysis (Note 13 procedure) was repeated. Intrinsic MAC rose for both states (State 7: 0.146→0.158; State 8: 0.127→0.137); the comparisons State 1 vs 7 and State 5 vs 7 strengthened from p.adj ** to ***, and no significant comparison lost significance. The leaky models suppress rather than amplify the primed-SRC rigidity signal; the full-pool analysis is therefore conservative.

**Note 15. CCNB1 is rigid, by a measure that does not require kinase landmarks.** The manuscript describes CCNB1 as a rigid cofactor, and MAC cannot support that. Global and intrinsic MAC are built from inter-landmark distances assigned by the kinase profile HMMs of Module 1, so neither is defined for a cyclin. CCNB1 contributes no landmark, and the two cofactor distances in the master table (CoFactor_aC_Dist and CoFactor_ActLoop_Dist) are measured from CDK1 to CCNB1 rather than within CCNB1. We therefore assessed CCNB1 with a quantity that needs no landmarks, the per-residue Cα RMSF across each condition’s 100 models after chain-local superposition (Methods), which reports how reproducibly AlphaFold3 places the chain’s own fold. CCNB1 is the less variable chain in every complex, and its value barely moves: median RMSF 0.14, 0.12, 0.12 and 0.13 Å in +CCNB1, +CCNB1+ATP, pT161 and triple-P respectively, against 0.39, 0.23, 0.14 and 0.17 Å for CDK1 in the same models (S9 Table). CCNB1 is thus indifferent to nucleotide and to phosphorylation state, whereas CDK1 varies three-fold across the same series. Mean pLDDT is also higher for CCNB1 in every condition (95.1 to 95.8 against 90.8 to 95.6), so the difference is not a confidence artifact in the direction that would matter. CCNB1’s rigidity is in the same sense that the SRC activator is called rigid in S5 Fig, a low and uniform Cα RMSF across the ensemble, with the difference that the superposition here is chain-local and so excludes motion of one subunit against the other. The comparison is not, however, one of a soft kinase against a stiff cofactor. CDK1’s excess variability is localized rather than global. In the +CCNB1 complex the residues above 1.0 Å are confined to the Gly-rich P-loop and its inhibitory phosphosites (12 to 15), the β3-αC region (36 to 51), the activation loop carrying T161 (150 to 166, peaking at 6.3 Å) and the C-terminal tail (291 to 297), which is 45 of 297 residues. Once the activating mark is present, CDK1’s median falls to 0.14 Å against CCNB1’s 0.12 Å and the two chains are effectively indistinguishable. CCNB1 therefore presents the same structure in every condition while CDK1’s residual variability sits on the regulatory elements this study is about, which is the structural content of describing CCNB1 as a constitutive scaffold. One limit applies beyond the distinction between MAC and RMSF set out in Methods. RMSF here is dispersion across independently seeded predictions rather than a thermodynamic fluctuation, so a low value means AlphaFold3 places the fold consistently. It does not mean the protein is mechanically stiff.

**Note 16. The doubly phosphorylated SRC state, and the conditions under which it has been observed.** The main text treats the Y530 mark as one whose consequence is deferred rather than immediate. Two independent studies support that reading, and each carries conditions worth stating. In a reconstituted system of purified enzymes, Src autophosphorylated in its activation loop is still phosphorylated by Csk at the C-terminal tyrosine, but is no longer inactivated by it, and the doubly phosphorylated kinase is hyperactive. Treating that Src with PTP1B, which under the conditions used preferentially removes the activation-loop phosphate, restores Csk sensitivity. A Yes mutant unable to autophosphorylate is inactivated by sub-stoichiometric Csk [4]. In cells, Lck activated by hydrogen peroxide carries phosphate at both the activation loop and the C-terminal tyrosine while remaining active, so tail dephosphorylation is not a prerequisite for activation [5]. Both systems depart from the unperturbed cell. The first lacks membrane, scaffolds, SH2 ligands and endogenous phosphatases, and its enzyme ratios are set by design. The second disables tyrosine phosphatases broadly, which is how the doubly phosphorylated form accumulates. What the two establish jointly is narrower than either claims alone, but it is robust to those caveats. The doubly phosphorylated SFK is chemically accessible and catalytically active, the two marks are compatible, and CSK will act on an already-primed SRC. The two perturbations are also worth comparing directly. One supplies phosphatase activity and one removes it, and both implicate a phosphatase as the step that decides the outcome. Neither study identifies the cellular phosphatase. PTP1B served in the first as a broad-specificity tool, not as a candidate for the physiological step. Purified PTP1B is promiscuous in solution, and in cells it acts on SRC at the C-terminal tyrosine, activating SRC rather than sensitizing it to CSK [6]. Candidate enzymes have since been identified, and one prediction of the latent-mark reading has been partly borne out. Sun et al. noted that colorectal tumours can carry active SRC alongside abundant, fully active CSK, proposed that persistent autophosphorylation would explain it, and called for the phosphorylation state of both sites to be examined in tumours. PTEN was subsequently shown to dephosphorylate SRC at the activation-loop site, with loss of PTEN leaving that site phosphorylated in trastuzumab-resistant breast cancer [7]. CSK was not measured in that work, so the two accounts are joined here by inference rather than by either study.

### Supplementary Figures

**S1 Fig.**
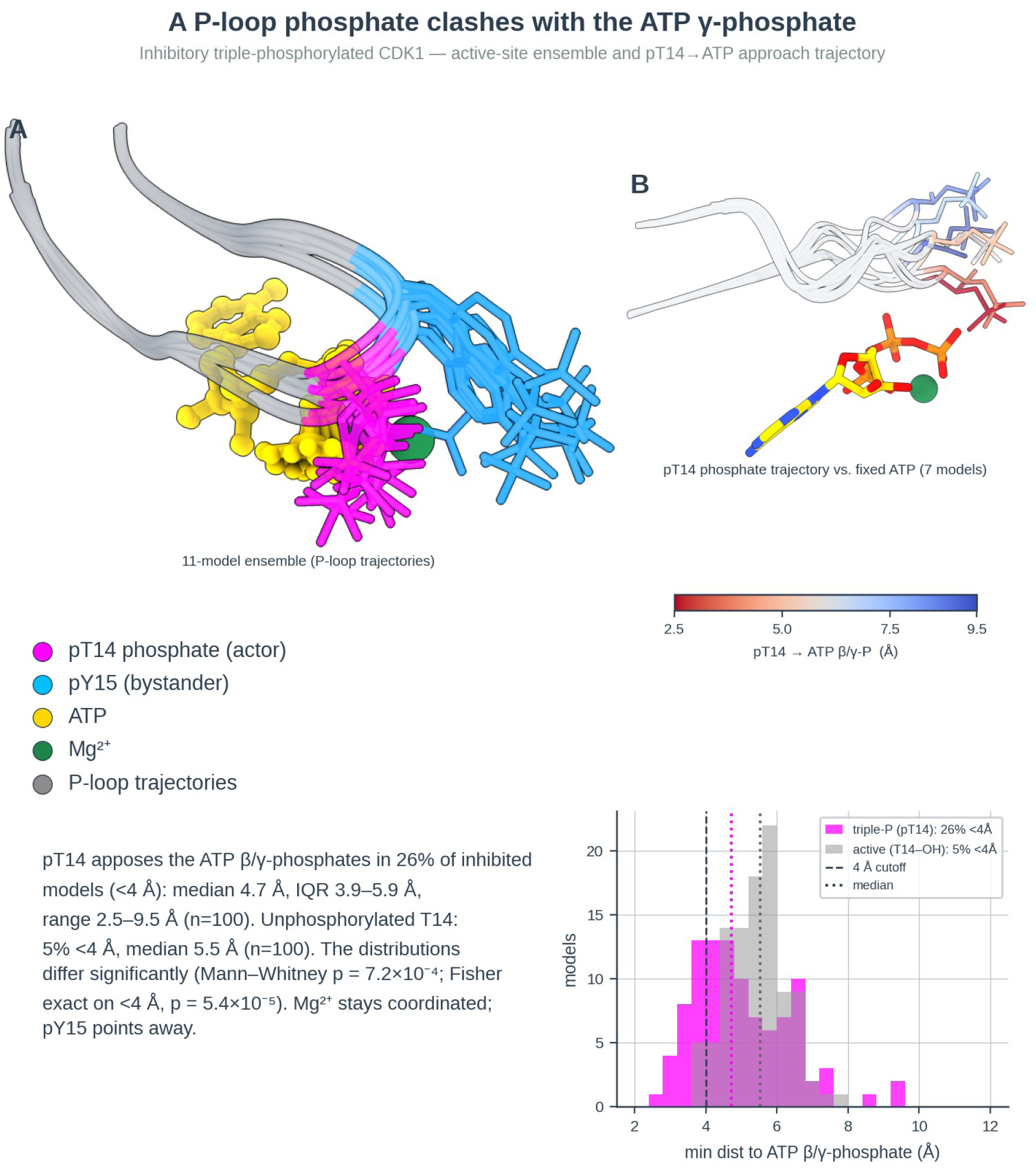
The P-loop pT14 phosphate clashes with the ATP γ-phosphate in inhibited CDK1. *(Top)* Active-site close-up of the triple-phosphorylated CDK1 ensemble (11 models superimposed on ATP). The pT14 phosphate (magenta, “actor”) projects from the glycine-rich P-loop directly onto the ATP β/γ-phosphates, whereas the second inhibitory mark pY15 (deep-sky-blue, “bystander”) points away from the pocket; ATP is gold, the catalytic Mg^2+^ is green, and the P-loop backbone is gray. *(Bottom right)* Distribution of the minimum distance from the T14 side chain to the ATP β/γ-phosphate across all models: in the triple-P ensemble the phosphate lies within 4 Å in 26% of models (median 4.7 Å, IQR 3.9–5.9 Å, range 2.5–9.5 Å, *n* = 100), versus 5% for the unphosphorylated T14 hydroxyl of the active pT161 complex (median 5.5 Å, IQR 4.9–6.0 Å, *n* = 100). The two distributions differ significantly (Mann–Whitney *U* test, *p* = 7.2×10^-4^; Fisher exact test on the <4 Å proportion, *p* = 5.4×10^-5^, odds ratio 6.7). The dashed line marks the 4 Å contact cutoff; dotted lines mark the per-group medians. The Mg^2+^ ion remains canonically coordinated by the ATP phosphates and the DFG-Asp (≈2 Å in every model), indicating that inhibition operates through direct phosphate–phosphate electrostatic repulsion at the transferable γ-phosphate rather than metal sequestration.

**S2 Fig.**
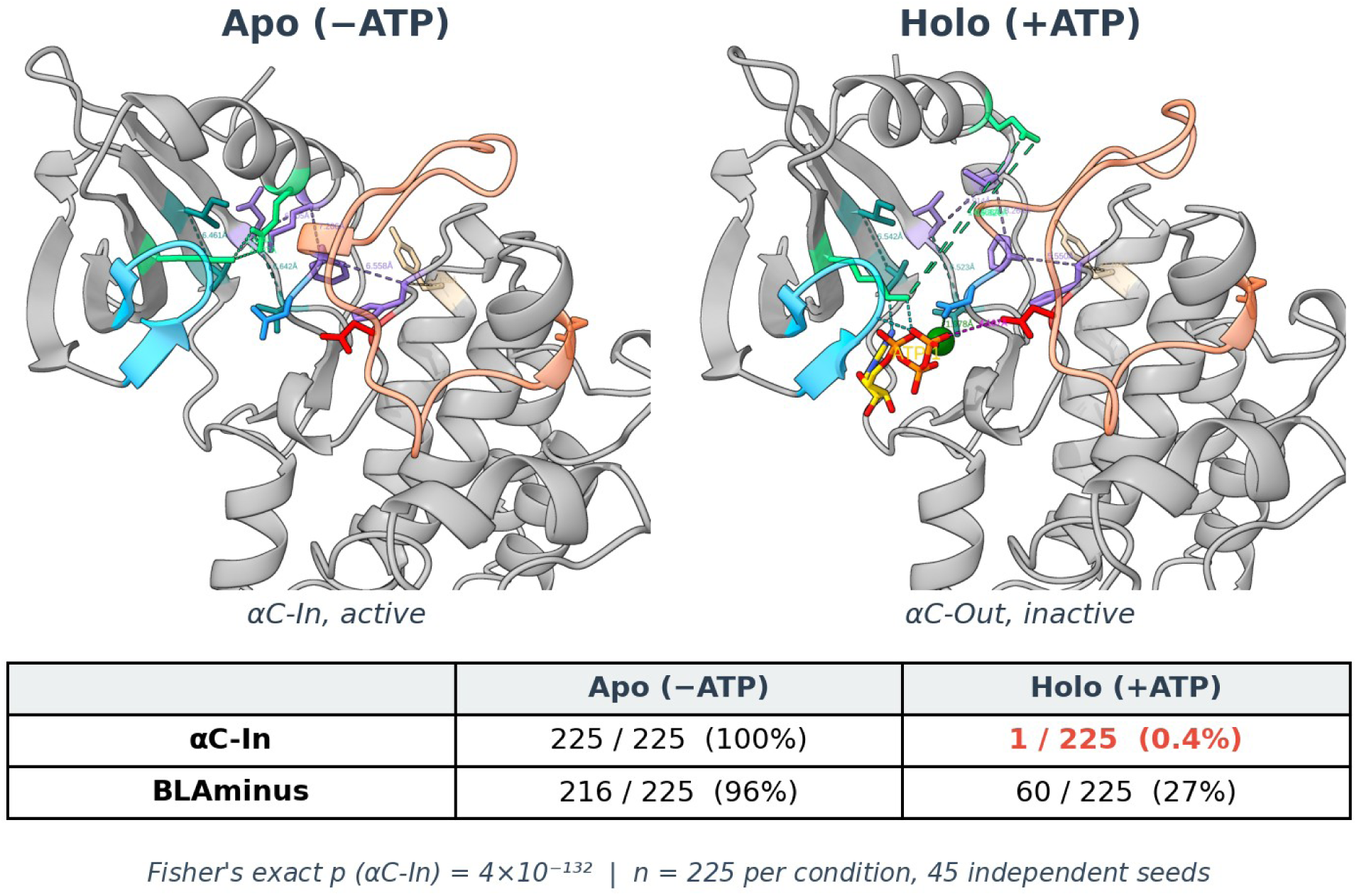
Nucleotide loading locks monomeric CSK out of the active αC-In conformation. Representative AlphaFold3 models (full-MSA, per-condition centroids; element coloring as in Fig. 3) of unbound apo (left) and ATP-loaded holo CSK (right). Apo: αC-In, β3-Lys–αC-Glu salt bridge formed (mean 2.8 Å, 95% CI 2.78–2.84 Å; spring green). Holo: αC-Out, salt bridge broken (mean 16.3 Å, 95% CI 16.1–16.6 Å). Table: ensemble statistics (45 seeds × 5 models per condition, n = 225; Fisher’s exact OR = ∞, p = 4 × 10⁻¹³²).

**S3 Fig.**
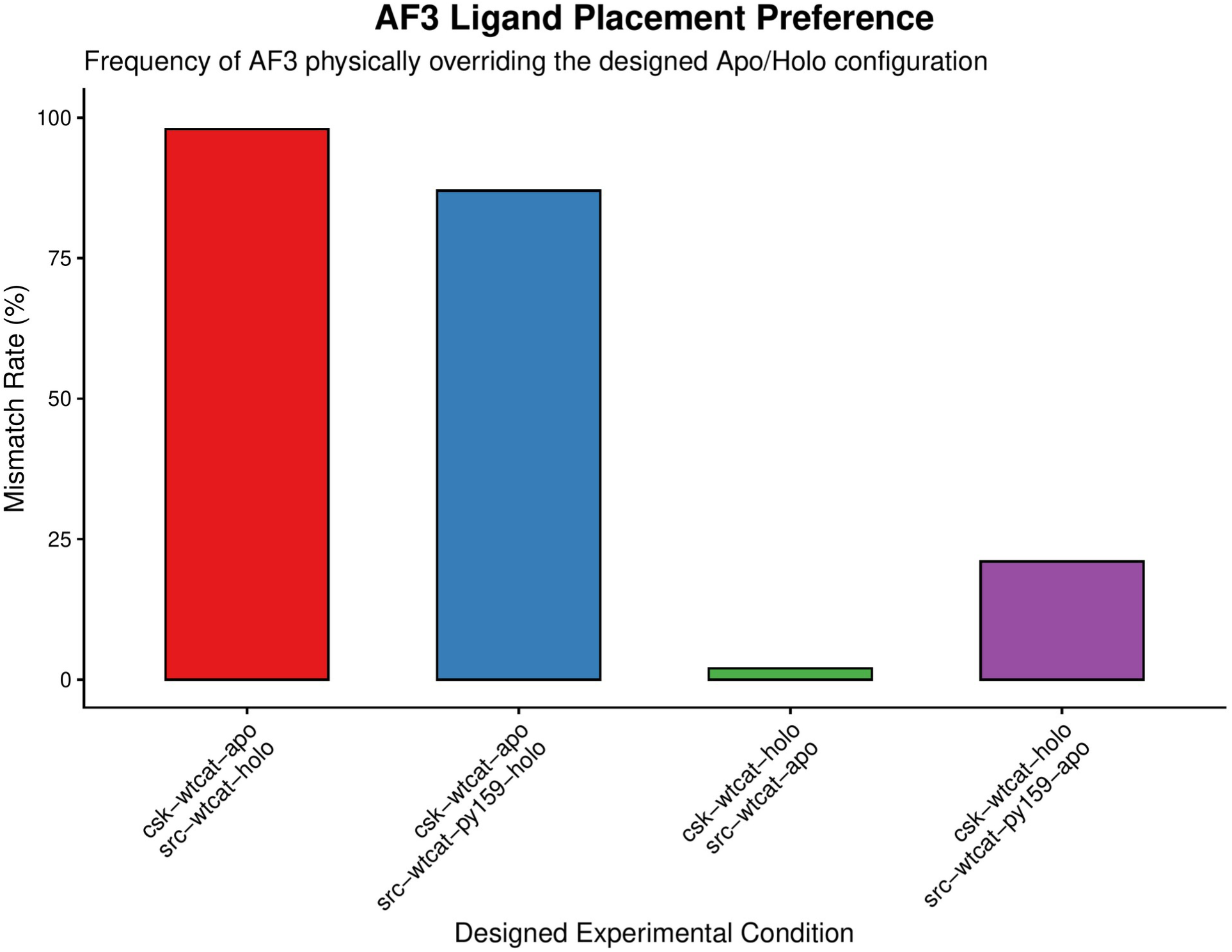
AlphaFold 3 overrides the designed ligand state, asymmetrically favoring the SRC-primed complex. For each competitive asymmetric CSK–SRC design (one subunit seeded apo, the other holo), the “Truth Overwrite” discrepancy algorithm (AlloQuant Module 1) compares the designed apo/holo configuration to the physically realized one. A model is scored as overwritten when the final ATP geometry, judged by its proximity to both the catalytic (C-)spine and the HRD-motif aspartate, no longer matches the seeded assignment. Bars give the per-condition overwrite (mismatch) rate (% of models; *n* = 100 per condition). AF3 overwhelmingly re-allocates ATP into the SRC-holo–bearing complexes: 98% for csk-apo/src-holo and 87% for csk-apo/src-py159-holo, versus only 2% for the reciprocal csk-holo/src-apo and 21% for csk-holo/src-py159-apo. Symmetric designs (apo/apo, holo/holo) showed no overwrites and are omitted. The re-allocation is significant both between the two SRC-holo complexes (± pY419 priming; Fisher exact *p* = 0.0054) and between the reciprocal csk-holo/src-apo and csk-apo/src-holo designs (Fisher exact *p* = 1.5×10^-12^), consistent with docking of an active SRC lowering the barrier to ATP coordination in CSK, an inference from the model’s placement preference rather than a measured energy.

**S4 Fig.**
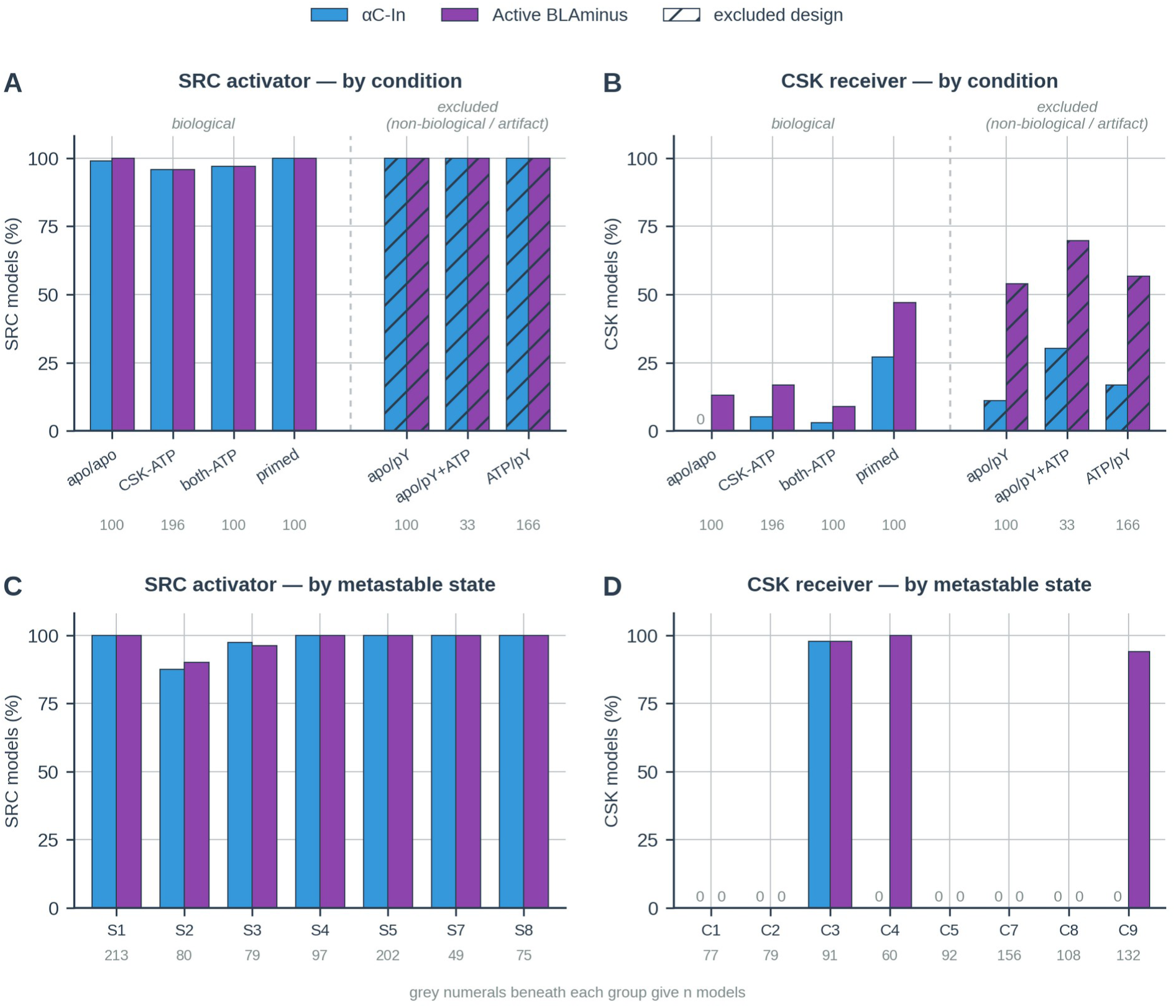
Categorical activation status is invariant in the SRC activator and all-or-none in the CSK receiver. Percentage of models scored αC-In and Active BLAminus, across all 795 CSK–SRC models. **(A, B)** By experimental condition: the four biological conditions in the order of S5 Table, then the three excluded designs, hatched and set apart: apo/pY and ATP/pY carry pY419 on an otherwise apo SRC and are non-biological, whereas apo/pY+ATP is an ATP-placement artifact (87% overwritten; Note 6). Designs are labelled CSK state / SRC state, where pY denotes the SRC activation-loop phosphomark pY419, as in S5 Table. SRC is 96– 100% αC-In and Active BLAminus in every design, biological or not, consistent with a domain that arrives active rather than switching into activity (A), whereas CSK climbs from 0% to 27% αC-In and 13% to 47% Active BLAminus along the biological progression (B). **(C, D)** By unsupervised GMM metastable state, pooled across all seven conditions. SRC states are labelled S1 to S8 and CSK states C1 to C9, following the contingency grid of Fig. 5B, so that one basin carries one name throughout. Every SRC state is 88–100% active on both calls, so the pY419-driven redistribution from State 4 into States 8 and 7 (Fig. 5A) moves SRC between categorically indistinguishable states: a conformational switch, not an activation switch (C). CSK is instead partitioned: αC-In is confined to State 3, the Active BLAminus population outside it comes entirely from States 9 and 4, and States 1, 2, 5, 7 and 8 contain neither (D) (S4 Table). The ensemble-level rise in (B) is therefore a change in state occupancy rather than a graded shift across models. DFG-in is omitted from all panels: it is 100% in every condition and every state and so discriminates nothing. Grey numerals give *n* models per group. Per-condition counts are unequal because each model is attributed to the condition it physically realized (Truth Overwrite; Note 6), which also leaves apo/pY+ATP with only 33 non-overwritten models, so its CSK values in (B) rest on a small, selected subset and are not comparable with the biological conditions. Zero values are labelled.

**S5 Fig.**
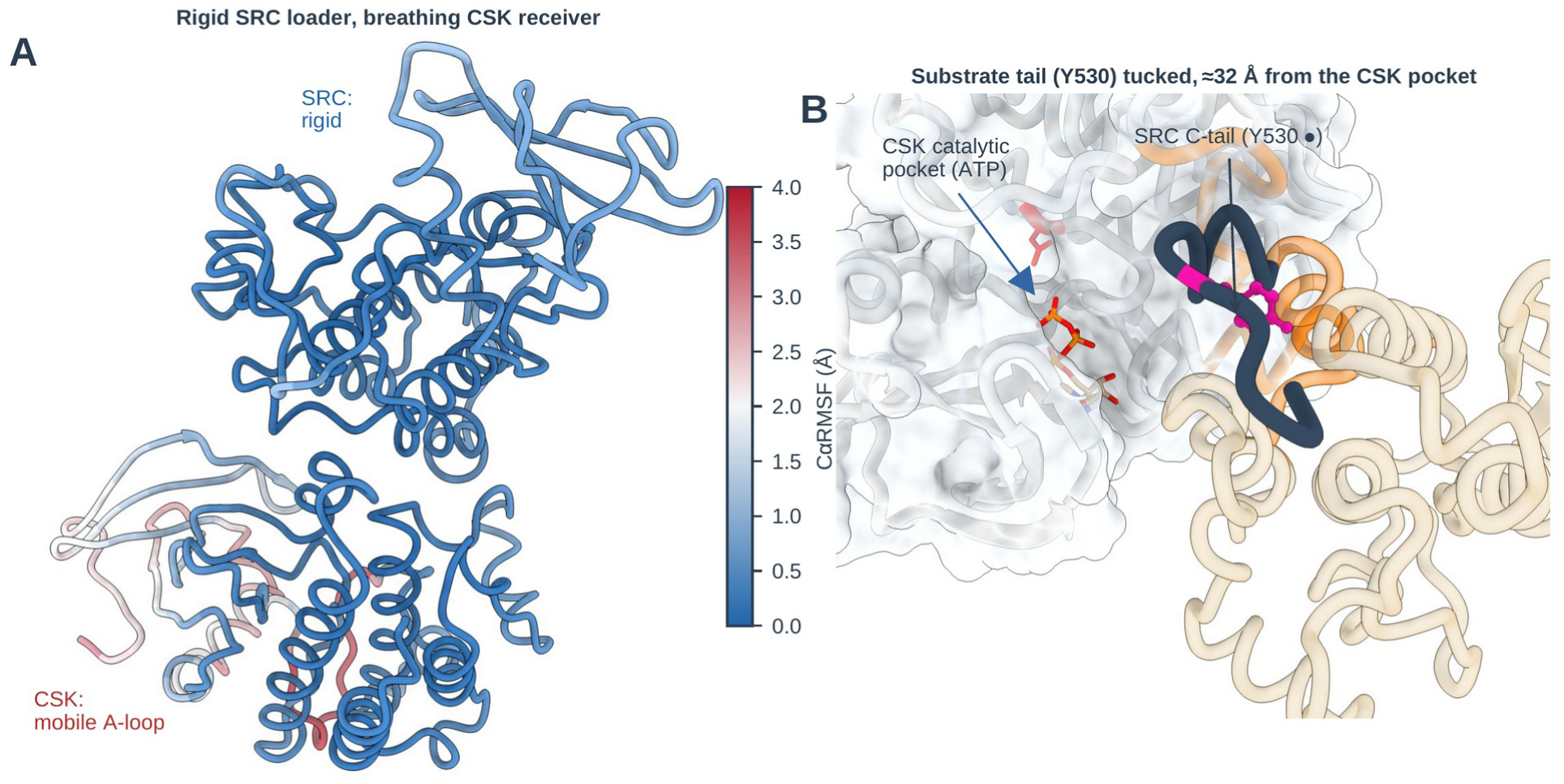
The SRC activator is rigid while the CSK receiver breathes, and the SRC substrate tail is sequestered in the docked complex. **(A)** Per-residue Cα root-mean-square fluctuation (RMSF) across the primed heterodimer ensemble (superposed on the rigid combined C-lobe core), mapped onto a representative complex (blue rigid to red mobile). SRC (activator) is uniformly rigid (mean RMSF 0.34 Å), whereas the CSK receiver breathes, with a markedly mobile activation loop (up to ∼3.7 Å). This is the structural counterpart of the activator/receiver asymmetry in Note 7 in S1 Text. **(B)** In the docked complex, the SRC C-terminal tail (residues 262–276; slate) folds back against SRC’s own C-lobe helices, placing the substrate tyrosine Y530 (magenta) ∼32 Å from the accessible CSK catalytic pocket (ATP, shown in the semi-transparent CSK surface). AlphaFold3 models this tucked tail with high confidence; productive presentation of Y530 to CSK would require the tail to release and extend (see Discussion). *n* = 100 models per condition.

**S6 Fig.**
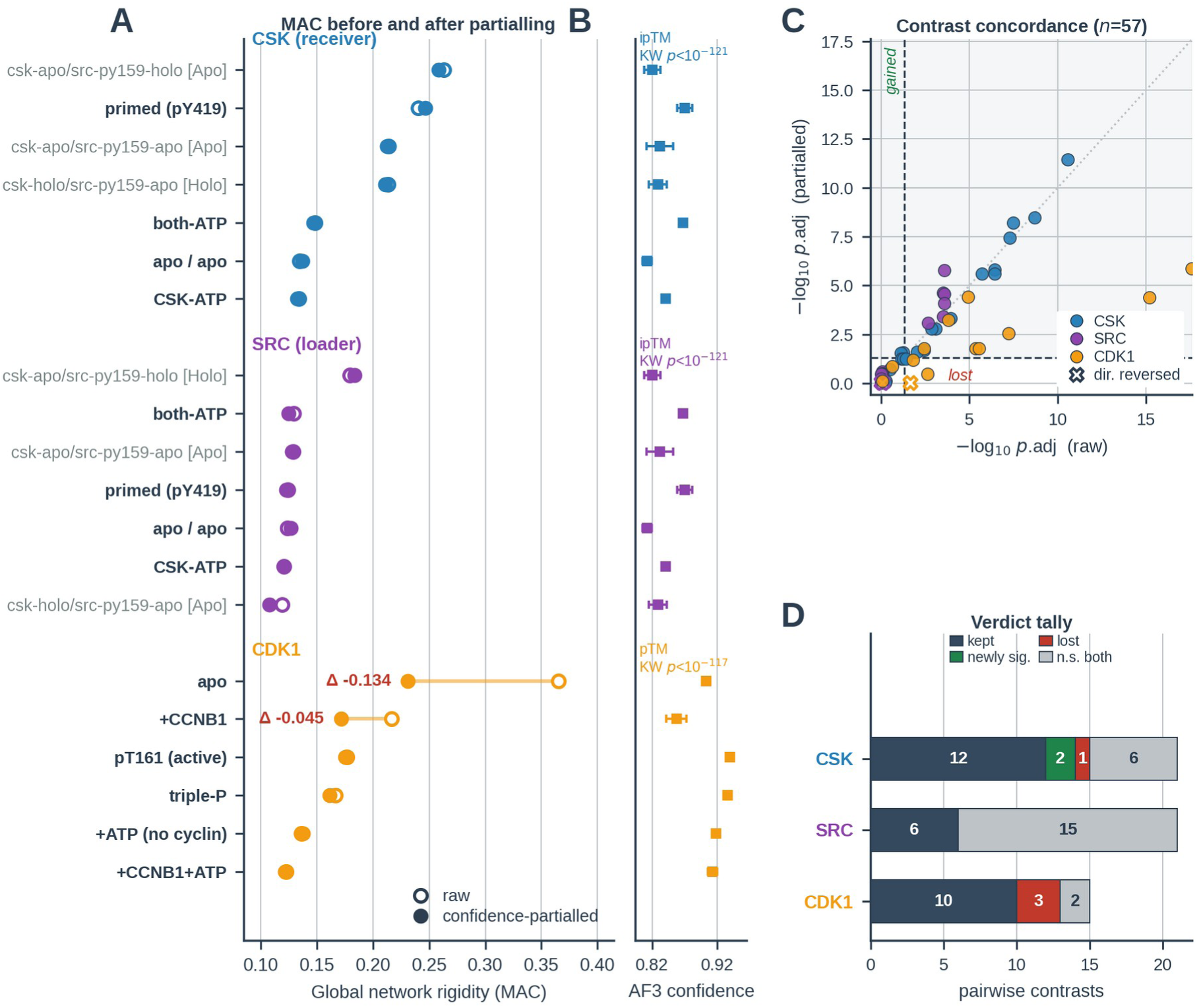
Network rigidity differences are not an AlphaFold3 confidence artifact. Companion to Note 13 and S5 and S6 Tables. **(A)** Global network rigidity (MAC) of every experimental condition before (open circles) and after (filled circles) recomputing each network edge as a partial Spearman correlation conditioned on per-model AlphaFold3 confidence (Methods). Rows are ordered by raw MAC within each series; conditions carried in the main text are labeled in bold with their Fig. 2A and Fig. 5 names, the remaining CSK–SRC designs in gray. Δ is printed where |ΔMAC| ≥ 0.02: only the two nucleotide-free CDK1 ensembles move appreciably, cutting the amplitude of the CCNB1 decoupling by roughly 60% (ΔMAC 0.148 to 0.060) without changing its direction or significance. **(B)** The confound is real: mean ± SD of the partialled covariate across the same conditions (interface ipTM for the CSK–SRC series; pTM for the CDK1 series, whose apo conditions are monomeric and have no interface). Confidence differs strongly between conditions (Kruskal–Wallis: ipTM *p* = 2.8×10^-122^, *n* = 795; pTM *p* = 9.2×10^-118^, *n* = 600), yet the MAC values in (A) are essentially unmoved. Mean interface PAE was partialled out alongside ipTM for the CSK–SRC series (S5 Table). **(C)** FDR-adjusted *p* for all 57 pairwise MAC contrasts, before versus after partialling (dashed lines, *p*.adj = 0.05; dotted line, identity; shaded band, significant after partialling). Crosses mark contrasts whose direction reverses; all fall in the non-significant region except one CDK1 contrast (nucleotide-free CDK1–CCNB1 versus active pT161, *p*.adj 0.01 to 0.99) that is not cited in this work. **(D)** Outcome of every contrast, by series: all 20 contrasts that were significant in the CSK–SRC series are kept and two more cross the threshold, whereas 10 of 13 CDK1 contrasts are kept, the three lost being listed in S6 Table. All four transitions annotated in Fig. 2A retain their verdicts.

**S7 Fig.**
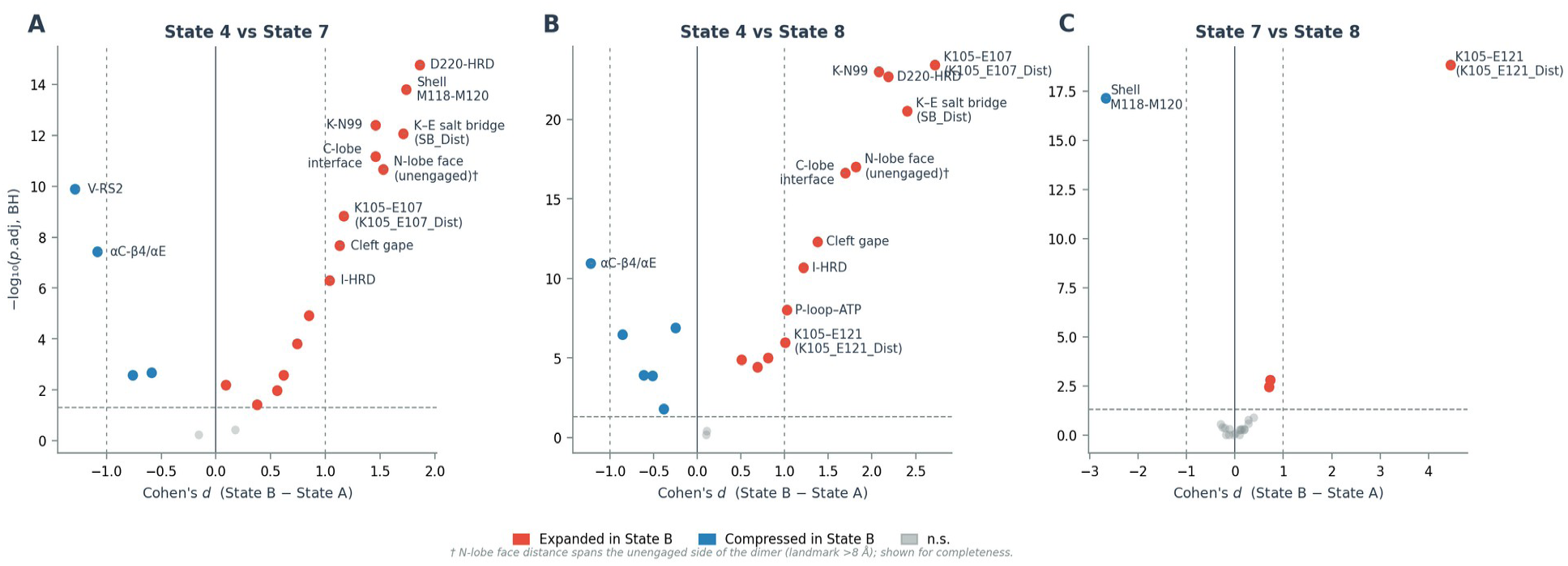
pY419 drives broad structural rearrangements distinguishing SRC State 4 from States 7 and 8, which are near-equivalent. Pairwise volcano plots from Phase 8 of the SRC metastable-state analysis (dimer ensemble 260718 rerun). **(A)** State 4 (ATP-loaded, unphosphorylated) vs State 7; **(B)** State 4 vs State 8; **(C)** State 7 vs State 8. x-axis: Cohen’s d (positive = expanded in State B); y-axis: −log_10_(p.adj, Wilcoxon + Benjamini–Hochberg). Dashed lines mark |d| = 1.0 and p.adj = 0.05; points labeled where |d| ≥ 1.0. States 7 and 8 each depart broadly from State 4 across the docking interface, K–E salt bridge, and catalytic cleft (A, B), while differing from each other in only 4 of 21 metrics (C); K105–E121 is the dominant divergence (d = 4.45). The internal spine bridge, closed to 4.3 Å in State 4 by ATP loading, changes by only 0.09 Å (4→7) and 0.26 Å (4→8). Metric column names follow the AlloQuant naming scheme (S1 Dataset). † N-lobe face distance spans the unengaged side of the dimer; see main text.

### Supplementary Tables

**S1 Table.** AlloQuant versus Kincore on the CSK–SRC crystal structure 3D7T. Every quantity both tools report, for both chains of PDB 3D7T [1], AlloQuant values are v7r3, the release used throughout this work. This comparison prompted two landmark-anchoring corrections, which v7r3 carries: the Dunbrack D1 coordinate is measured from the fourth residue after the αC-glutamate rather than from the glutamate itself, and the activation-loop C-terminal partner is the HRD arginine (HRD+1); a missing-density guard was also added so that landmarks separated by a disordered gap return not-available instead of a distance measured across it. Five calls moved into agreement as a result (CSK D1 and A-loop N-terminal, and SRC D1, DFG spatial class and A-loop C-terminal) and are therefore not flagged separately in the table. Two documented divergences remain, neither used anywhere in this study: AlloQuant scans an APE−6 to APE−9 window and falls back to resolved members, so it still calls the CSK A-loop C-terminal where Kincore declines; and its A-loop N-terminal test uses a 5.5 Å all-atom window against Kincore’s 3.6 Å backbone hydrogen bond, so it over-calls SRC NT-in. The dihedral cluster labels are not comparable by design: AlloQuant concatenates three independently thresholded Ramachandran regions, whereas Kincore snaps a seven-dimensional vector to canonical medoids and returns None beyond a cutoff, so AlloQuant can emit labels outside Kincore’s vocabulary. Both tools define Active as DFG-in with a BLAminus activation loop. Distances in Å, dihedrals in degrees.

| Chain | Quantity | AlloQuant v7r3 | Kincore | Verdict |
| --- | --- | --- | --- | --- |
| CSK | D1 (Glu4–Phe) | 4.85 | 4.85 | exact |
| CSK | D2 (Lys–Phe) | 13.73 | 13.73 | exact |
| CSK | β3K–αCE salt bridge | 3.50 | 3.50 | exact |
| CSK | DFG-Asp $\phi$, $\psi$ | –112.49,<br>120.17 | –112.49,<br>120.17 | exact |
| CSK | DFG spatial class | DFGin | DFGin | agree |
| CSK | αC orientation | In | Chelix-in | agree |
| CSK | Regulatory spine | Intact | Spine-in | agree |
| CSK | A-loop N-terminal | not available | None | agree |
| CSK | A-loop C-terminal | CTin | None | divergent |
| CSK | A-loop dihedral cluster | ABBminus | None | out of vocabulary |
| CSK | Active / non-active | non-active | Inactive | agree |
| SRC | D1 (Glu4–Phe) | 12.81 | 12.81 | exact |
| SRC | D2 (Lys–Phe) | 10.12 | 10.12 | exact |
| SRC | β3K–αCE salt bridge | 2.62 | 2.62 | exact |
| SRC | DFG-Asp $\phi$, $\psi$ | –79.66, 41.23 | –79.66, 41.23 | exact |
| SRC | DFG spatial class | DFGout | DFGout | agree |
| SRC | αC orientation | In | Chelix-in | agree |
| SRC | Regulatory spine | Broken | Spine-out | agree |
| SRC | A-loop N-terminal | NTin | NTout | divergent |
| SRC | A-loop C-terminal | CTin | CTin | agree |
| SRC | A-loop dihedral cluster | BABminus | None | out of vocabulary |
| SRC | Active / non-active | non-active | Inactive | agree |

**S2 Table.** Pairwise comparisons of per-state intrinsic network rigidity (CDK1–CCNB1). Each metastable state’s intrinsic MAC is the mean absolute Spearman correlation across its structural-metric network. State pairs were compared by Wilcoxon rank-sum test on the per-edge absolute correlations, with Benjamini–Hochberg adjustment across all 21 comparisons; rows are ordered by *p*.adj. These tests define the three intrinsic-rigidity tiers annotated in Fig. 2B: State 4 (trapped, inactive) is significantly more rigid than every other state; State 7 forms an intermediate tier and likewise differs from every other state; and States 8, 1, 6, 5 and 9 form a relaxed, fluid tier that is mutually indistinguishable across all ten of its pairwise comparisons. Significance: **** *p*.adj < 10^-4^; *** < 10^-3^; ** < 10^-2^; * < 0.05; ns, not significant.

| State A | State B | MAC A | MAC B | $p$ (Wilcoxon) | $p$.adj (BH) | Signif. |
| --- | --- | --- | --- | --- | --- | --- |
| S4 | S9 | 0.361 | 0.147 | $2.7 \times 10^{-11}$ | $5.7 \times 10^{-10}$ | **** |
| S4 | S5 | 0.361 | 0.156 | $1.9 \times 10^{-10}$ | $2.0 \times 10^{-9}$ | **** |
| S4 | S6 | 0.361 | 0.167 | $2.0 \times 10^{-9}$ | $1.4 \times 10^{-8}$ | **** |
| S1 | S4 | 0.169 | 0.361 | $5.2 \times 10^{-9}$ | $2.7 \times 10^{-8}$ | **** |
| S4 | S8 | 0.361 | 0.183 | $4.9 \times 10^{-8}$ | $2.0 \times 10^{-7}$ | **** |
| S7 | S9 | 0.264 | 0.147 | $2.6 \times 10^{-5}$ | $8.9 \times 10^{-5}$ | **** |
| S5 | S7 | 0.156 | 0.264 | $1.0 \times 10^{-4}$ | $3.0 \times 10^{-4}$ | *** |
| S6 | S7 | 0.167 | 0.264 | $7.3 \times 10^{-4}$ | $1.9 \times 10^{-3}$ | ** |
| S1 | S7 | 0.169 | 0.264 | $1.9 \times 10^{-3}$ | $4.5 \times 10^{-3}$ | ** |
| S4 | S7 | 0.361 | 0.264 | $3.1 \times 10^{-3}$ | $6.5 \times 10^{-3}$ | ** |
| S7 | S8 | 0.264 | 0.183 | $4.7 \times 10^{-3}$ | $9.0 \times 10^{-3}$ | ** |
| S8 | S9 | 0.183 | 0.147 | 0.14 | 0.24 | ns |
| S1 | S9 | 0.169 | 0.147 | 0.16 | 0.26 | ns |
| S5 | S8 | 0.156 | 0.183 | 0.19 | 0.29 | ns |
| S1 | S5 | 0.169 | 0.156 | 0.25 | 0.35 | ns |
| S6 | S9 | 0.167 | 0.147 | 0.36 | 0.47 | ns |
| S5 | S6 | 0.156 | 0.167 | 0.48 | 0.59 | ns |
| S1 | S6 | 0.169 | 0.167 | 0.65 | 0.72 | ns |
| S6 | S8 | 0.167 | 0.183 | 0.62 | 0.72 | ns |
| S1 | S8 | 0.169 | 0.183 | 0.88 | 0.93 | ns |
| S5 | S9 | 0.156 | 0.147 | 0.95 | 0.95 | ns |

**S3 Table.** CSK global network rigidity: all pairwise contrasts among the four biological conditions. Global MAC of the CSK receiver is 0.134 (apo/apo), 0.134 (CSK-ATP), 0.148 (both-ATP) and 0.240 (primed, pY419). Each contrast is a two-sided Wilcoxon rank-sum test on the two conditions’ distributions of |Spearman ρ| across the 210 network edges (21 inter-landmark distances), Benjamini–Hochberg-adjusted over the 21 pairwise condition comparisons of the full series. The three unprimed conditions are mutually indistinguishable, so CSK rigidity is statistically flat until pY419 is installed; the main-text comparison (both-ATP versus primed) therefore isolates the mark as the single changed variable, and the stiffening cannot be attributed to nucleotide loading in either subunit. Δ MAC is the second condition minus the first. *n* = 100 models per condition; n.s., not significant; ***, *p*.adj < 10^-3^. Values are pooled per condition over that condition’s own metric panel (21 metrics and 210 edges for the ATP-loaded CSK conditions, 16 and 120 for apo), and so are sensitive to the condition’s metastable-state composition (Note 10).

| Contrast | $\Delta$ MAC | $p.\text{adj}$ (BH) | Signif. |
| --- | --- | --- | --- |
| apo/apo → CSK-ATP | −0.000 | 0.319 | n.s. |
| apo/apo → both-ATP | +0.014 | 0.522 | n.s. |
| CSK-ATP → both-ATP | +0.014 | 0.070 | n.s. |
| apo/apo → primed (pY419) | +0.106 | $5.3 \times 10^{-8}$ | *** |
| CSK-ATP → primed (pY419) | +0.106 | $2.7 \times 10^{-11}$ | *** |
| both-ATP → primed (pY419) | +0.092 | $3.3 \times 10^{-8}$ | *** |

**S4 Table.**
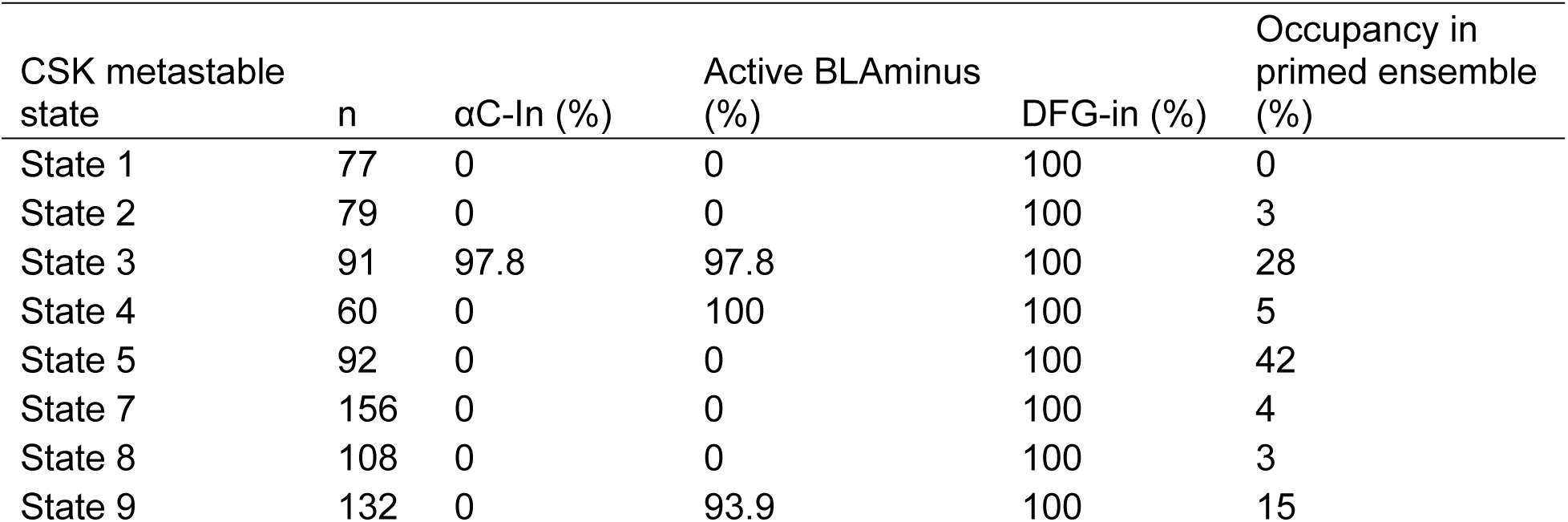
CSK conformational signatures are partitioned by metastable state, not graded across the ensemble. Categorical calls for all 795 CSK chains, tabulated by unsupervised GMM metastable state (states are defined across all seven conditions; Methods). The αC-In signature is confined to State 3; States 1, 2, 5, 7 and 8 contain no αC-In model at all, and the Active BLAminus population outside State 3 comes entirely from States 9 and 4, which adopt the active activation-loop backbone with the αC-helix still extruded. DFG-in is saturated in every state and therefore does not discriminate. The final column gives each state’s occupancy in the SRC pY419-primed condition (csk-holo/src-py159-holo, n = 100), from which the ensemble-level values quoted in Results follow to within a percentage point: 28% × 97.8% = 27% αC-In, and States 3, 9 and 4 together give 47% Active BLAminus. Percentages are of the models within that state. Source: plots_and_stats_CSK_GMM/Phase7_Complete_Structural_Metadata.csv.

| CSK metastable state | n | αC-In (%) | Active BLAminus (%) | DFG-in (%) | Occupancy in primed ensemble (%) |
| --- | --- | --- | --- | --- | --- |
| State 1 | 77 | 0 | 0 | 100 | 0 |
| State 2 | 79 | 0 | 0 | 100 | 3 |
| State 3 | 91 | 97.8 | 97.8 | 100 | 28 |
| State 4 | 60 | 0 | 100 | 100 | 5 |
| State 5 | 92 | 0 | 0 | 100 | 42 |
| State 7 | 156 | 0 | 0 | 100 | 4 |
| State 8 | 108 | 0 | 0 | 100 | 3 |
| State 9 | 132 | 0 | 93.9 | 100 | 15 |

**S5 Table.** Global network rigidity (MAC) before and after partialling out AlphaFold3 prediction confidence. For each experimental condition the correlation network was recomputed with per-model confidence removed from every edge (Methods), giving a partial-correlation MAC directly comparable to the published raw value. *n*, models per condition; Edges, unique metric pairs in that condition’s network; Δ, MAC (partial) − MAC (raw). The covariates partialled out are ipTM and mean interface PAE for the CSK–SRC series, and pTM for the CDK1 series, whose apo conditions are monomeric and have no interface. CSK–SRC conditions are given as CSK state / SRC state, where pY denotes the SRC activation-loop phosphomark pY419; CDK1 conditions are labelled by cofactor and phosphomark, with “(mono)” marking the cyclin-free monomers and triple-P the inhibitory pT14/pY15/pT161 complex. Partialling removes variance and therefore tends to lower MAC; the small positive Δ reflect noise in the edge-wise estimate. What is under test is the between-condition contrast (S6 Table), not the absolute value. Rows are ordered by raw MAC within each series. See Note 13.

| Series | Condition | $n$ | Edges | MAC (raw) | MAC (partial) | $\Delta$ |
| --- | --- | --- | --- | --- | --- | --- |
| CSK | apo/pY+ATP | 33 | 120 | 0.263 | 0.258 | -0.005 |
| CSK | ATP/pY+ATP | 100 | 210 | 0.240 | 0.246 | +0.006 |
| CSK | apo/pY | 100 | 120 | 0.214 | 0.212 | -0.002 |
| CSK | ATP/pY | 166 | 210 | 0.214 | 0.211 | -0.002 |
| CSK | ATP/ATP | 100 | 210 | 0.148 | 0.147 | -0.001 |
| CSK | apo/apo | 100 | 120 | 0.134 | 0.137 | +0.003 |
| CSK | ATP/apo | 196 | 210 | 0.134 | 0.133 | -0.002 |
| SRC | apo/pY+ATP | 33 | 210 | 0.180 | 0.184 | +0.004 |
| SRC | ATP/ATP | 100 | 210 | 0.129 | 0.125 | -0.005 |
| SRC | apo/pY | 100 | 120 | 0.129 | 0.128 | -0.001 |
| SRC | ATP/pY+ATP | 100 | 210 | 0.124 | 0.123 | -0.001 |
| SRC | apo/apo | 100 | 120 | 0.124 | 0.127 | +0.003 |
| SRC | ATP/apo | 196 | 120 | 0.120 | 0.121 | +0.000 |
| SRC | ATP/pY | 166 | 120 | 0.119 | 0.108 | -0.011 |
| CDK1 | apo (mono) | 100 | 91 | 0.365 | 0.231 | -0.134 |
| CDK1 | +CCNB1 | 100 | 120 | 0.217 | 0.171 | -0.045 |
| CDK1 | pT161 | 100 | 210 | 0.177 | 0.175 | -0.002 |
| CDK1 | triple-P | 100 | 210 | 0.166 | 0.161 | -0.005 |
| CDK1 | ATP (mono) | 100 | 171 | 0.136 | 0.138 | +0.002 |
| CDK1 | +CCNB1+ATP | 100 | 210 | 0.123 | 0.122 | -0.001 |

**S6 Table.**
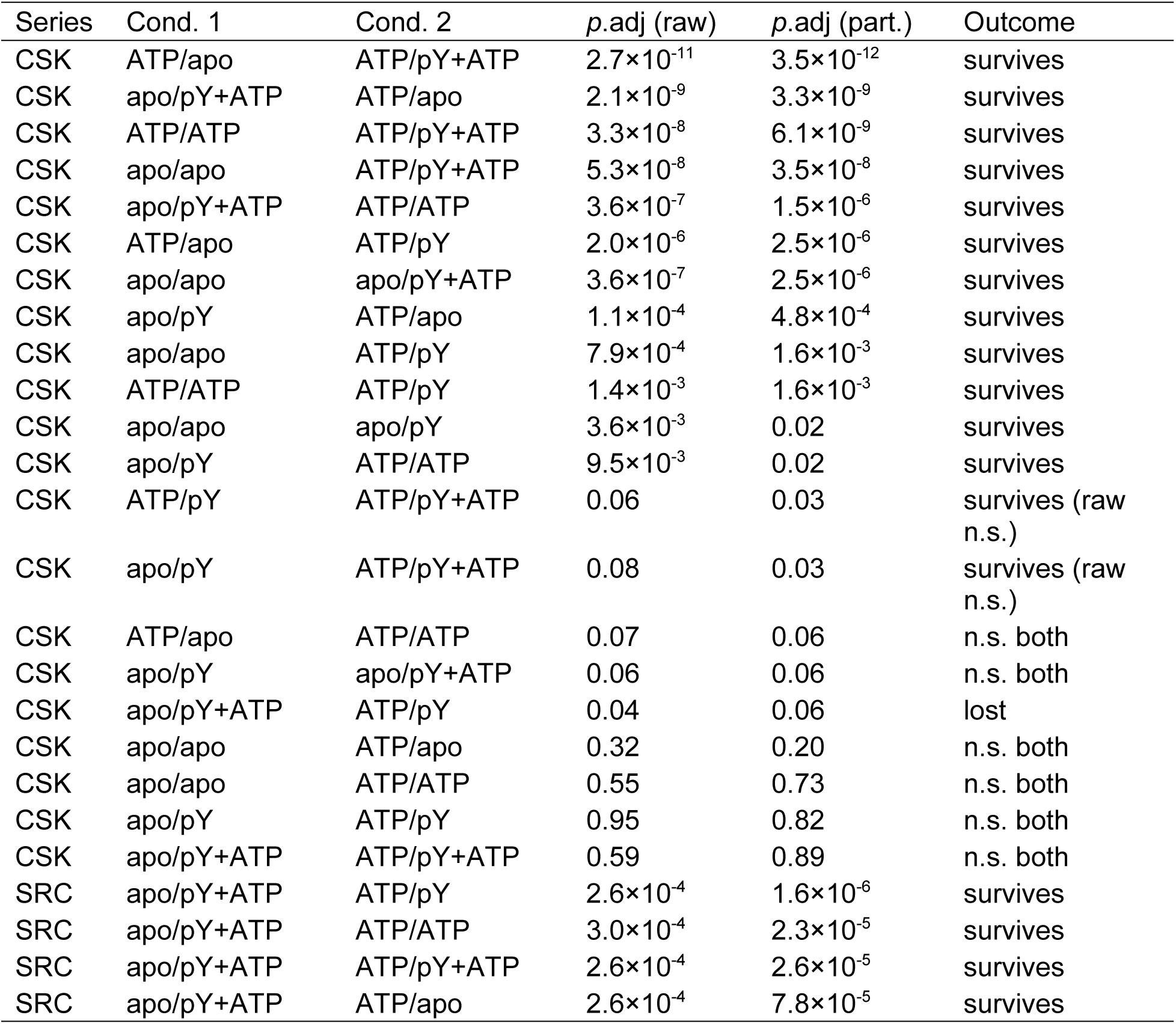
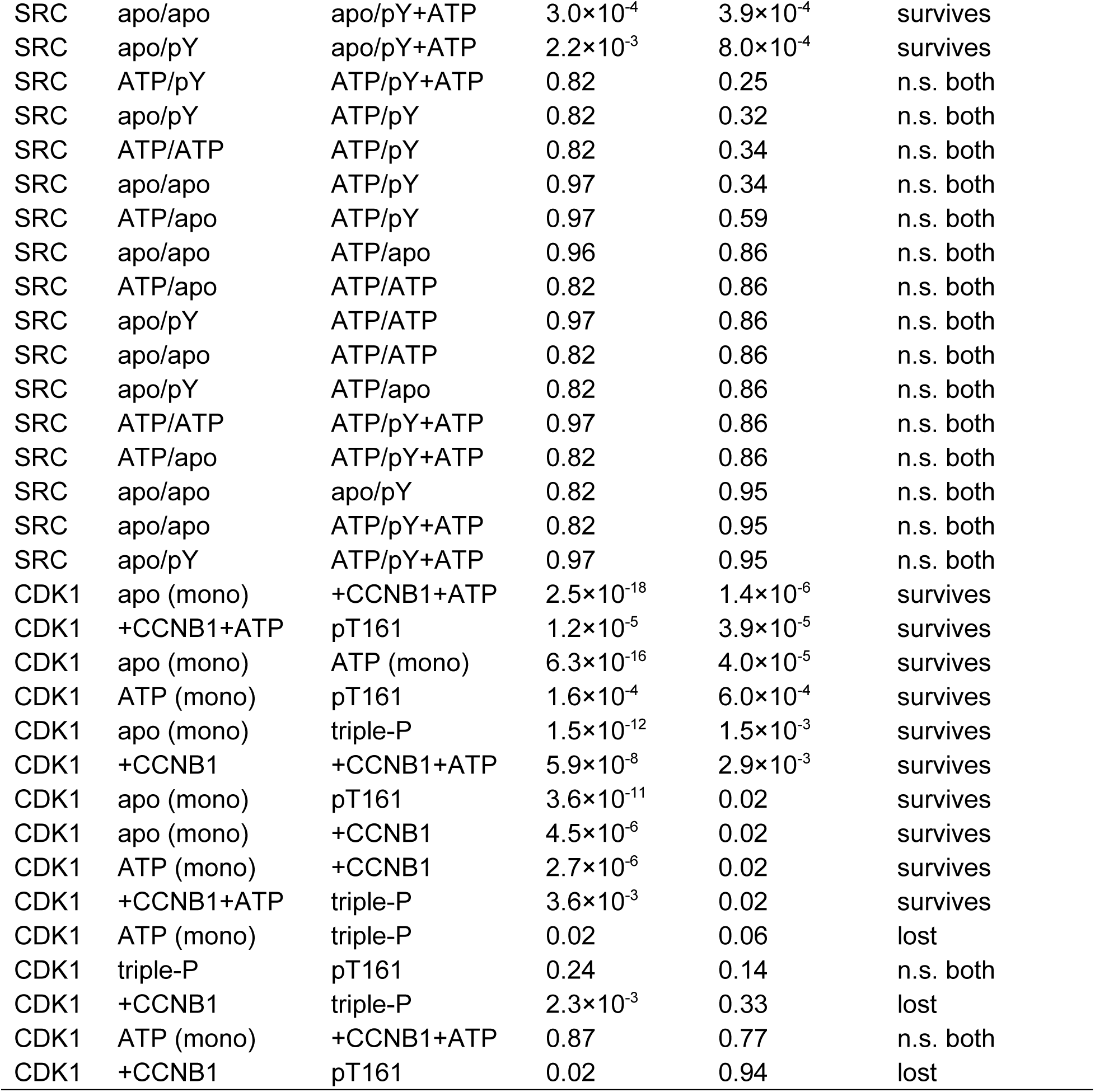
Between-condition MAC contrasts before and after partialling out prediction confidence. Every pairwise condition contrast in each series, tested by Mann–Whitney on the per-edge absolute correlations with Benjamini–Hochberg adjustment within series, computed on the raw edges (*p*.adj raw) and on the confidence-partialled edges (*p*.adj part.). Condition labels, and the raw and partial MAC value of every condition, are given in S5 Table. Outcome: “survives”, significant (*p*.adj < 0.05) both before and after partialling with the direction of the difference preserved; “survives (raw n.s.)”, significant only after partialling; “lost”, significant before but not after; “n.s. both”, not significant either way. Rows are ordered by *p*.adj (partial) within each series. See Note 13.

**S7 Table.** Pooled and per-state network rigidity for the four biological CSK conditions. Pooled MAC is the single condition-level network for each condition. Per-state values recompute the same statistic within each macro-state the condition occupies with at least 8 models, on that condition’s own metric panel. They are tabulated rather than plotted alongside, because MAC rises as sample size falls and the two are not comparable by magnitude. Metrics / edges is the size of that condition’s network. The panel differs between apo and ATP-loaded conditions because ligand-anchored metrics are undefined without bound nucleotide, so per-state values are comparable with their own condition’s pooled value but not between conditions. The occupancy-weighted row averages a condition’s per-state values by occupancy over the states listed and therefore carries a smaller *n* than the condition, the difference being the states below the 8-model threshold. Occupancy is the state’s share of the condition after Truth Overwrite reassignment, which is why CSK-ATP carries *n* = 196. The null floor is the mean |ρ| expected between independent variables at that sample size, the square root of 2/[π(*n* – 1)]; a value approaching its floor carries no information, which is why the sparsest states of apo/apo and CSK-ATP show the table’s largest MAC. The size-matched null is the mean MAC of 2000 random subsets of the same condition at that row’s own sample size, state labels ignored, with *z* the departure from it in null standard deviations. It is the reference for whether a state is less coupled than the ensemble containing it, whereas the null floor beside it addresses only whether the cell carries information; pooled rows, being the whole condition, have none. See Note 10.

| Condition | Metrics / edges | Macro-state | $n$ | Occupancy | MAC | Null floor | Size-matched null | $z$ |
| --- | --- | --- | --- | --- | --- | --- | --- | --- |
| apo/apo | 16 / 120 | pooled | 100 | 100% | 0.134 | 0.080 | — | — |
| apo/apo | 16 / 120 | State 1 | 73 | 73% | 0.138 | 0.094 | 0.143 | -0.72 |
| apo/apo | 16 / 120 | State 2 | 13 | 13% | 0.288 | 0.230 | 0.259 | +1.39 |
| apo/apo | 16 / 120 | State 9 | 12 | 12% | 0.270 | 0.241 | 0.270 | +0.02 |
| apo/apo | 16 / 120 | occupancy-weighted | 98 | 98% | 0.174 | 0.130 | 0.174 | +0.03 |
| CSK-ATP | 21 / 210 | pooled | 196 | 100% | 0.134 | 0.057 | — | — |

| Condition | Metrics /<br>edges | Macro-state | $n$ | Occupancy | MAC | Null<br>floor | Size-matched<br>null | $z$ |
| --- | --- | --- | --- | --- | --- | --- | --- | --- |
| CSK-ATP | 21 / 210 | State 7 | 96 | 49% | 0.147 | 0.082 | 0.150 | -0.45 |
| CSK-ATP | 21 / 210 | State 8 | 62 | 32% | 0.142 | 0.102 | 0.165 | -2.10 |
| CSK-ATP | 21 / 210 | State 9 | 22 | 11% | 0.172 | 0.174 | 0.221 | -2.91 |
| CSK-ATP | 21 / 210 | State 3 | 10 | 5% | 0.281 | 0.266 | 0.304 | -1.00 |
| CSK-ATP | 21 / 210 | occupancy<br>-weighted | 190 | 97% | 0.155 | 0.109 | 0.171 | -4.72 |
| both-ATP | 21 / 210 | pooled | 100 | 100% | 0.148 | 0.080 | — | — |
| both-ATP | 21 / 210 | State 7 | 45 | 45% | 0.198 | 0.120 | 0.176 | +2.47 |
| both-ATP | 21 / 210 | State 8 | 40 | 40% | 0.161 | 0.128 | 0.182 | -2.05 |
| both-ATP | 21 / 210 | occupancy<br>-weighted | 85 | 85% | 0.181 | 0.124 | 0.179 | +0.45 |
| primed | 21 / 210 | pooled | 100 | 100% | 0.240 | 0.080 | — | — |
| primed | 21 / 210 | State 5 | 42 | 42% | 0.166 | 0.125 | 0.257 | -5.21 |
| primed | 21 / 210 | State 3 | 28 | 28% | 0.168 | 0.154 | 0.271 | -4.84 |
| primed | 21 / 210 | State 9 | 15 | 15% | 0.236 | 0.213 | 0.307 | -2.43 |
| primed | 21 / 210 | occupancy<br>-weighted | 85 | 85% | 0.179 | 0.150 | 0.270 | -11.19 |

**S8 Table.** Pairwise comparisons of per-state intrinsic network rigidity (CSK). The CSK counterpart of S2 Table. Each metastable state’s intrinsic MAC is the mean absolute Spearman correlation across its structural-metric network, computed on the universal 16-metric, 120-edge panel so that every state shares one basis; it is therefore not comparable with the pooled per-condition values of S7 Table, which use each condition’s own panel. Intrinsic MAC pools all models of a state regardless of condition, including the overwrite-artifact designs excluded from Fig. 5, as in Note 14; each state’s cell size is given beside its label (*n* = 60 to 156 models). State pairs were compared by Wilcoxon rank-sum test on the per-edge absolute correlations, with Benjamini–Hochberg adjustment across all 28 comparisons; rows are ordered by *p*.adj. Only State 2 separates from any other state. The three states that co-occur with primed SRC (States 5, 3 and 9; Fig. 5B) are indistinguishable from each other and from States 7 and 8, which dominate the unprimed complex, in all nine of those comparisons: CSK gains no internal coupling on priming (Notes 9 and 10). Margins over the small-sample floor range from 1.4-fold (State 4, 0.143 against 0.104) to 2.2-fold (State 9, 0.154 against 0.070), so none of these comparisons is floor-limited. Significance: **** *p*.adj < 10^-4^; *** < 10^-3^; ** < 10^-2^; * < 0.05; ns, not significant.

| State A | State B | MAC A | MAC B | $p$ (Wilcoxon) | $p$.adj (BH) | Signif. |
| --- | --- | --- | --- | --- | --- | --- |
| S2 (79) | S5 (92) | 0.172 | 0.121 | $7.3 \times 10^{-4}$ | $9.9 \times 10^{-3}$ | ** |
| S2 (79) | S7 (156) | 0.172 | 0.121 | $1.1 \times 10^{-3}$ | $9.9 \times 10^{-3}$ | ** |
| S2 (79) | S8 (108) | 0.172 | 0.121 | $1.1 \times 10^{-3}$ | $9.9 \times 10^{-3}$ | ** |
| S2 (79) | S3 (91) | 0.172 | 0.127 | $3.1 \times 10^{-3}$ | 0.02 | * |
| S1 (77) | S2 (79) | 0.137 | 0.172 | 0.02 | 0.10 | ns |
| S2 (79) | S4 (60) | 0.172 | 0.143 | 0.04 | 0.13 | ns |
| S5 (92) | S9 (132) | 0.121 | 0.154 | 0.03 | 0.13 | ns |
| S7 (156) | S9 (132) | 0.121 | 0.154 | 0.04 | 0.13 | ns |
| S8 (108) | S9 (132) | 0.121 | 0.154 | 0.03 | 0.13 | ns |
| S2 (79) | S9 (132) | 0.172 | 0.154 | 0.15 | 0.34 | ns |
| S3 (91) | S9 (132) | 0.127 | 0.154 | 0.16 | 0.34 | ns |
| S4 (60) | S5 (92) | 0.143 | 0.121 | 0.15 | 0.34 | ns |
| S4 (60) | S8 (108) | 0.143 | 0.121 | 0.16 | 0.34 | ns |
| S4 (60) | S7 (156) | 0.143 | 0.121 | 0.19 | 0.38 | ns |
| S1 (77) | S5 (92) | 0.137 | 0.121 | 0.30 | 0.46 | ns |
| S1 (77) | S7 (156) | 0.137 | 0.121 | 0.29 | 0.46 | ns |
| S1 (77) | S8 (108) | 0.137 | 0.121 | 0.30 | 0.46 | ns |
| S1 (77) | S9 (132) | 0.137 | 0.154 | 0.26 | 0.46 | ns |
| S3 (91) | S4 (60) | 0.127 | 0.143 | 0.46 | 0.58 | ns |
| S3 (91) | S5 (92) | 0.127 | 0.121 | 0.46 | 0.58 | ns |
| S3 (91) | S7 (156) | 0.127 | 0.121 | 0.48 | 0.58 | ns |
| S3 (91) | S8 (108) | 0.127 | 0.121 | 0.45 | 0.58 | ns |
| S4 (60) | S9 (132) | 0.143 | 0.154 | 0.48 | 0.58 | ns |
| S1 (77) | S3 (91) | 0.137 | 0.127 | 0.70 | 0.78 | ns |
| S1 (77) | S4 (60) | 0.137 | 0.143 | 0.70 | 0.78 | ns |
| S5 (92) | S7 (156) | 0.121 | 0.121 | 0.91 | 0.95 | ns |
| S7 (156) | S8 (108) | 0.121 | 0.121 | 0.91 | 0.95 | ns |
| S5 (92) | S8 (108) | 0.121 | 0.121 | 0.99 | 0.99 | ns |

**S9 Table.**
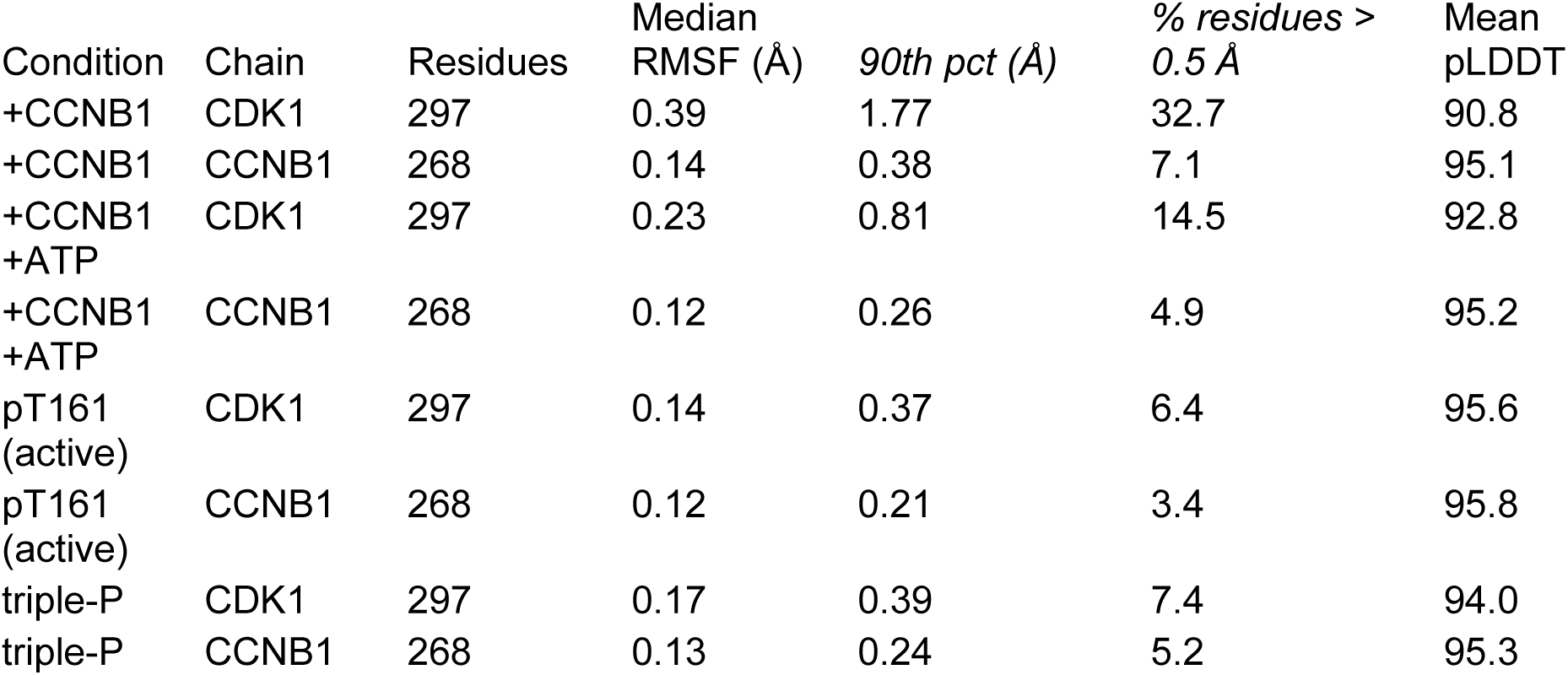
Chain-level conformational stability in the CDK1–CCNB1 complexes. Per-residue Cα RMSF after chain-local superposition, across the 100 models of each condition (Methods). Lower values indicate a more reproducibly placed fold. CCNB1 is the less variable chain in every condition and is insensitive to nucleotide and phosphorylation state. MAC is not reported for CCNB1 because it is undefined for a non-kinase chain. Monomer context, not part of the comparison: apo CDK1 alone gives median 0.46 Å (mean pLDDT 89.0) and CDK1+ATP alone 0.17 Å (93.6).

### Supplementary Data

**S1 Dataset.** AlloQuant metric dictionary. AlloQuant_master_CSV_data_dictionary_v7r3.pdf defines all per-model output columns, their biological meaning, measurement approach, and units. Column names in this document and in figure panel labels refer to entries in this dictionary.

**S2 Dataset.** AlloQuant output file manifest. AlloQuant_output_file_manifest_v7r3.pdf lists every file generated by AlloQuant, its format, and the information it contains.

**S3 Dataset.** AlloQuant outputs underlying the reported values. S3_Dataset.zip holds the Module 1 per-model measurements and the Module 2 statistical outputs from which every number reported in this work is computed, together with a README describing the layout and a manifest of md5 checksums for all 195 files. The same files are archived with the analysis code (doi:10.5281/zenodo.22208357).

