## Supplementary material for "Trans-Allosteric Activation Releases Distinct Conformational Traps in Kinase Heterodimers": S1 Data

### AlloQuant master results - data dictionary

*File: master_kinase_analysis_results_v7r3.csv | 1600 rows x 48 columns | generated 2026-07-30*

- This dictionary defines the AlloQuant v7r3 master-CSV SCHEMA. The 48 columns are identical across all v7r3 masters (SRC 260606, SRC 260718 rerun, CDK1-CCNB1); only which rows are populated varies by run.
- Anchored to the SRC 260718 rerun (the AF3-complete master). Empirical counts/ranges below are from that file: 8 systems (CSK apo/holo x SRC apo/holo x SRC WT/pY159) x 25 AF3 models x 2 chains = 1600 rows.
- AF3 confidence columns (ipTM/pTM/PAE_*) are populated in this rerun (it was re-predicted to capture summary_confidences.json/confidences.json). They are N/A in the original SRC 260606 master.
- The CDK1-CCNB1 master is single-chain (kinase only, 600 rows, same 48 columns). Its AF3 columns are not uniformly N/A: pTM 600/600; ipTM 500/600 (N/A only for the ligand-free apo monomer); interface-PAE 400/600 (the CCNB1-containing rows only).
- holo = ATP(/Mg)-bound; apo = no nucleotide. C_Spine == 'No Ligand' flags an apo chain.
- Distances in Angstrom (A), angles in degrees. Landmark indices are 0-based HMM-aligned positions; numbered landmarks (n99, v104, k105, e107, m118, m120, e121, i150, y156, d220) are labelled in CSK numbering and map to the equivalent target position via the HMM.
- Missing / uninformative values: geometry that fails to compute is written as the sentinel 999.9; everything else uninformative is the string 'N/A' (or 'None' for CoFactor_Name / Partner). Downstream R reads '', 'NA', 'N/A', 'None' all as NA.
- Writers: columns 1-43 by modules/chimerax_hmm_worker_v7r3.py; columns 44-48 (ipTM, pTM, PAE_*) by modules/extract_af3_metrics.py, joined on Directory.
- v7r2 -> v7r3 (2026-07-29) CHANGES REPORTED VALUES: re-run any dataset before comparing across versions. Three Module-1 corrections, below. Dihedral, D2_Dist, SB_Dist and every other column are unchanged.
- (i) D1_Dist is now measured from the alphaC-Glu(+4) Calpha per Modi & Dunbrack, PNAS 2019 (116:6818), not the alphaC-Glu itself; the published 11/14 A cutoffs are calibrated on the (+4) anchor. This also changes Spatial and State.
- (ii) the ActLoop_CT anchor is the HRD arginine (hrd+1), not hrd+6, which is not a conserved position (an Arg in CSK but an Ala in SRC).
- (iii) a missing-density guard makes ActLoop_NT/ActLoop_CT return N/A across unresolved residues instead of measuring over the gap; no effect on complete AlphaFold3 chains.

| # | Column | Group | Type | Units | Populated | Allowed values / range | Empirical (this CSV) | Description |
|---|---|---|---|---|---|---|---|---|
| 1 | **Simulation_ID** | Identity | string | | always | | filled 1600/1600 | Unique model id, from the model path via get_core_name() (dedups repeated path tokens). Encodes the two chains, AF3 seed and sample, e.g. a-csk-wtcat-apo_b-src-wtcat-holo_seed-..._sample-N_model. |
| 2 | **Directory** | Identity | string (path) | | always | | filled 1600/1600 | Relative path to the AF3 prediction folder (system / seed-sample). Also the join key used to merge in the AF3 confidence columns. |
| 3 | **File** | Identity | string | | always | model.cif | filled 1600/1600 | Structure file analysed within Directory (AF3 predicted mmCIF). |
| 4 | **Chain** | Identity | categorical | | always | A, B, D | filled 1600/1600; observed: A 800, B 400, D 400 | mmCIF chain id of the protein chain in this row. A = CSK protomer; B or D = SRC protomer (the letter shifts to D when holo ligand chains are inserted ahead of it). One row per protein chain; 2 rows per model. |
| 5 | **Type** | Identity | categorical | | always | CSK-WT, SRC-WT | filled 1600/1600; observed: CSK-WT 800, SRC-WT 800 | Kinase identity assigned by best sequence-identity match to the FASTA landmark set (get_best_landmark_for_chain). Note the SRC pY159 variant still matches as SRC-WT; the pY159 vs WT distinction lives in Simulation_ID/Directory, not here. |
| 6 | **State** | Conformation | categorical | | always | see Description | filled 1600/1600; observed: Active (BLAminus) 1063, Inactive (BLBplus) 524, Inactive (BLBminus) 5, Inactive (BLAplus) 5, Inactive (BLBtrans) 1, Active-Like (ABAminus) 1, DFGinter (BABplus) 1 | Human-readable state = Spatial + Dihedral with an activity gloss: Active (BLAminus), Active-Like (ABAminus), Inactive (BLBplus / BLBtrans / BLAplus / BLBminus / BBAminus / BABtrans), else a fall-through <spatial> (<dihedral>) such as DFGinter (BABplus). Downstream analysis treats 'Active (BLAminus)' as the active target. |
| 7 | **Role** | Interface | categorical | | always (dimer) | C-lobe Donor, N-lobe Receiver, Symmetric, Unpaired, Co-factor Bound | filled 1600/1600; observed: N-lobe Receiver 799, C-lobe Donor 799, Unpaired 2 | Role in the asymmetric dimer from analyze_dimer_interface: a chain is C-lobe Donor / N-lobe Receiver when its C-lobe contacts the partner N-lobe within 8 A (Symmetric if mutual, Unpaired if no contact). For CSK-SRC: CSK=N-lobe Receiver, SRC=C-lobe Donor. |
| 8 | **Partner** | Interface | categorical | | always (dimer) | A, B, D | filled 1598/1600, blank 2; observed: A 799, B 400, D 399 | Chain id of this chain's interface partner (or None if Unpaired; a cofactor name for Co-factor Bound systems). |
| 9 | **CoFactor_Name** | Cofactor | string | | N/A (no cofactor) | | filled 0/1600, blank 1600 | Name of the nearest non-kinase chain within 15 A, else None. Populated only for cofactor systems (CDK-cyclin, RAF/14-3-3, etc.); empty for the CSK-SRC pair. |
| 10 | **CoFactor_aC_Dist** | Cofactor | float | A | N/A (no cofactor) | | filled 0/1600, blank 1600 | Min distance from the kinase alphaC region (c-4..c+4) to the cofactor chain. Cofactor systems only. |
| 11 | **CoFactor_ActLoop_Dist** | Cofactor | float | A | N/A (no cofactor) | | filled 0/1600, blank 1600 | Min distance from the kinase activation loop (f..APE) to the cofactor chain. Cofactor systems only. |
| 12 | **Interface_C_Lobe_Donor_Dist** | Interface | float | A | always (dimer) | 2.44 to 13.24 | filled 1598/1600, blank 2 | Min distance between chain-A C-lobe and chain-B N-lobe atoms (donor face). 999.9 sentinel if lobes cannot be split; N/A if unpaired. |
| 13 | **Interface_N_Lobe_Rec_Dist** | Interface | float | A | always (dimer) | 2.44 to 13.24 | filled 1598/1600, blank 2 | Min distance between chain-B C-lobe and chain-A N-lobe atoms (receiver face). Same sentinel/NA conventions. |
| 14 | **R_Spine** | Spine/helix | categorical | | always | Intact, Broken, Missing | filled 1600/1600; observed: Intact 1447, Broken 153 | Regulatory-spine integrity: Intact if all three R-spine contacts (HRD-His-DFG-Phe, DFG-Phe-rs1, rs1-rs2) are <4.5 A, else Broken (Missing if landmarks absent). |
| 15 | **C_Spine** | Spine/helix | categorical | | always | Intact, Ligand Distant, No Ligand | filled 1600/1600; observed: No Ligand 800, Intact 800 | Catalytic-spine vs adenine: Intact if the VAIK region (k..k-3) is <6 A from a bound ligand, Ligand Distant if >=6, No Ligand if apo. Serves as the apo/holo discriminator downstream (No Ligand = apo chain). |
| 16 | **C_Helix** | Spine/helix | categorical | | always | In, Out | filled 1600/1600; observed: In 878, Out 722 | alphaC rotamer: In if the VAIK-Lys CB to alphaC-Glu CB distance is <=10 A (salt-bridge competent), else Out. In = active-like. |
| 17 | **Shell_State** | Spine/helix | categorical | | always | Packed, Loose | filled 1600/1600; observed: Packed 1600 | Regulatory-shell packing: Packed if Shell_M118_M120_Dist <5 A, else Loose (only Packed observed in this run). |

- v7r2 -> v7r3 (2026-07-29) CHANGES REPORTED VALUES: re-run any dataset before comparing across versions. Three Module-1 corrections, below. Dihedral, D2_Dist, SB_Dist and every other column are unchanged.

| | | | | | | | | |
|---|---|---|---|---|---|---|---|---|
| 18 | **Spatial** | Conformation | categorical | | always | DFGin, DFGout, DFGinter, Outlier | filled 1600/1600; observed: DFGin 1599, DFGinter 1 | Dunbrack spatial DFG group from D1_Dist/D2_Dist: DFGin (D1<=11 & D2>=11), DFGout (D1>11 & D2<=14), DFGinter (both <=11), else Outlier. All four are reachable; correcting the D1 anchor in v7r3 removed the inflated Outlier/DFGout calls seen in v7r2. |
| 19 | **Dihedral** | Conformation | categorical | | always | BLAminus, BLBplus, ... | filled 1600/1600; observed: BLAminus 1063, BLBplus 524, BLBminus 5, BLAplus 5, BLBtrans 1, ABAminus 1, BABplus 1 | KinCore backbone cluster = Ramachandran regions of X(f-2), Asp(f-1), Phe(f) + DFG-Phe chi1 rotamer, e.g. BLAminus/ABAminus/BLBplus/BLBtrans/BLAplus/BLBminus. Region codes A/B/L/E; rotamer plus/trans/minus. BLAminus = canonical active. |
| 20 | **ActLoop_NT** | Conformation | categorical | | always | NTin, NTout, N/A | filled 1600/1600; observed: NTin 1170, NTout 430 | Activation-loop N-terminal packing: NTin if residues f+3..f+6 come within 5.5 A of the residue before HRD (hrd-1), else NTout; N/A if those residues are not contiguous in the deposited numbering (missing-density guard). |
| 21 | **ActLoop_CT** | Conformation | categorical | | always | CTin, CTout, N/A | filled 1600/1600; observed: CTin 1564, CTout 36 | Activation-loop C-terminal packing: CTin if the pre-APE window (ape-6 down to ape-9, floored at f+2) comes within 5.5 A (all-atom) of the HRD arginine (hrd+1), else CTout; N/A if the residues are not contiguous in the deposited numbering. The hrd+1 anchor follows Modi & Dunbrack's APE9-Arg contact (v7r2 used hrd+6, which is an Arg in CSK but an Ala in SRC). The 5.5 A cutoff is AlloQuant's own sensitivity choice, deliberately looser than Kincore's 6.0 A. |
| 22 | **Phi_D** | Conformation | float | degrees | always | -100.82 to 70.18 | filled 1600/1600 | Backbone phi dihedral of the DFG-Asp (f-1). |
| 23 | **Psi_D** | Conformation | float | degrees | always | 15.46 to 113.62 | filled 1600/1600 | Backbone psi dihedral of the DFG-Asp (f-1). |
| 24 | **Cleft_Gape_Dist** | Active site | float | A | always | 11.65 to 19.61 | filled 1600/1600 | Inter-lobe cleft opening: roof Calpha to floor Calpha, where roof = a P-loop residue (phospho/Tyr/Thr/Ser or P-loop midpoint) and floor = catalytic HRD-Asp (hrd+2). Geometric, so populated for apo and holo. |
| 25 | **Mg_Hijack_Dist** | Active site | float | A | holo + Mg only | 7.64 to 64.16 | filled 1200/1600, blank 400 | Min distance from the roof residue phosphate/hydroxyl atoms to any Mg2+ ion. Requires a bound Mg2+; N/A on apo chains. |
| 26 | **Substrate_Clearance_Angle** | Active site | float | degrees | nucleotide-bound only | 88.33 to 158.70 | filled 800/1600, blank 800 | Angle roof-Calpha / ATP-phosphate / floor-Calpha (phosphate atom as vertex). Requires a bound nucleotide; N/A on apo chains. |
| 27 | **D1_Dist** | Active site | float | A | always | 4.69 to 9.89 | filled 1600/1600 | Dunbrack D1 coordinate: alphaC-Glu(+4) Calpha to DFG-Phe CZ (farthest side-chain atom if CZ absent). The (+4) anchor is the one the 11/14 A Spatial cutoffs are calibrated on (Modi & Dunbrack, PNAS 2019, 116:6818); v7r2 measured from the alphaC-Glu itself, which compressed the range. |
| 28 | **D2_Dist** | Active site | float | A | always | 7.80 to 16.74 | filled 1600/1600 | Dunbrack D2 coordinate: VAIK-Lys Calpha to DFG-Phe CZ. |
| 29 | **SB_Dist** | Active site | float | A | always | 2.44 to 18.43 | filled 1600/1600 | beta3 Lys - alphaC Glu salt bridge: min distance from VAIK-Lys NZ to the nearest polar (O*/N*) side-chain atom of the alphaC-Glu. Short (~2.5-3 A) = intact/active. |
| 30 | **HRD_ATP_Dist** | Active site | float | A | holo only | 2.47 to 8.92 | filled 800/1600, blank 800 | Min distance from catalytic HRD-Asp (hrd+2) side-chain O*/N* to the ATP gamma/beta/alpha-phosphate group. Requires a bound nucleotide; N/A on apo chains. |
| 31 | **DFG_Mg_Dist** | Active site | float | A | holo + Mg only | 0.91 to 6.97 | filled 796/1600, blank 804 | Min distance DFG-Asp (f-1) side-chain O*/N* to the nearest Mg2+ (recorded only if <=15 A). Requires Mg2+; N/A on apo chains. |
| 32 | **DFG_ATP_Dist** | Active site | float | A | holo only | 1.26 to 4.70 | filled 800/1600, blank 800 | Min distance DFG-Asp side-chain O*/N* to ATP phosphate atoms. Requires a bound nucleotide; N/A on apo chains. |
| 33 | **PLoop_ATP_Dist** | Active site | float | A | holo only | 1.44 to 5.22 | filled 800/1600, blank 800 | Min distance from P-loop backbone/CB atoms to ligand phosphate atoms. Requires a bound nucleotide; N/A on apo chains. |
| 34 | **aCb4_aE_Dist** | Allosteric | float | A | always | 2.81 to 3.72 | filled 1600/1600 | Min polar-atom (N*/O*) distance between the alphaC-beta4 loop (c+8..c+14) and the alphaE helix (hrd-25..hrd-10). Allosteric relay bridge. |
| 35 | **Spine_Bridge_Dist** | Allosteric | float | A | always | 3.57 to 8.36 | filled 1600/1600 | alphaC-beta4 loop to ligand if a ligand is bound; otherwise (apo) loop to k-3 residue. Populated for both apo and holo (fallback for apo). |
| 36 | **V104_RS2_Dist** | Allosteric | float | A | always | 6.81 to 7.38 | filled 1600/1600 | Min side-chain distance V104 to R-spine residue rs2 (beta4). |
| 37 | **I150_HRD_Dist** | Allosteric | float | A | always | 9.65 to 10.51 | filled 1600/1600 | Min side-chain distance I150 to the catalytic HRD-His. |
| 38 | **Shell_M118_M120_Dist** | Allosteric | float | A | always | 3.57 to 4.45 | filled 1600/1600 | Min side-chain distance between the two shell methionines M118 and M120 (drives Shell_State). |
| 39 | **Y156_N99_Dist** | Allosteric | float | A | always | 7.87 to 9.86 | filled 1600/1600 | Residue-min distance Y156 to N99 (alphaE anchor). |
| 40 | **K105_E107_Dist** | Allosteric | float | A | always | 2.95 to 5.71 | filled 1600/1600 | Min side-chain distance K105 to E107. |
| 41 | **K105_E121_Dist** | Allosteric | float | A | always | 2.97 to 4.16 | filled 1600/1600 | Min side-chain distance between residues at Pfam Pkinase HMM node 62 (PKA canonical K105 position; GLN65 in CSK domain numbering) and HMM node 78 (PKA canonical E121 position; GLU82 in CSK domain numbering). Part of the alphaC-beta4 loop contact network. NOT the beta3-Lys/alphaC-Glu regulatory salt bridge -- for that, see SB_Dist. |
| 42 | **K105_N99_Dist** | Allosteric | float | A | always | 2.72 to 7.11 | filled 1600/1600 | Min side-chain distance K105 to N99. |
| 43 | **D220_HRD_Dist** | Allosteric | float | A | always | 3.35 to 3.84 | filled 1600/1600 | Min side-chain distance D220 to the catalytic HRD-His. |
| 44 | **ipTM** | AF3 confidence | float | 0-1 | dimers | 0.32 to 0.90 | filled 1600/1600 | AF3 interface pTM from summary_confidences.json (>0.8 high, 0.6-0.8 moderate, <0.6 low). N/A for single-entity/monomer models. Populated in this rerun; N/A in the original 260606 master. |
| 45 | **pTM** | AF3 confidence | float | 0-1 | always | 0.57 to 0.91 | filled 1600/1600 | AF3 predicted TM-score. Populated in this rerun; N/A in the original 260606 master. |
| 46 | **PAE_A_to_B** | AF3 confidence | float | A | dimers | 8.40 to 27.45 | filled 1600/1600 | Mean predicted aligned error over the [chain-A rows, chain-B cols] PAE block (<5 rigid, >15 non-interacting). N/A for monomers. |
| 47 | **PAE_B_to_A** | AF3 confidence | float | A | dimers | 7.99 to 26.56 | filled 1600/1600 | Mean PAE over the transposed [chain-B rows, chain-A cols] block. N/A for monomers. |
| 48 | **PAE_Mean_AB** | AF3 confidence | float | A | dimers | 8.43 to 27.00 | filled 1600/1600 | Symmetric interface PAE = mean of PAE_A_to_B and PAE_B_to_A. N/A for monomers. |
