## Supplementary material for "Trans-Allosteric Activation Releases Distinct Conformational Traps in Kinase Heterodimers": S2 Data

### AlloQuant - output-file manifest

*Reference run: CSK-SRC dimer scan, 260718 rerun (AF3-complete) | generated 2026-08-31*

- AlloQuant is a two-module pipeline. Module 1 (structural extraction) writes the run-root files below, except row 1: experiment.csv is the user-supplied design file it reads. Module 2 (allostery discovery) writes one plots_and_stats_{KINASE}_{METHOD}/ folder per analysed kinase.
- Tokens: {KINASE} = analysed kinase (CSK, SRC, CDK1, ...); {METHOD} = clustering method (GMM default, or KMEANS); {ver} = pipeline version (v7r3); {i}/{j} = discovered-state indices.
- For this CSK-SRC rerun there are two Module-2 folders: plots_and_stats_CSK_GMM/ and plots_and_stats_SRC_GMM/ (identical file set, different target kinase).
- Interactive HTMLs need their sibling *_files/ asset folder to render offline; keep the pair together when sharing.
- MAC = Mean Absolute Correlation (global network rigidity) = mean of absolute metric-metric correlation-edge weights; used both per-condition (Phase 5) and per-state (Phase 6b).
- Discovered-state LABELS are not always contiguous: this run's CSK target populates 8 states labelled up to State 9 (State 6 is empty), so Phase 8 emits C(8,2) = 28 state pairs, not C(9,2).
- 'exists (this run)' is auto-checked against plots_and_stats_CSK_GMM/ (Module 2) and the run root (Module 1).

| # | Stage | Phase | File / pattern | Format | Description | exists (this run) |
|---|---|---|---|---|---|---|
| 1 | User input | Setup | `experiment.csv` | CSV | User-supplied experimental design metadata: one row per dimer condition (chain_a, condition_a, chain_b, ptm_b, condition_b). Read by the pipeline to define the systems built and analysed; it is not written by AlloQuant and is excluded from run archiving. | yes |
| 2 | Module 1 | Setup | `generated_matrix.csv` | CSV | Combinatorial multimer matrix (chain-A rows x chain-B columns) marking which pairwise AF3 jobs were generated. | yes |
| 3 | Module 1 | Setup | `fasta_source_map.json` | JSON | Maps each construct label to its source FASTA sequence and provenance (used to identify/type each chain). | yes |
| 4 | Module 1 | Landmarks | `hmm_landmarks.json` | JSON | Per-kinase-type HMM landmark residue indices (VAIK-Lys k, alphaC-Glu c, R-spine rs1/rs2, DFG f, HRD, and numbered allosteric landmarks). Consumed by every distance metric. | yes |
| 5 | Module 1 | Master | `master_kinase_analysis_results_{ver}.csv` | CSV | PRIMARY OUTPUT: one row per protein chain per model, with all conformational labels and allosteric distances. Documented column-by-column in the companion data dictionary. | yes |
| 6 | Module 2 | Phase 0 | `Phase0_AF3_Confidence_Metrics.pdf` | PDF | AF3 confidence & QC plots (ipTM / pTM / PAE distributions) across conditions. | yes |
| 7 | Module 2 | Phase 0 | `Phase0_AF3_Quality_Summary.csv` | CSV | Per-condition AF3 confidence/quality summary table underlying the Phase 0 plots. | yes |
| 8 | Module 2 | Phase 1 | `Phase1_AF3_Ligand_Placement_Preference.pdf` | PDF | System-level ligand occupancy ('tug-of-war'): where AF3 places ATP/ligand between the two chains. Ligand-bearing systems only (absent for ligand-free targets). | yes |
| 9 | Module 2 | Phase 1 | `Phase1_Ligand_QC_Review.csv` | CSV | Per-model ligand-placement QC flags and mismatch-override decisions (apo/holo assignment). | yes |
| 10 | Module 2 | Phase 2 | `Phase2_Macro_States.pdf` | PDF | Categorical state composition (the State column: Spatial + Dihedral labels) per condition. | yes |
| 11 | Module 2 | Phase 2 | `Phase2_Pairwise_Categorical_Stats.csv` | CSV | Pairwise categorical tests (state-label distributions) between conditions. | yes |
| 12 | Module 2 | Phase 3 | `Phase3_2D_PhaseSpace.pdf` | PDF | 2D conformational phase space: DFG dihedral (Phi/Psi) and Dunbrack spatial (D1/D2) density panels, coloured by condition. | yes |
| 13 | Module 2 | Phase 4 | `Phase4_Allosteric_Distances.pdf` | PDF | 1D distributions (violins) of every allosteric-network distance metric across conditions. | yes |
| 14 | Module 2 | Phase 4 | `Phase4_Kruskal_Summary.csv` | CSV | Kruskal-Wallis omnibus test per distance metric across conditions. | yes |
| 15 | Module 2 | Phase 4 | `Phase4_Pairwise_Summary.csv` | CSV | Post-hoc pairwise comparisons (with effect sizes) per metric between conditions. | yes |
| 16 | Module 2 | Phase 5 | `Phase5_Correlation_Heatmaps.pdf` | PDF | Per-condition metric-metric correlation heatmaps (the coupling networks). | yes |
| 17 | Module 2 | Phase 5 | `Phase5_Correlation_Shift_Volcanos.pdf` | PDF | Volcano plots of correlation-edge shifts between conditions (which couplings strengthen/weaken). | yes |
| 18 | Module 2 | Phase 5 | `Phase5_Differential_Correlations.csv` | CSV | Per-edge differential-correlation table between conditions (values behind the shift volcanos). | yes |
| 19 | Module 2 | Phase 5 | `Phase5_Global_Network_Density.csv` | CSV | Global coupling score (MAC = mean absolute correlation-edge weight) per condition. | yes |
| 20 | Module 2 | Phase 5 | `Phase5_Global_Network_Density.pdf` | PDF | Plot of the per-condition global coupling score (network density). | yes |
| 21 | Module 2 | Phase 5 | `Phase5_Global_Network_Density_Stats.csv` | CSV | Pairwise statistics comparing global coupling scores between conditions. | yes |
| 22 | Module 2 | Phase 5 | `Phase5_PCA_Biplot.pdf` | PDF | PCA biplot of conditions in the allosteric-metric space (loadings + scores). | yes |
| 23 | Module 2 | Phase 5 | `Phase5_Interactive_3D_Condition_Space.html` | HTML | Interactive 3D (plotly) view of conditions in metric space. Companion assets live in Phase5_Interactive_3D_Condition_Space_files/. | yes |
| 24 | Module 2 | Phase 5 | `Phase5_Interactive_3D_Condition_Space_files/` | dir (assets) | Supporting JS/CSS libraries for the Phase 5 interactive HTML (plotly, htmlwidgets, jquery). Ship alongside the .html. | yes |
| 25 | Module 2 | Phase 6 | `Phase6_State_Assignments.csv` | CSV | Unsupervised (GMM/KMEANS) state-cluster assignment per model (State 1..N). | yes |
| 26 | Module 2 | Phase 6 | `Phase6_State_Clusters_PCA.pdf` | PDF | PCA scatter of models coloured by discovered state cluster. | yes |
| 27 | Module 2 | Phase 6 | `Phase6_State_Composition.pdf` | PDF | Composition of each discovered state by condition (how conditions populate the states). | yes |
| 28 | Module 2 | Phase 6 | `Phase6_Interactive_3D_State_Space.html` | HTML | Interactive 3D (plotly) view of models coloured by state. Assets in Phase6_Interactive_3D_State_Space_files/. | yes |

| | | | | | | |
|---|---|---|---|---|---|---|
| 29 | Module 2 | Phase 6 | `Phase6_Interactive_3D_State_Space_files/` | dir (assets) | Supporting JS/CSS libraries for the Phase 6 interactive HTML. | yes |
| 30 | Module 2 | Phase 6b | `Phase6b_State_Intrinsic_Rigidity_MAC.csv` | CSV | Per-state intrinsic mechanical rigidity: within-state MAC (mean absolute pairwise coupling); higher = more rigid/coupled. | yes |
| 31 | Module 2 | Phase 6b | `Phase6b_State_Intrinsic_Rigidity.pdf` | PDF | Plot of per-state intrinsic rigidity (MAC). | yes |
| 32 | Module 2 | Phase 6b | `Phase6b_State_Intrinsic_Rigidity_Stats.csv` | CSV | Pairwise Wilcoxon statistics on per-state intrinsic MAC (source of manuscript Supplementary Table 1). | yes |
| 33 | Module 2 | Phase 7 | `Phase7_Complete_Structural_Metadata.csv` | CSV | Full merged per-model metadata: master metrics plus derived condition/state columns (analysis-ready table). | yes |
| 34 | Module 2 | Phase 7 | `Phase7_MacroState_Signatures.pdf` | PDF | Biological-signature profiles mapping discovered states to structural features. | yes |
| 35 | Module 2 | Phase 7 | `Phase7_MacroState_Signatures_Stats.csv` | CSV | Statistics behind the metastable-state signature profiles. | yes |
| 36 | Module 2 | Phase 8 | `Phase8_Volcanos/` | dir | Differential allosteric-driver analysis: one volcano per ordered pair of discovered states. For N states = C(N,2) pairs; here 8 populated states -> 28 pairs. Also holds the condition-mode outputs (see below) when Phase 8 is run per condition. | yes |
| 37 | Module 2 | Phase 8 | `Phase8_Volcanos/Stats_State_{i}_vs_State_{j}.csv` | CSV | Per-metric differential statistics (signed effect size, -log10 p) for state i vs state j. | (pattern) |
| 38 | Module 2 | Phase 8 | `Phase8_Volcanos/Volcano_State_{i}_vs_State_{j}.pdf` | PDF | Volcano plot of the metrics that most distinguish state i from state j. | (pattern) |
| 39 | Module 2 | Phase 8 | `Phase8_Volcanos/Stats_{condA}_vs_{condB}.csv` | CSV | Per-metric differential statistics for condition A vs condition B, written when Phase 8 is run in condition mode (as opposed to state mode). Condition tokens are the full system labels, e.g. csk-wtcat-apo_src-wtcat-apo__CSK___Apo_. Additive: the state-pair files above are not replaced. | (pattern) |
| 40 | Module 2 | Phase 8 | `Phase8_Volcanos/Volcano_{condA}_vs_{condB}.pdf` | PDF | Volcano plot of the metrics that most distinguish condition A from condition B (condition mode). | (pattern) |
