## Supplementary material for "Trans-Allosteric Activation Releases Distinct Conformational Traps in Kinase Heterodimers": S3 Data: proteins.yaml

```
metadata:
  generated: '2026-07-14T16:17:02.509233'
  tool: generate_config.py
pattern_matches:
  a-cdk1:
    proteins:
    - name: cdk1
  a-cdk1_b-ccnb1-166:
    proteins:
    - name: cdk1
    - name: ccnb1-166
  a-cdk1-pt14-py15-pt161_b-ccnb1-166:
    proteins:
    - name: cdk1
    - name: ccnb1-166
  a-cdk1-pt161_b-ccnb1-166:
    proteins:
    - name: cdk1
    - name: ccnb1-166
```
