## Supplementary material for "Trans-Allosteric Activation Releases Distinct Conformational Traps in Kinase Heterodimers": S3 Data: proteins.yaml

```
metadata:
  generated: '2026-07-18T20:27:24.268568'
  tool: generate_config.py
pattern_matches:
  a-csk-wtcat-apo_b-src-wtcat-apo:
    proteins:
    - name: csk-wt
    - name: src-wt
  a-csk-wtcat-apo_b-src-wtcat-holo:
    proteins:
    - name: csk-wt
    - name: src-wt
  a-csk-wtcat-apo_b-src-wtcat-py159-apo:
    proteins:
    - name: csk-wt
    - name: src-wt
  a-csk-wtcat-apo_b-src-wtcat-py159-holo:
    proteins:
    - name: csk-wt
    - name: src-wt
  a-csk-wtcat-holo_b-src-wtcat-apo:
    proteins:
    - name: csk-wt
    - name: src-wt
  a-csk-wtcat-holo_b-src-wtcat-holo:
    proteins:
    - name: csk-wt
    - name: src-wt
  a-csk-wtcat-holo_b-src-wtcat-py159-apo:
    proteins:
    - name: csk-wt
    - name: src-wt
  a-csk-wtcat-holo_b-src-wtcat-py159-holo:
    proteins:
    - name: csk-wt
    - name: src-wt
```
