## Supplementary material for "Trans-Allosteric Activation Releases Distinct Conformational Traps in Kinase Heterodimers": S3 Data: README.md

# Underlying data for the trans-allosteric kinase heterodimer study

These are the AlloQuant outputs from which every number reported in the manuscript, its
supplementary notes, its tables and its figures is computed. Distributed with the paper as
**S3 Dataset**, and archived with this repository.

**What is here.** A file is included if a number reported in the paper traces to it, or if a
script shipped in this repository reads it. Excluded: intermediate and temporary files
(`temp_*`, `chunk_*`, `generated_matrix.csv`), superseded v7r2 outputs, plot images, and
internal control runs whose results are not reported as manuscript values.

**How to read the columns.** Every column of the `master_kinase_analysis_results_v7r3.csv`
files is defined in the AlloQuant master CSV data dictionary (**S1 Dataset**; also
`docs/AlloQuant_master_CSV_data_dictionary_v7r3.*`). Every output file produced by the
pipeline, including the ones here, is described in the AlloQuant output file manifest
(**S2 Dataset**; also `docs/AlloQuant_output_file_manifest_v7r3.*`).

## Contents

| Directory | Ensemble | Supports |
|---|---|---|
| `cdk1_ccnb1_260517/` | CDK1–CCNB1, 100 models per condition (600 total) | Fig 2; the quantities annotated on Fig 3; S2 and S9 Tables; Notes 2 to 5 and 15 |
| `csk_src_dimer_260718/` | CSK–SRC heterodimer, 7 conditions (795 paired models) | Figs 5 and 6; S3, S4 and S7 Figs; S3, S4, S7 and S8 Tables; Notes 6 to 14 |
| `csk_monomer_260806_fullmsa/` | Monomeric CSK apo and holo, full-MSA, 45 seeds pooled (n = 225 per condition) | The counts tabulated in S2 Fig; Results on nucleotide-locked monomeric CSK |
| `interface_analyses/` | Derived interface and composition analyses of the dimer ensemble | Fig 5A; Notes 8 and 10 |
| `validation_3d7t/` | AlloQuant applied to the CSK–SRC crystal structure 3D7T, with the Kincore reference output | S1 Table; Note 1 |

Figures that are structure renderings (Figs 1, 3, 4; S1, S2 and S5 Figs) are reproducible from
the model coordinates rather than from these tables. The same applies to the per-residue Cα RMSF
colouring of S5 Fig panel A, which is produced inside UCSF ChimeraX. The predicted coordinates
themselves are not deposited, running to several gigabytes. The `inputs/` folders record what is
needed to regenerate them with AlphaFold3.

Within each ensemble directory:

- `inputs/` holds the AlphaFold3 inputs that define the ensemble: `experiment.csv`, the
  experimental design matrix listing one row per condition, together with `sequences.fasta` and
  `proteins.yaml`, the construct sequences and their per-chain configuration. The monomer
  ensemble has no design matrix because it has no pairing to specify.
- `master_kinase_analysis_results_v7r3.csv` is the Module 1 output, one row per model chain,
  carrying every geometric measurement and categorical state call. The monomer ensemble splits
  this across `_40seeds` and `_5seeds` files, which pool to the 45 seeds and n = 225 per
  condition reported in the paper.
- `plots_and_stats_*_GMM/Phase0` to `Phase7` are the Module 2 outputs: AF3 quality summary,
  ligand-state QC review, categorical statistics, per-metric group tests, global network
  rigidity (MAC) and its differential correlations, unsupervised state assignments,
  per-state intrinsic rigidity, and the complete per-chain structural metadata with state
  labels.
- `plots_and_stats_*_GMM/Phase8_Volcanos/Stats_*.csv` are the post-hoc pairwise driver
  analyses, one file per contrast, giving effect size and adjusted significance per metric.
- `addons/coupling_out_v7r3/` holds the paired cross-chain coupling results (Fig 6).
- `addons/mac_confidence_out_v7r3/` holds the prediction-confidence control, in which MAC is
  recomputed with AlphaFold3 confidence partialled out (S6 Fig; S5 and S6 Tables; Note 13).
- `addons/chain_local_rmsf_out/` holds the chain-local Cα RMSF summary reported as S9 Table,
  and the full per-residue profiles behind it (Note 15). It carries no version tag because
  `addons/chain_local_rmsf.py` reads the model coordinates directly and so does not depend on
  the AlloQuant metric version.

## Two things a reanalyst should know

**Join on identity, not on row order.** Row order in the master CSVs is not stable across
runs. Align records on `Simulation_ID` together with `Chain`.

**Group the CSK–SRC conditions by `Condition_reviewed`, never by directory name.** AlphaFold3
frequently overrides the designed ligand placement, so a model's directory records what was
requested, not what was built. `Condition_designed` cannot detect this; the Truth Overwrite
review in `Phase1_Ligand_QC_Review.csv` resolves it, and `Condition_reviewed` carries the
corrected assignment. This is why the CSK-ATP condition has n = 196 rather than 100, and why
one condition retains only 33 models.

## Not included here

AlphaFold3 model coordinates (2,200 mmCIF files) and the per-model confidence JSONs are too
large for journal supporting information and are deposited separately; see the Data
Availability statement in the paper for the archive DOI.
